# Guided-topic modelling of single-cell transcriptomes enables sub-cell-type and disease-subtype deconvolution of bulk transcriptomes

**DOI:** 10.1101/2022.12.22.521640

**Authors:** Lakshmipuram Seshadri Swapna, Michael Huang, Yue Li

**Affiliations:** School of Computer Science, McGill University, Montreal, QC, Canada

**Keywords:** Cell-type deconvolution, single-cell transcriptome, topic models, variational inference, Bayesian modeling, Type 2 Diabetes, cancer transcriptome, disease biomarkers.

## Abstract

Cell-type composition is an important indicator of health. We present Guided Topic Model for deconvolution (GTM-decon) to automatically infer cell-type-specific gene topic distributions from single-cell RNA-seq data for deconvolving bulk transcriptomes. GTM-decon performs competitively on deconvolving simulated and real bulk data compared with the state-of-the-art methods. Moreover, as demonstrated in deconvolving disease transcriptomes, GTM-decon can infer multiple cell-type-specific gene topic distributions per cell type, which captures sub-cell-type variations. GTM-decon can also use phenotype labels from single-cell or bulk data as a guide to infer phenotype-specific gene distributions. In a nested-guided design, GTM-decon identified cell-type-specific differentially expressed genes from bulk breast cancer transcriptomes.

## Background

Cell-type composition and its relative proportions in a tissue is an indicator of health. For example, several studies have shown that Type 2 Diabetes is characterized by reduced beta cell mass and number in pancreatic tissue [1, 2]. In acute myeloid leukemia (AML), cell-type abundance variation between patients was indicative of degree of malignancy [3]. Experimental approaches such as fluorescence-activated cell sorting (FACS) and immunohistochemistry (IHC) are used to elucidate cell-type composition of biological samples. Single-cell RNA-sequencing (scRNA-seq) technology enables high-resolution cell-type-specific (CTS) transcriptome analysis, providing molecular insights into the cell-type composition, cell-state behaviour, and cell-type heterogeneity [4–7]. In the context of cancer research, scRNA-seq has led to identification of distinct cancer cell states. These are shown to occur across a range of cancer types, and are implicated in tumor progression, with high cell-type heterogeneity associated with poor prognostic outcomes [8, 9]. However, challenges such as high cost, lower throughput, and difficulties in dissociation of cell types in solid samples, make it hard to apply these experimental approaches at the patient population scale.

On the other hand, bulk RNA-seq has been the workhorse behind transcriptome research over the past decades. Its falling costs and ease of experimental setup make it an attractive tool to work with any organism [10]. Several databases host tremendous amounts of bulk RNA-seq data such as Gene Expression Omnibus (GEO) [11, 12], Genotype-Tissue Expression (GTEx) [13], The Cancer Genome Atlas (TCGA), a repository of bulk RNA-seq data for more than 11,000 primary cancer samples [14]. However, bulk RNA-seq data are mixture of gene expression profiles in the tissue. Computational approaches have been developed to deconvolve these bulk RNA-seq profiles into their constituent cell types in the form of cell-type proportions, since these are substantially cheaper and easier to obtain than conducting scRNA-seq experiments. Moreover, deconvolving the bulk data into constituent cell-type components can not only yield the cell-type proportions facilitating clinical investigation but also enable high-resolution differential analysis of gene expression. Studying the differentially expressed genes at the cell-type or cell-state level can help uncover gene regulatory programs that drive different tissue states. Many deconvolution methods were developed to this end.

Most of the deconvolution approaches require a set of gene markers for each cell type of interest. These marker genes are derived from expert knowledge or differential expression analysis of purified samples of specific cell types. Early methods that leverage these marker genes could achieve good performance in deconvolving mixtures with highly distinct cell types such as blood [15]. CIBERSORT improved on these approaches by incorporating a feature selection step, where genes are adaptively selected from the signature matrix based on the input bulk RNA-seq data [16]. It uses a linear support vector regression (SVR) to delineate closely related cell-types such as leukocytes. BSEQ-sc combined with CIBERSORT was an early method that used scRNA-seq data as a reference for bulk deconvolution [17]. CIBERSORTx also uses scRNA-seq as reference profiles, along with improved normalization schemes to suppress cross-platform variation, and an adaptive noise filter to eliminate unreliably estimated genes [18]. MuSiC adopts a weighted non-negative least-squares regression approach and addresses the issue of cross-subject heterogeneity as well as within-cell-type variation of gene expression [19]. EPIC accommodates user-defined reference profiles to account for the presence of uncharacterized cell types in the target bulk samples [20]. Bisque learns gene-specific transformations of the bulk data based on the single-cell reference profiles and the corresponding cell proportions to account for their differences [21]. BayesPrism uses Bayesian inference to model scRNA-seq data jointly with bulk RNA-seq data to infer cell-type composition and their proportions [22]. The joint modeling overcomes biases that may arise due to technical and biological differences. Beyond cell-type deconvolution, some recently developed methods can estimate CTS gene expression from the bulk samples [23, 24]. However, these methods rely on an external cell-type deconvolution method like those aforementioned ones and do not utilize or model the distribution of the scRNA-seq reference data to properly express statistical uncertainty while leveraging their information richness.

In this study, we present a guided topic model for cell-type deconvolution (GTM-decon). As an overview of our analysis, we first benchmark GTM-decon on deconvolving simulated and real bulk data in comparison with the state-of-the-art deconvolution methods. We then train GTM-decon on pancreatic and breast scRNA-seq datasets to deconvolve bulk RNA-seq datasets from pancreatic and breast tissues, respectively. When applied to deconvolving cancer bulk transcriptomes, GTM-decon successfully identifies the cell type of origin for pancreatic and breast cancer datasets. Interestingly, the results for human pancreatic cancer are recapitulated using the CTS topics inferred from the mouse pancreas scRNA-seq data, postulating cross-species deconvolution as an option where it is difficult to obtain scRNA-seq data due to technical or ethical challenges. Furthermore, our GTM framework also enables the inference of phenotype-specific topic distributions from bulk RNA-seq data by using the phenotypes (e.g., breast cancer subtypes) from single cell or bulk RNA-seq data as a guide for the topic inference. We leverage this capability to distinguish Basal from Estrogen receptor (ER) positive (ER+) breast tumor samples, not only achieving high classification accuracy but also identifying the genes and pathways that segregate the cancer subtypes. By fine-tuning the inferred CTS topic distributions guided by the breast cancer subtypes, we deconvolve the average differential gene expression into CTS expression changes, thereby enabling discovery of the subtype-specific aberrance of the gene regulatory programs.

## Results

### GTM-decon model overview

In GTM-decon, we have made <u>three methodological contributions</u>. As our <u>first and the main</u> <u>contribution</u>, GTM-decon is a marker-free method and automatically infers the contribution of each gene for each cell type in the form of CTS categorical distributions, which we define as “topics” [25], without using marker gene information. Each CTS topic distribution is related to the transcriptional rate of each gene for each cell type. For instance, B cells have higher transcription rate for *CD19* compared to alpha cells, which have relatively high rate for *FXYD5*. Conceptually, we consider genes as vocabulary and cells as documents whose word tokens (i.e., scRNA-seq reads) are sampled from the vocabulary with the CTS topic probabilities. We incorporate the observed cell-type labels for each cell in the form of topic prior to guide the inference of CTS topic mixture, which reflects the uncertainty of the noisy cell-type label. Specifically, the cell-type mixture for cell *m* follows a K-dimensional asymmetric Dirichlet distribution, **θ**_*m*_∼*Dir*(**α**_*m*_ + 0.1), with the hyperparameter *α*_*m*,*k*_ set to a relatively high value (i.e., 0.9 by default) given the cell-type label *y*_*m*_ = *k*; the rest of the K-1 α_*m*,k′_ values, where *y*_*m*_ ≠ *k*^′^, are set to a relatively low values (i.e., randomly sampled from a range between 0.1 and 0.01). As a result, each topic is automatically identified with exactly one cell type. This differs from the standard topic model, where topics are not directly associated with any known concept and require *post hoc* manual inspection based on their top scoring words to interpret them. Given the CTS gene distributions, we can infer the CTS topic mixtures from the bulk transcriptomes, which are the desired cell-type mixing proportion in the context of deconvolution (**Fig. 1a**).

**Figure 1.**
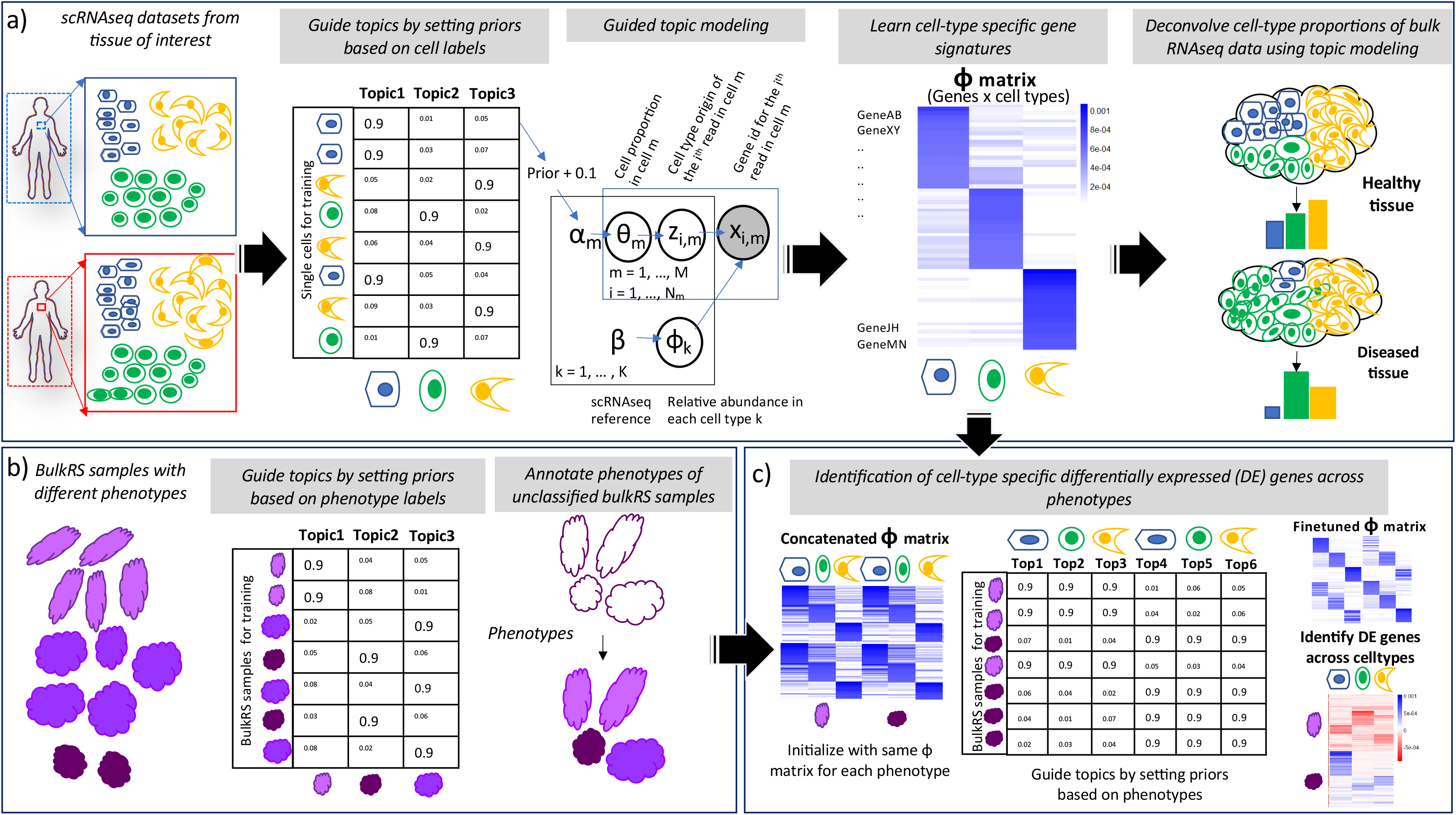
GTM-decon overview. **a**) Inferring cell-type-specific (CTS) topics from scRNA-seq reference data. In brief, GTM-decon infers CTS topics from scRNA-seq data by using a guided topic modeling approach utilizing cell-type labels from the reference. High prior values are assigned to the topic corresponding to the cell type, and lower prior values are assigned to the other topics, enabling it to learn a genes-by-CTS topics matrix, with each topic anchored to a specific cell type. This matrix is used to infer cell-type proportions in bulk RNA-seq data using standard topic modeling, capturing variations in cell type proportions between healthy and diseased tissue. The probabilistic graphical model (PGM) diagram depicts the data generative process assumed by the proposed guided topic model. Suppose there are K cell types in the scRNA-seq data. For each cell indexed by *m* ∈ {1, …, *M*}, we use K-dimensional Dirichlet-distributed cell-type topic mixture **θ**_*m*_∼Dir(**α**_*m*_) to represent the statistical uncertainty of the noisy cell-type label *y*_*m*_ ∈ {1, …, *K*}. Specifically, we clamp the Dirichlet hyperparameter *α*_*m*,*ym*_ of the Dirichlet variable to a relatively high value while setting the rest of the values of *α*_*m*,*k′*_ (*y*_*m*_ ≠ *k*^′^) to relatively low values (i.e., 0.9 and [0.01, 0.1], respectively in the cartoon illustration of M=8 cells and K=3 cell types). The non-zero prior values for the K – 1 unobserved cell types allow the cell-type mixture variable **θ**_*m*_ to have non-zero density over those cell types as dictated by the scRNA-seq data likelihood and therefore account for potentially mislabelled cell types. Suppose there are in total *N*_*m*_ reads in cell *m*. Each scRNA-seq read *i* ∈ {1, …, *N*_*m*_} is assumed to be originated from one of the K CTS topics with the Categorical rates fixed to the cell-type mixture, i.e., *z*_*i*,*m*_ ∼ Cat(**θ**_*m*_). Given cell-type topic assignment *z*_*i*,*m*_ ∈ {1, …, K}, the *i^th^* read is then mapped to one of the G genes as indexed by *x*_*i*,*m*_ with Categorical rates set to be **ϕ**_*zi*,*m*_, which itself is a G-dimensional Dirichlet variable of flat hyperparameter *β*, i.e., *x*_*i*,*m*_∼Cat(**ϕ**_*zi*,*m*_). To infer the latent variables, namely cell-type mixture proportion **θ**_*m*_∼Dir(α), CTS topic assignments for each read *z*_*i*,*m*_, and CTS topic distributions **Φ**, we employ an efficient collapsed variational Bayes algorithm as detailed in the **Methods** section. The genes-by-CTS-topics **Φ̂** matrix estimated from the scRNA-seq reference then serves as a template when it comes to infer the cell-type mixing proportions θ_*j*_ of a bulk RNA-seq sample *j* using essentially the same inference algorithm as in the scRNA-seq data modeling except for having a flat hyperparameter for the prior (e.g., α_*k*_ = 1∀*k* by default) while fixing **Φ̂** and only inferring the expected total reads allocated for each CTS topics (i.e., *E*_*q*_[*n*_.,*j*,*k*_] = *E*_*q*_[∑_*i*_[*z*_*i*,*j*_ = *k*]]). **b**) Phenotype-guided modeling of bulk RNA-seq data. GTM-decon can also use phenotype labels as a guide for topic inference to model sparsified bulk transcriptomes in a disease study. In this design, instead of having each row as a cell and each column as a cell type, each row corresponds to a bulk sample and each column to a phenotype class. For each subject *j*, we set the topic hyperparameter *α*_*j*,*yj*_ based on the noisy phenotype label *y*_*j*_ of the subject. The inference algorithm is the same as in modeling the scRNA-seq reference data. Given a test subject *j*^′^, the inferred topic mixture **θ**_*j′*_ represents the phenotypic probabilities of the subject. **c**) Nested-guided topic model for detecting cell-type-specific differentially expressed genes between phenotypes. In this nested design, we treat the phenotype as level 1 and the cell types as level 2. The pre-trained genes-by-CTS-topics distribution **Φ̂** learned from panel **a**) are used to initialize the topic distributions for each phenotype in a sparsified bulk transcriptome disease study. As illustrated in the cartoon, for example, for 2 phenotypes and 3 cell types, there are 6 topics. GTM-decon then fine-tunes the combined cell-type-specific topic distribution by running the same algorithm described in panel **b**). The resulting topic distributions reflect the phenotypic influences on CTS gene distributions, which are the statistics for conducting differential expression analysis in a case-control study design.

As our <u>second contribution</u>, we extend GTM-decon to infer multiple topics per cell type. The rationale is that cells of the same cell type can manifest in different cell states due to the changes of environments or stimuli. As a result, these cells may exhibit expression patterns that are different from the canonical CTS expression pattern. While there are sophisticated hierarchical topic models involving Dirichlet Processes [26], we took a simple and elegant design. Specifically, we extend the basic GTM-decon model to infer sub-cell-type topics by dedicating multiple topics per cell type (**Additional file 1**: **Figure S1**). As our <u>third contribution</u>, we extend GTM-decon to infer phenotype-specific (PTS) topic distributions using the phenotype label (e.g., cancer subtypes or cancer stages) available in the single-cell or bulk transcriptome data as a guide to detect PTS gene expression (**Fig. 1b**). We then further extend it to a nested-guided topic model to conduct CTS differential expression analysis in the single-cell or bulk patient cohort data (**Fig. 1c**). To that end, we use the phenotype labels as the level-1 guide and the cell-type labels as the level 2 guide. Through the same guided topic mechanism, GTM-decon updates the CTS topic distributions under each phenotype by fitting the data likelihood of the transcriptomes from either the single-cell or bulk data. The algorithmic details for the 3 contributions were described in **Methods**.

### Experimentation of data preprocessing and GTM-decon model configurations

We experimented gene selection, data normalization, hyperparameter settings, and number of topics per cell type. We find that GTM-decon works the best with raw read count data using all genes (**Additional file 1**: **Figure S2** and **S3)**, and it is fairly robust to different hyperparameter values for the topic mixture prior (**Figure S4**-**S6**) and the CTS topic prior values (**Figure S7)**. In general, GTM-decon confers better deconvolution performance using multiple topics per cell type than the baseline GTM-decon with one topic per cell type (**Figure S8**). Please refer to **Additional file 1 Section S1-S5** for more details.

### Evaluation of deconvolution of simulated bulk from scRNA-seq data

To quantitatively benchmark GTM-decon, we compared it against five existing deconvolution methods, namely Bisque [21], Bseq-sc [17], CIBERSORTx [18],MuSiC [19], and BayesPrism [22] on artificially simulated bulk RNA-seq datasets from the scRNA-seq data (**Additional file 1: Table S1**). To simulate bulk data, three human scRNA-seq datasets (Pancreas – E-MTAB-5061, Breast Tissue with GEO accession number GSE113197, and Rheumatoid Arthritis (RA) Synovium – SDY998), generated using different technologies (Smart-seq2, 10x Genomics Chromium, CEL-Seq2) were used. The artificial bulk data for each individual was constructed by summing up the counts for each gene from all cells in that individual [19]. This allowed us to use the cell-type proportions from the single-cell data as the ground-truth proportions. Artificially constructed bulk data from scRNA-seq data appear to be a good surrogate of the real bulk data, as observed from the excellent correlation of the log-transformed artificial counts with the log-transformed counts from real bulk data for each gene (**Additional file 1**: **Figure S9**). We performed leave-one-out cross validation (LOOCV), to avoid any leakage from training data, and used the single-cell RNA-seq of the left-out individual to simulate the bulk RNA-seq (i.e., the total read counts of each gene for that sample) as the validation data and its cell-type proportions as the ground-truth mixing proportions. GTM-decon performs better than other models on the Pancreas and Breast Tissue datasets in terms of both Spearman Rank-based correlation (Spearman R) and Cross Entropy and conferred comparable performance on the RA Synovium dataset (**Additional file 1**: **Figure S10**). Notably, GTM-decon achieves smaller variance for both the pancreatic and breast tissue datasets. We further ascertained the qualities of the predicted cell-type proportions of each method against the ground-truth cell-type proportions (**Figure S10a**) and observed that GTM-decon recapitulates cell-type proportions well for pancreatic (**Figure S10b**), breast tissues (**Figure S10c**), and RA synovium dataset (**Figure S10d**).

### Evaluation of deconvolution of real bulk with ground-truth cell-type proportions

We also benchmarked GTM-decon on 5 real bulk RNA-seq data with known ground-truth cell-type proportions from 3 different tissue types (**Additional file 1**: **Table S1** and **S2**; **Methods**). We evaluated deconvolution performance using Spearman R and Root Mean Square Error (RMSE), by comparing the inferred proportions for the matching cell types against the ground-truth proportions for each sample. GTM-decon conferred on-par or superior performance compared to the existing methods for all the datasets (**Fig. 2a**). In particular, GTM-ALL performed the best for deconvolving PBMC-1 and PBMC-2. For deconvolving Whole Blood (WB), GTM-HVG performed the best in terms of both metrics and MuSiC is a close second. While deconvolving bulk RNA-seq from the prefrontal brain region from ROSMAP dataset, all methods except Bseq-sc and GTM-HVG performed reasonably well. It is possible that the HVG are the genes having high variance within the cell types, which caused the poor performance of GTM-HVG on this dataset. For the pancreatic dataset with the paired single-cell and bulk RNA-seq data collected from the same individuals, GTM-HVG performs the best with GTM-PP as the runner-up in terms of Spearman R, and RMSE are similar among all methods except MuSiC with notably higher error. In summary, GTM-ALL performed the best in 3 datasets; GTM-HVG performed the best in the other two datasets, where the CTS gene expression might exhibit more distinct inter-cell-type variability. Furthermore, we also evaluated the deconvolution performance for separate cell-types in terms of the correlation between ground-truth proportions and predicted proportions for each cell type across samples. Overall, GTM-decon conferred competitive performance (i.e., the right red bar plots in Figure R9b) with the runner up method being BayesPrism (**Fig. 2b**; **Additional File 1: Figure S11-S15)**. Therefore, the results suggest the general effectiveness of topic modeling in cell-type deconvolution and the additional benefits conferred by GTM-decon due to its algorithmic innovations.

**Figure 2.**
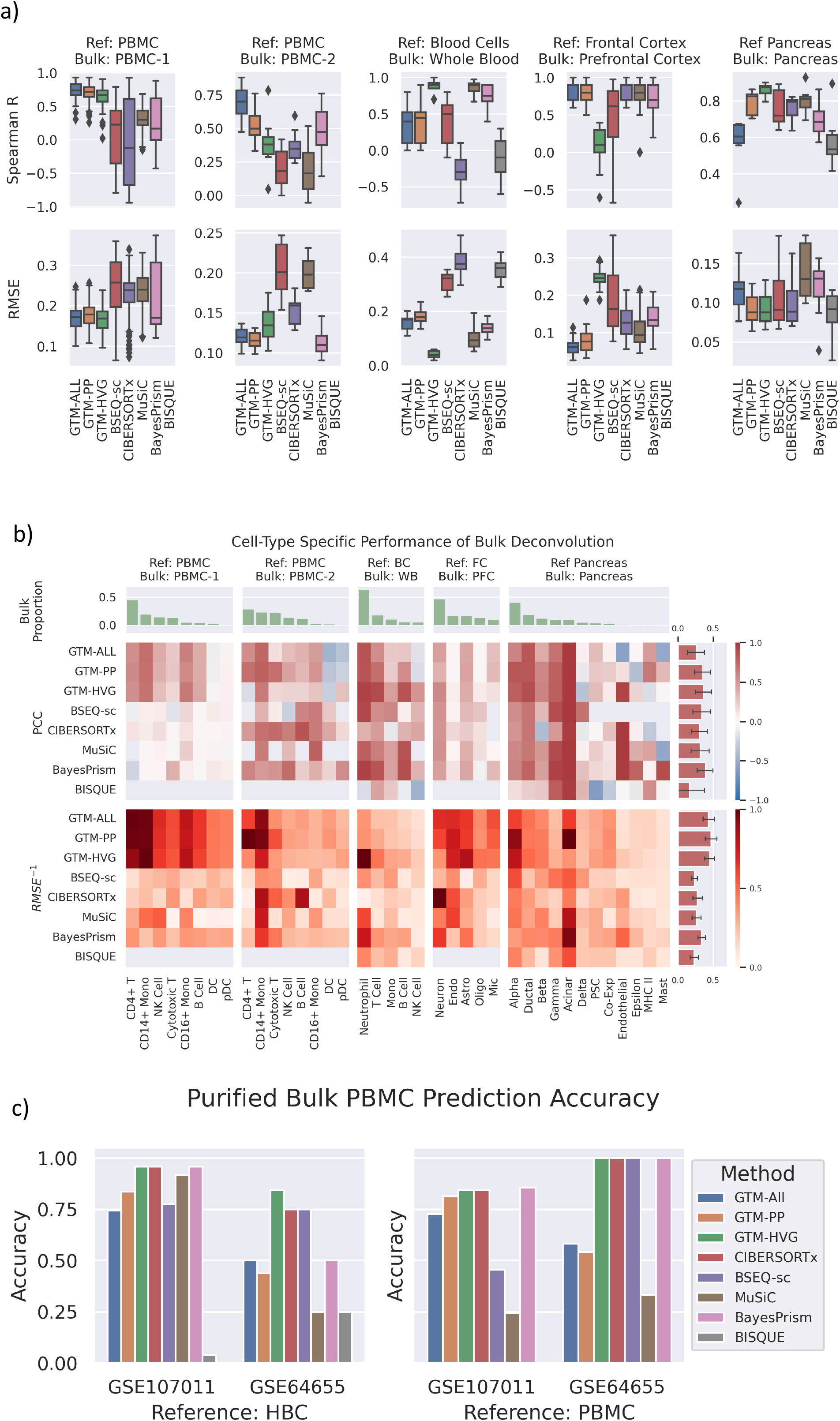
Evaluation of deconvolution performance on real bulk data. **a**) Evaluation of sample deconvolution. We evaluated the deconvolution performance of GTM-decon using all genes (GTM-ALL), preprocessed genes (GTM-PP), and highly variable genes (GTM-HVG) with five SOTA methods – CIBERSORTx, MUSIC, BSEQ-sc, BISQUE, BayesPrism. The 3 immune bulk data and the brain data were deconvolved using independent references of a similar tissue, while the pancreas bulk data is deconvolved using single cell reference from the same individuals in a leave-one-out cross-validation (LOOCV) manner. The bulk labelled PBMC-1 corresponds to SDY67 dataset, PBMC-2 corresponds to S13 cohort, Whole Blood to Whole Blood dataset, and Prefrontal cortex to ROSMAP dataset (**Additional file 1**: **Table S2**). For each test bulk sample, Spearman Correlation and Root Mean Square Error (RMSE) were computed between its ground-truth and predicted cell-type proportions by each method. The box and whiskers in each boxplot indicate the 25%-75% quartile and min-max of the evaluation scores over all samples in a dataset, respectively. **b)** Heatmaps comparing the cell-type specific deconvolution performance of GTM-decon against existing methods on 5 different real bulk datasets with known ground truth mixing proportions. The cell-types are ordered from most to least prevalent in the bulk data (green barplots in first row indicate average proportion for each cell-type in the bulk data). The middle row shows the Pearson Correlation Coefficient between the predicted and known cell-type proportions. The lower row shows the Inverse RMSE (higher is better, scaled between 0 - 1), per cell-type per dataset. The barplots on the right shows the average performance over all cell-types for each method. **c)** Cell-type prediction accuracy of the purified immune bulk RNA-seq samples. The two panels indicate the use of different independent immune references, for the deconvolution of two purified bulk immune datasets (Accession Numbers: GSE107011, GSE64655). For each purified bulk sample, the cell type corresponding to the highest inferred cell-type proportion by each method was used as the predicted cell type. The barplots show the prediction accuracy as the percentage of the correctly predicted samples.

Additionally, we evaluated the deconvolution accuracy for purified bulk RNA-seq data of immune cells (GEO accession numbers: GSE107011 and GSE64655; **Table S2**) using two different independent references (HBC and PBMC2; **Table S1**). Overlapping cell types were used to evaluate the performance on the purified bulk samples, whereby the highest deconvolution proportion was used as the predicted cell-type label for computing the prediction accuracy. GTM-HVG achieved the highest accuracy across all four experiments (**Fig. 2c**). Moreover, since some cell types present in the bulk are missing in the scRNA-seq reference, a robust model should either find the closest-matched cell types or properly express statistical uncertainty in this situation. To this end, we examined the inferred cell proportions of the purified bulk for those missing cell types. Plasmablast samples are present among the purified PBMC samples (GSE107011) but absent in the HBC reference (**Additional file 1**: **Figure S16a**). GTM-decon assigned Plasmablast samples with high cell-type proportions for B-cell, a cell type that shares Plasmablast cell lineage. The inferred cell-type proportions for Basophils purified samples were split between HSPCs (immune progenitor cells) and Neutrophils, which is also classified as Granulocyte. Granulocytes are the most common white blood cells, consisting of 3 specific cell types – neutrophils, eosinophils, and basophils. Using PBMC2 as a reference to deconvolve the same purified bulk immune samples led to similar deconvolution patterns for the missing cell types of Plasmablast and Basophils (**Additional file 1**: **Figure S16b**). Interestingly, Neutrophils were absent in the PMBC2 reference and inferred to be monocytes, which are related to the granulocyte family – a class of immune cells that include Basophils, Eosinophils, and Neutrophils [27]. Another missing cell type HSPC (haematopoietic stem and progenitor cells) have their signal spread across all the cell types. These results suggest that when some cell types are missing in the reference, GTM-decon either finds the closest match or appropriately expresses uncertainty.

Finally, we performed benchmarking on the time and memory usage of our GTM-decon software. GTM-decon scales linearly with both the number of topics per cell type and the number of cells (**Additional file 1**: **Section S6**; **Figure S17**), which is what we expected since its time and space complexity are both O(*N* × *G* × *K*) for N cells, G genes, and K topics. For large number of cells, we can also perform stochastic variational inference [28] to rapidly update model parameters based on mini-batches of cells with much lower memory overhead. It also compares favourably with BISQUE, BSEQ-sc, and MuSiC in terms of running time and memory usage.

### GTM-decon automatically learns CTS gene signatures from scRNA-seq reference

We evaluated the performance of GTM-decon in recapitulating cell-type specific information as well as deconvolution using pancreatic tissue as a reference. The pancreas consists of several cell types including exocrine and endocrine. While the former aids digestion by secreting several enzymes, the latter regulates glucose uptake and processing by secreting hormones. With a vast literature documenting the biological roles of several cell types and their behaviour in healthy and diseased conditions, such as diabetes and cancer, the pancreatic tissue serves as a good benchmark to assess GTM-decon. We trained GTM-decon on an scRNA-seq reference dataset of pancreatic tissue from Segerstolpe et. al [29], consisting of 2,209 cells, corresponding to 14 cell types from 10 individuals. GTM-decon captured distinct sets of CTS gene signatures, as shown by the gene-by-topic probability distributions (i.e., the matrix **ϕ**) for the top 20 genes in each topic (**Fig. 3a**). Indeed, each topic recovers a large number of marker genes for the corresponding cell types based on two databases, namely CellMarker database [30], a manually curated resource of cell markers in human and mouse and the PanglaoDB [31], a database of marker genes generated from scRNA-seq datasets (**Fig. 3a**). We further ascertained the cell-type coherence of each topic by Gene Set Enrichment Analysis (GSEA), while using the probabilities learnt for each cell type against the CellMarkerDB. For the three cell types with abundant marker genes – acinar, alpha, and beta, each of the 5 topics recovers the exact cell type as the top-most hit in the analysis, with the adjusted p-value ≤ 1×10^-15^ (permutation test) in most cases (**Fig. 3b**). Furthermore, the enrichment of known marker genes for the main cell types suggested that GTM-decon with 5 topics per cell type best captures the cell-type specific signatures (**Additional file 1**: **Figure S18**). We also evaluated the effect of different number of cells per cell type. As expected, the topic confidence scores as measured by the average probabilities over the CTS gene distributions increase with the increasing number of cells for that cell type (i.e., evidence) (**Additional file 1**: **Figure S19**).

**Figure 3.**
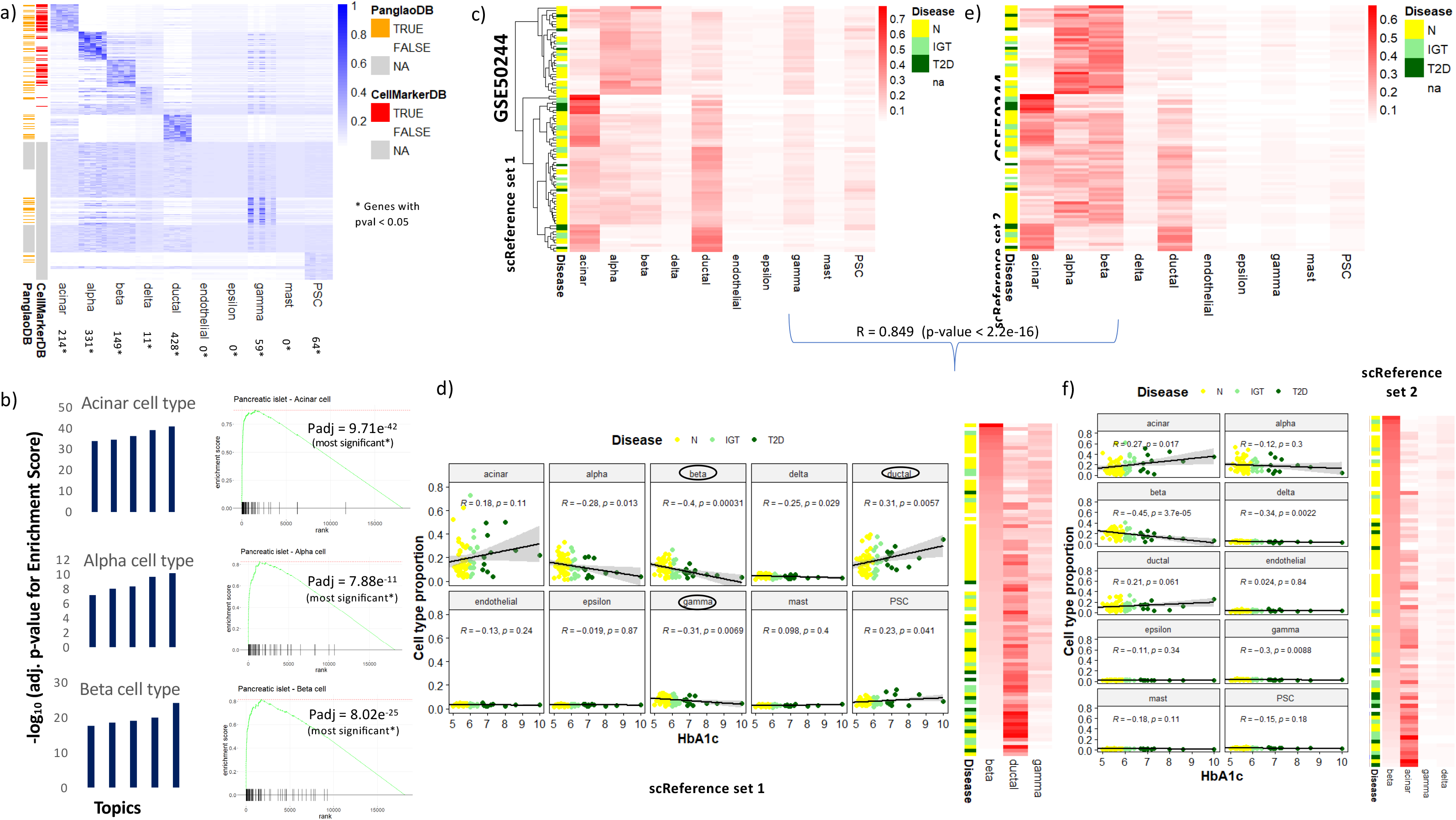
Cell-type-specific topic inference and deconvolution of pancreatic tissue. **a**) Gene signatures of cell-type-specific topics in pancreas. We trained GTM-decon on the PI-Segerstolpe scRNA-seq dataset of pancreas tissue. We used 5 topics per cell type, allowing sub cell-type inference. For the inferred genes-by-cell-types matrix Φ, we took the top 20 genes under each topic and visualized their topic distributions in heatmap. Whenever available from CellMarkerDB and PanglaoDB, cell-type marker genes are indicated on the left. For the cell types where marker genes are not available, “NA” were indicated on the left. The number of statistically significantly different genes in each cell type based on their topic scores (p-value < 0.05; permutation test) is shown below. **b**) Gene set enrichment analysis (GSEA) of inferred topics based on known marker genes. Cell-type-specific topics for Acinar, Alpha, and Beta were evaluated based on whether the top genes are enriched for the known marker genes under that cell type. The bar plots show the -log_10_ (p.adj values of enrichment score) for the gene set enrichment analysis for each of the 5 topics. The leading-edge plot for the topic with the best adj. p-value for that cell type is shown on the right. In each of the leading-edge plots, genes were ordered in decreasing order from left to right. The green curves indicate the running scores of enrichments. The barcode bars indicate cell-type marker genes. Adjusted p-values based on the GSEA enrichment scores are indicated in each panel. The three large panels display the most significantly enriched topic of among the five topics for each cell type and the 12 small panels display the remaining topics. **c**) and **e**) Deconvolution of bulk RNA-seq samples of 89 human pancreatic islet donors. The GTM-decon models separately trained on the Segerstolpe pancreas islet dataset (i.e., panel c) and Baron pancreas islet data (i.e., panel e) reference datasets were used to deconvolve the 89 bulk transcriptomes. As indicated by the legend, the 89 subjects consist of 51 normal, 15 impaired glucose tolerance, and 12 T2D individuals. In the heatmap, the rows represent subjects, and the columns represent cell types; the color intensity are proportional to the inferred cell-type proportions. **d**) and **f**) Deconvolved cell-type proportions as a function of Hemoglobin A1c (HbA1c) level. GTM-decon were trained on Segerstolpe (i.e., panel d) and Baron scRNA-seq (i.e., panel f) reference datasets. Each of the 10 panels displays a scatter plot of inferred cell-type proportion (y-axis) and HbA1c level (x-axis). The color legend indicates the 3 phenotypes. The heatmap on the right shows the deconvolved proportion of 3 most indicative cell types with subjects (rows) ordered on the basis of inferred beta cell-type proportions.

### GTM-decon delineates the variations of cell-type proportions in pancreatic tissues of healthy and T2D subjects

Based on the inferred CTS topic distributions **ϕ**, we used GTM-decon with 5 topics per cell type to infer the cell-type proportions of bulk RNA-seq data from a cohort of 89 human pancreatic islet donors with and without Type 2 Diabetes (GEO accession number: GSE50244) [32] (**Fig. 3c**). The dataset consists of 51 individuals with normal glucose tolerance (N), 15 with impaired glucose tolerance (IGT) and 12 with Type 2 Diabetes (T2D); it also has a good segregation of males (N_1_=54) and females (N_2_=35). As expected, the inferred proportions of most types of cells for these two sets of individuals are similar since they came from the same tissue. However, GTM-decon predicted a significant reduction in beta cells in T2D individuals (Pearson correlation coefficient (PCC) with HbA1c = -0.4, p-value = 0.00031; t-distribution with n-2 degrees of freedom) (**Fig. 3d**). IGT and T2D individuals exhibit low Beta cell-type proportions (**Fig. 3d**), which was supported by the literature [1, 2]. The increase in ductal cells is possibly caused by their regulation of glucose uptake (**Fig. 3d**). These results are consistent when using a different pancreatic dataset as reference (i.e., the PI-Baron reference dataset, generated via Drop-seq instead of PI-Segerstolpe, generated via Smart-seq) (**Fig. 3e**, **3f**).

### Deconvolving human pancreatic data from mouse pancreas scRNA-seq reference

For cases where scRNA-seq data were not available for the organism of interest, due to either ethical, technical, or financial challenges, there is a need to leverage scRNA-seq collected from a model organism. To this end, we investigated the possibility of deconvolving bulk RNA-seq data by training GTM-decon on a mouse scRNA-seq data. We separately trained two GTM-decon models on the human and mouse pancreatic datasets. Specifically, the human datasets include the Segerstolpe and Baron datasets (**Additional file 1**: **Table S1**), which consist of 2,209 cells from 10 individuals and 8,569 cells from 4 individuals, respectively; the mouse dataset consists of 1886 cells collected from 2 mice. For comparative analysis, we focused on only the common set of high-confidence orthologous genes between the two species mapped by the Ensembl database [33]. We visualized the cell-type proportions comparing against the ground-truth values (**Additional file 1**: **Figure S20a**). As expected, the cell-type proportions deconvolved using the two human datasets accurately recapitulate the ground-truth proportions (median PCC of 0.94 and 0.97). Interestingly, GTM-decon trained on the non-reference human dataset performed better than the one trained on the reference-matched dataset, which was probably due to the 10-fold higher number of cells in the former scRNA-seq dataset. Moreover, GTM-decon trained on the mouse reference dataset also performed quite well in terms of the concordant proportions of the shared cell types (PCC of 0.94).

### Deconvolving pancreatic cancer transcriptomes identified tumor cell-type origin

We next turned to deconvolving pancreatic adenocarcinoma (PAAD) (also known as pancreatic ductal adenocarcinoma or PDAC) tumor bulk samples from TCGA. For this application, we also used GTM-decon with 5 topics per cell type. Since the tumor microenvironment is known to be infiltrated with immune cells [34, 35], we sought to train GTM-decon on a single-cell reference dataset derived from individuals with pancreatic cancer, in order to capture the cell types of both the tissue of interest and the immune cells in its tumor microenvironment. To this end, we trained GTM-decon on an scRNA-seq dataset comprised of the transcriptomic profiles of about 57,000 cells from 24 primary PAAD tumors and 11 healthy control pancreas samples [36] in order to deconvolve the 174 bulk RNA-seq profiles from the TCGA PAAD tumor samples. Additionally, we also sought to identify possible novel cell types or pathways present in the bulk RNA-seq, which are not represented in the reference profiles. This is achieved by running an unguided topic model (i.e., a standard LDA) on the sparsified bulk RNA-seq data to detect *de novo* bulk RNA-seq (bulkRS) topics (**Methods**). We empirically chose the number of *de novo* bulkRS topics based on how well they could explain the variation observed in the clinical phenotypes.

We observe that the most prevalent cell types are 4 main pancreatic cell types, namely ductal (type 2), acinar, endocrine (alpha and beta cell types), and fibroblasts (**Fig. 4a**). Notably, the cell type of tumor origin is correctly predicted for the samples: Ductal cells have the highest proportion among the PAAD samples (**Fig. 4b**; brown rectangle), and acinar for a subset of the PAAD samples (**Fig. 4b**; blue rectangle). This recapitulates the literature remarkably well, as ductal cells are known to be the site of the tumor origin for most cases of PAAD; however, a subset arises from acinar cells [37]. More significantly, an unknown subtype is predicted to originate from endocrine cells (**Fig. 4b**; green rectangle). This is supported by the recent literature, which reported these samples as the derivatives of the pancreatic neuroendocrine tumor (PNET) were in fact misclassified as PAAD [38]. PNET are supposed to originate from alpha or beta cells (endocrine cells) [39]. In addition, most of the samples are predicted to have a high proportion of fibroblasts, which are known to be prevalent in pancreatic cancer (**Fig. 4b**; blue dot) [40–42]. Interestingly, deconvolving the pancreatic cancer PAAD dataset using mouse reference dataset also conferred high-quality patient clustering comparable to that of human dataset (**Additional file 1**: **Figure S20b**). Notably, the PAAD “other subtype” was predicted to originate from alpha cells (an endocrine cell), which mirrors the results from human reference data (**Figure S20b**; shown in blue rectangles).

**Figure 4.**
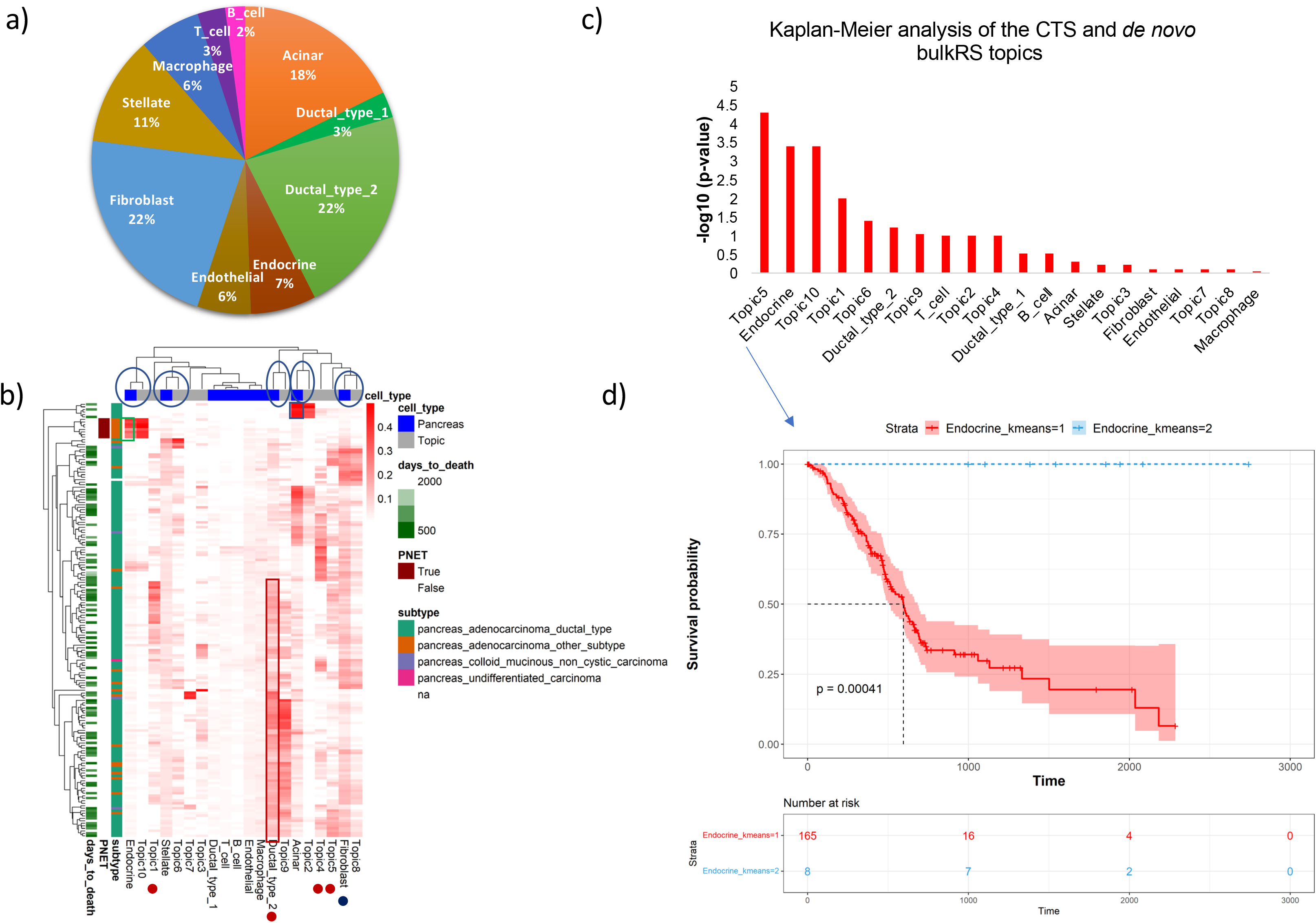
Deconvolution of bulk RNA-seq samples for pancreatic cancer from TCGA-PAAD. GTM-decon was first trained on an scRNA-seq dataset from individuals with pancreatic cancer. The trained GTM-decon model was then used to infer the cell-type proportions of the 174 TCGA-PAAD bulk RNA-seq profiles. **a**) The average inferred cell-type proportion across the TCGA-PAAD tumor samples. We summed up inferred cell-type proportions over all samples followed by normalization. The pie chart displays the resulting percentage of cell-type proportions. **b)** Inferred cell-type proportions of individual TCGA-PAAD tumor samples. To complement the inferred proportions of known cell types, we also ran unguided topic model (i.e., LDA) on the TCGA-PAAD bulk RNA-seq profiles directly to detect *de novo* topics that are not present in scRNA-seq reference. The heatmap visualizes the combined deconvolution results based on the 10 pancreatic cell types, and 10 *de novo* topics (i.e., columns). Each of the 174 rows represents a subject. Three types of demographic or clinical phenotypes were shown in the legend to aid result interpretation. These include “days to death”, cancer subtype, and whether the cancer type is PNET or not. The regions in the highlighted boxes were discussed in more details in the main text. **c**) Survival analysis of the CTS and *de novo* topics using inferred cell-type proportions. The 174 subjects were divided into two groups based on K*-means clustering with K* set to 2 (not to be confused with the K cell types or topics). Kaplan-Meier curves were generated for these groups and compared using log-rank test. The plot shows the –log_10_(p-value) from the log-rank test for all the CTS and *de novo* topics, in decreasing order of significance. **d)** Kaplan-Meier curve for Endocrine cell-type proportion. The curve and shaded area represent the mean and standard deviation of the cell-type proportions in the two groups, respectively. The number of subjects for each cluster were indicated in the bottom panel.

Interestingly, the *de novo* bulkRS topics cluster with the most abundant reference topics inferred from the scRNA-seq reference data (**Fig. 4b**, blue circles). For example, bulkRS Topic 2 corresponds to the CTS topic for Acinar cell type, Topic 10 to the CTS topic for Endocrine cell type, Topic 9 to Ductal_type_2, Topic 8 to Fibroblast, and Topic 6 to stellate cells. However, there are a few *de novo* bulkRS topics, such as bulkRS Topic 1, Topic 4, and Topic 5, capturing distinct distributions for specific subsets of samples (**Fig. 4b**; red dots). These topics could correspond to either novel cell types or gene pathways in the bulk but not implicated in the scRNA-seq reference.

We next estimated whether variation in cell-type proportions or bulkRS topics is indicative of survival time. To this end, we performed Cox Regression to regress the number of days patients lived since their cancer diagnosis on their inferred cell-type proportions as well as the *de novo* bulkRS topics. Overall, the Cox Regression model is statistically significant compared to the bias term (adjusted p-value = 9 × 10^-5^ based on likelihood ratio test). To explore the marginal effect of individual cell-type proportions on survival, we performed Kaplan-Meier analysis by separating patients into two groups based on K-means clustering (**Fig. 4c**). We observe that among the cell types, Endocrine cell type exhibits a significant hazard ratio, predicting a good survival outcome (**Fig. 4d**; p-value = 0.0091; log-rank test), which is supported by the literature as PNETs are mostly benign [43]. However, Ductal cell type 2 is associated with poor survival outcome, which is expected as pancreatic adeno carcinoma are aggressive (p-value = 0.03; log-rank test). On the other hand, Topic 1 and Topic 5 indicate poorer survival, with hazard ratios of 4 and 40 respectively (**Additional file 1**: **Figure S21**) although their roles are unclear since they do not cluster with any of the CTS topics (**Fig. 4b**). This reiterates the usefulness of using CTS topics in conjunction with *de novo* bulkRS topics to enable their interpretation wherever possible.

A similar analysis conducted using a separate scRNA-seq reference from healthy pancreatic subjects only reveals similar results (**Additional file 1**: **Figure S22**). However, in this analysis, only two of the *de novo* bulkRS topics clearly correspond to CTS topics (Acinar and Ductal), suggesting that the usage of an appropriate scRNA-seq reference with matched tumor environment is preferred.

### Deconvolving breast cancer transcriptomes revealed subtype-specific markers

To capture specific subtypes of breast tumor samples from the TCGA data, we trained GTM-decon on an scRNA-seq reference data from 26 primary tumors of breast cancer (BRCA) patients with three major clinical subtypes of BRCA, including 11 ER+, 5 HER2+ and 10 TNBC [44]. The data consists of about 1 million cells and covers 7 major cell types and 29 minor cell types. This served as a high-resolution reference for annotating the TCGA-BRCA tumor samples (n=1212) based on the major subtypes from the scRNA-seq dataset, namely Cancer-Basal, Cancer-Her2, Cancer-LumA, and Cancer-LumB (**Additional file 1**: **Figure S23a**). As expected, significantly higher proportions of Endothelial and Myoepithelial cell types are found in the normal-like samples, in comparison to the cancer subtypes (**Figure S23a**). Furthermore, Basal subtype is enriched for Cancer-Basal cells (**Figure S23b**, shown in dotted brown rectangle), and the Cancer-Associated Fibroblasts (CAFs) are enriched in almost all the samples (**Figure S23b,** shown in blue rectangles). Similar to the above analyses, some of the *de novo* bulkRS topics from sparsified samples overlap with the most represented cell types. For example, Topic 6 resembles Myofibroblast-like cancer-associated fibroblasts (myCAF-like), and Topic 8 resembles LumA subtype cancer cells.

Deconvolution using the scRNA-seq reference from the healthy individuals also captures the cell-type of origin for the different breast cancer subtypes implicated in the bulk samples (**Additional file 1: Figure S24**). For this analysis, using highly variable genes is more discriminatory than all genes (**Additional file 1: Figure S25**). The inferred CTS topic distributions from normal breast tissue recapitulates several marker genes from CellmarkerDB and PanglaoDB (**Figure S24a**). Furthermore, GTM-decon-inferred cell-type proportions clearly distinguish TCGA breast tumor samples from GTEx normal breast tissues (**Figure S24b**). Moreover, the Basal cell type is predicted to have the highest proportion in the Basal subtype defined by the PAM50 classification in comparison to other subtypes (**Figure S24c**) [45]. Also, we observed higher predicted proportion for Luminal_2 cell type in both LumA and LumB subtypes as expected [46] (**Figure S24c**). Among the *de novo* bulkRS topics from sparsified samples, Topic 5 is highly enriched in LumB, while Topic 7 and Topic 8 are enriched in Basal and LumA subtypes (**Figure S24c**). The guided topic score for the Basal subtype also correlates with higher proliferation score as expected [45]. Specifically, the Basal cell type is enriched in this subtype (**Figure S24d**; enclosed in green rectangle), whereas Luminal_2 is depleted (**Figure S24d**; enclosed in green dashed rectangle). In contrast, ER+ samples appear to be enriched for Luminal_2 cells (Wilcoxon test p-value = 4.5e-10; **Figure S26**), as expected [46, 47]. Furthermore, Topic 7 clearly captures the Basal subtype (enclosed in brown rectangle in **Figure S24d**), whereas there is no clear topic capturing the ER+ phenotype.

### GTM-decon learns phenotype-guided gene topics specific to BRCA subtypes

Our guided topic mechanism is not limited to inferring CTS topics, but can be extended to inferring phenotype-specific topics (i.e., topics capturing phenotype-specific gene expression) (**Fig. 1b**), thereby discovering gene signatures of subtypes or different cancer stages. We applied this approach to study the differences between Basal and ER+ BRCA subtypes (**Fig. 5a-c**) and the difference between ductal carcinoma and lobular carcinoma (**Fig. 5d-f**). We modelled each phenotype using 5 topics each, based on a sparsified matrix of bulk RNA-seq data (**Methods**), resulting in a genes-by-phenotypes matrix **ϕ** with 5 topics per phenotype (**Fig. 5a**). We then ranked genes by the topic scores under each topic. Almost all the genes identified by our approach were also deemed as differentially expressed (DE) genes by DESeq2 differential analysis [48] (**Fig. 5a**). GSEA of the topics shows differences between the Basal and ER+ phenotypes, although there is not much difference among the 5 topics for Basal (**Fig. 5c**). Moreover, the trained GTM-decon confers accurate phenotype classification with 97% accuracy for discriminating Basal and ER+ (**Fig. 5b**) and 82% for discriminating ductal and lobular subtypes (**Fig. 5d**), which is comparable to the traditional supervised learning methods namely logistic regression and Random Forest (**Fig. 5f**). We also evaluated phenotype classification accuracy as a function of sparsification rate (described **in Additional file 1 Section S7** and illustrated in **Additional file 1: Figure S27)** suggesting the importance of sparsification for inferring topics from bulk RNA-seq data using GTM-decon. We also trained GTM-decon only on the highly variable genes in the unsparsified bulk RNA-seq samples. This results in lower classification accuracy (76% for ductal-lobular samples as compared to 82%, and 95% for Basal-ER+ samples as compared to 97%), which might be due to information loss (HVG=1391 for Basal-ER+ samples, HVG=445 for ductal-lobular samples). Training using DE genes between the two phenotypes identified by DESeq2 also results in lower classification accuracy of 83% and 75%, respectively. These experiments suggest that GTM-decon can utilize more informative genes to discriminate the breast cancer subtypes than the traditional differential analysis approach.

**Figure 5.**
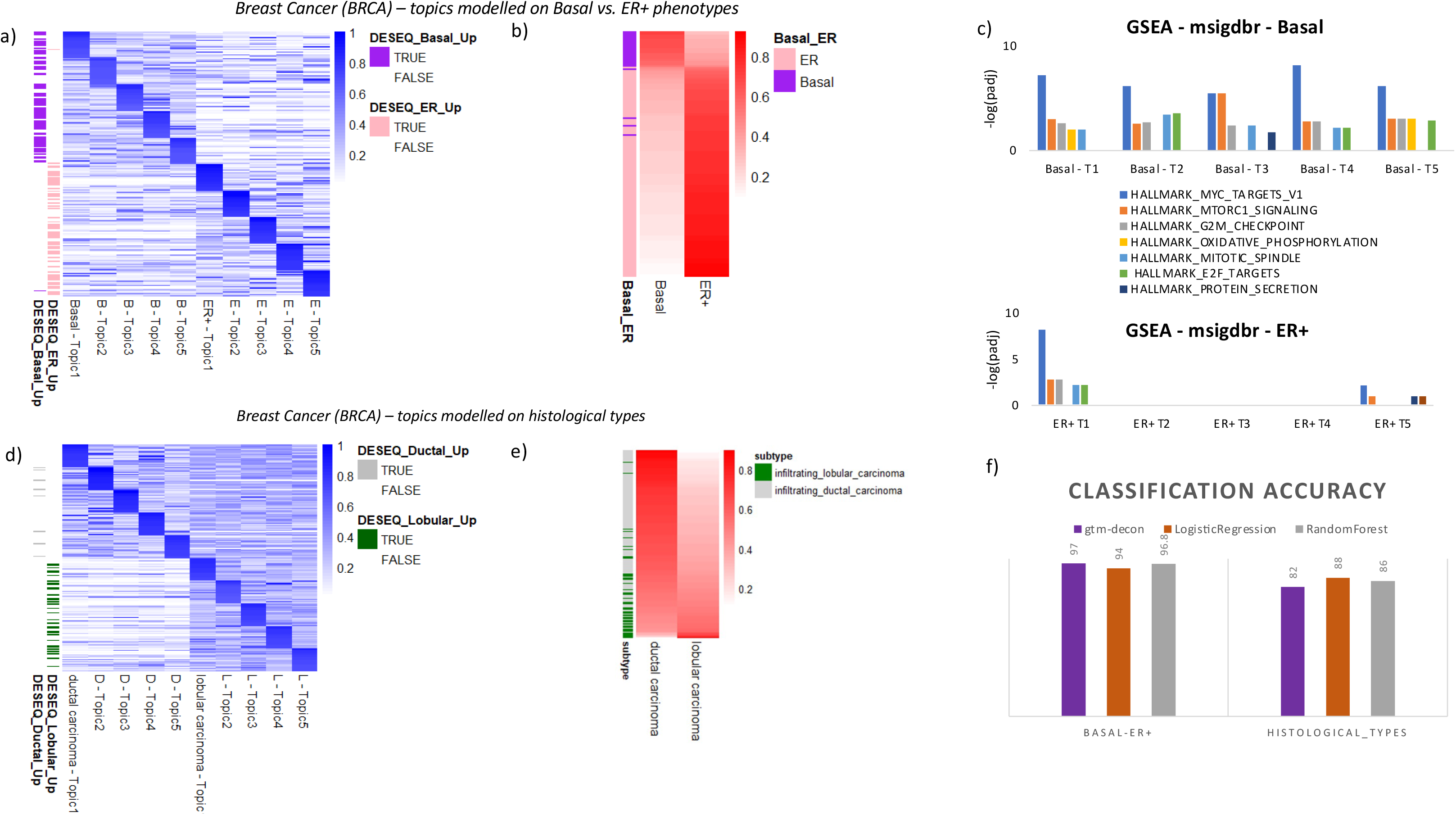
Phenotype-guided topic modeling of bulk RNA-seq data of breast cancer. **a**) Predicted top genes for the phenotype-guided topics for Basal and ER+ breast cancer subtypes. GTM-decon was trained on the sparsified TCGA-BRCA bulk RNA-seq data with the Basal and ER+ cancer subtypes as the guide. Five topics were used per subtype and therefore 10 topics in total. The heatmap illustrates the topic probabilities of the top 20 genes from each topic. As a comparison, the genes were also labelled as up or down-regulated if they were deemed differentially expressed by the DESeq2 analysis. **b**) Classification of Basal and ER+ subtypes based on phenotype-guided topic scores. The 5 topics for the same subtype were summed to obtain the overall score for Basal and ER+ subtype. Subjects in the rows were sorted by their Basal topic scores. **c**) GSEA analysis of the Basal and ER+ subtype topics. Significantly enriched MSigDb HALLMARK pathways were identified for each topic and displayed as barplots. The heights of the bar indicate the -log10 adjusted p-values and the colors indicate enriched pathways. **d)** Predicted top genes for the phenotype-guided topics for histological subtypes. Same as in panel **a**) but for Ductal and Lobular subtypes. **e)** Classification of histological subtypes. Same as in panel **b**) but for Ductal and Lobular subtypes. **f**) Evaluation of the subtype classification accuracy on the test breast tumor samples. We trained the phenotype-guided GTM-decon separately on 80% of the sparsified TCGA-BRCA tumor samples using Basal/ER+ and Histological types as the guides and evaluated its phenotype prediction accuracy on the 20% held sparsified samples. As a comparison, we also trained and evaluated Logistic Regression and Random Forest on the same training and test split, respectively. The classification accuracy on the test set by each method were displayed in the barplots.

### Nested-guided topic modeling identifies CTS DE genes in breast cancer subtypes

scRNA-seq can facilitate molecular understanding of genes and pathways in specific cell types with respect to phenotypic states. This level of detail is absent in bulk RNA-seq data, which profiles only the averaged gene expression from all cell types in the tissue. However, due to the cost, scRNA-seq profiles at the patient cohort size is rare. To take advantage of both types of data, we sought a way to identify cell-type specific gene expression differences corresponding to the phenotypes observed for the bulk samples. Briefly, we took a pretrained GTM-decon on a scRNA-seq reference data and then updated its CTS topics based on the corresponding phenotypes from the sparsified bulk data (**Fig. 1c**). This is equivalent to treating the phenotype as level 1 and cell types as level 2 in a two-stage nested factor design in statistics.

For the TCGA-BRCA data, in particular, we first initialized the genes-by-topics matrix **ϕ** with the *pre-trained* guided topics learned from the scRNA-seq reference for normal breast tissue. We then *fine-tuned* 5 CTS topics for each phenotype (i.e., ER+ or Basal) from sparsified bulk data, resulting in a 10-topic model. During the fine-tuning, all CTS topics corresponding to the patient’s phenotype are assigned a prior value of 0.9. Applying this approach to 798 sparsified samples from TCGA-BRCA, corresponding to Basal (N_1_=140) and ER+ (N_2_=658) led to a new genes-by-topics matrix **ϕ**^∗^. First of all, the top genes for the topics corresponding to the same cell type are similar in both phenotypes, as they are expected to capture CTS signatures (**Fig. 6a**). However, differential analysis of genes between two phenotypes revealed up-regulated genes in one phenotype being down-regulated in the other (**Fig. 6b**). These differences between Basal and ER+ are statistically significant across all genes for all the cell types based on paired Wilcoxon signed-rank tests. Next, we evaluated the ability of the fine-tuned GTM-decon to classify the held-out test set and observed a 97% classification accuracy (**Fig. 6c** top panel). These higher-resolution deconvolved CTS breast cancer profiles reveal that ER+ samples are enriched for Luminal-2 cell type, whereas the Basal subtype is depleted for that cell type (**Fig. 6c**).

**Figure 6.**
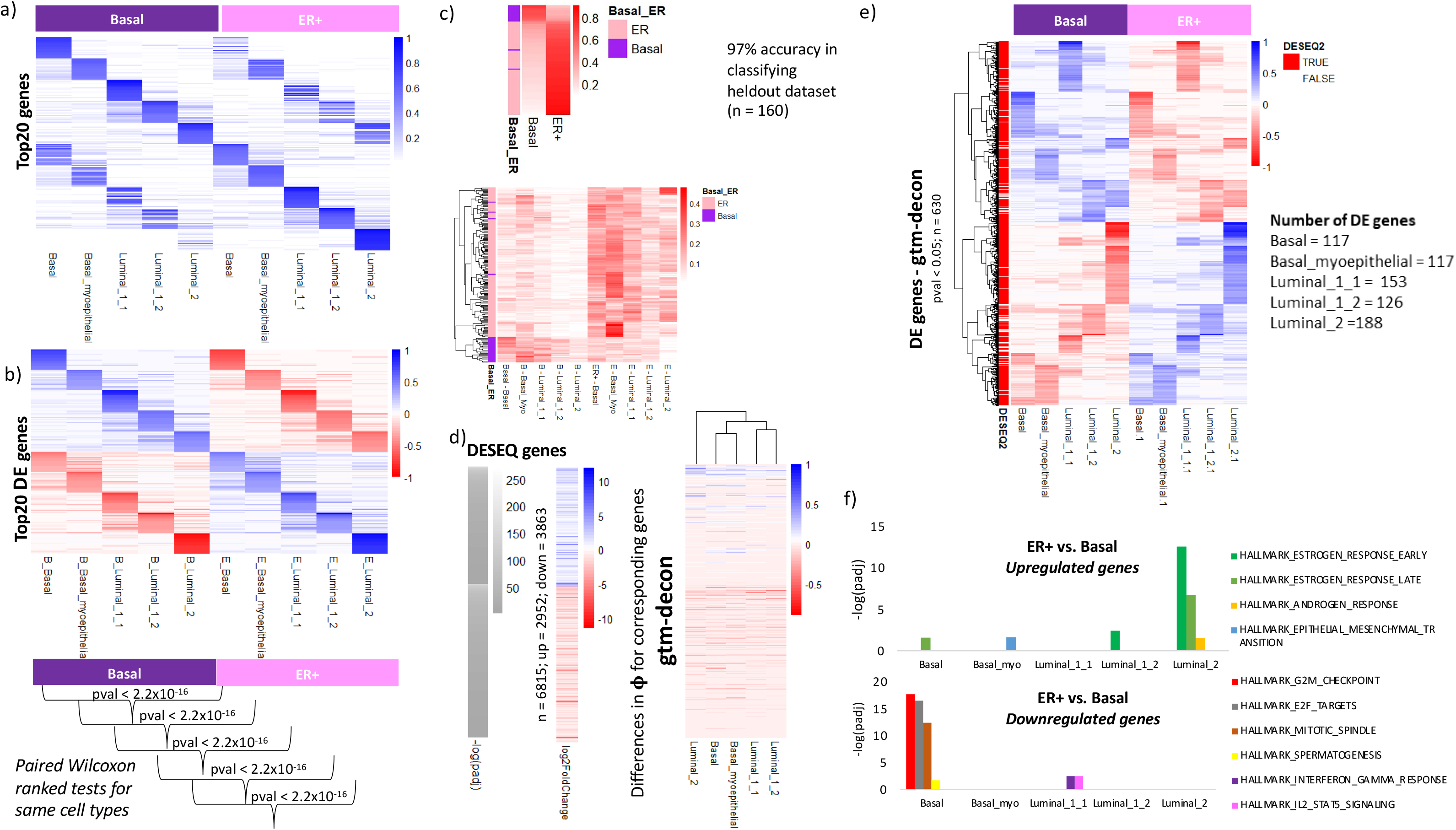
Identification of cell-type-specific differentially expressed genes from bulk RNA-seq data for Basal vs. ER+ subtypes. a) Top cell-type-specific gene signatures for Basal and ER+. GTM-decon was pretrained on a scRNA-seq reference dataset from normal breast tissue to infer the expression distribution of 5 cell types, namely Basal, Basal myoepithelial, Luminal 1-1, Luminal 1-2, and Luminal 2. The resulting genes-by-cell-types estimates were then used as the initial topic distributions for another GTM-decon, which is guided by the Basal and ER+ cancer subtypes in modeling the sparsified TCGA-BRCA bulk data. This led to a 10-topic distribution, each of which was specifically tailored for a combination of cell type and cancer subtype. The heatmap displays the probabilities of the top 20 genes for each topic. The left half displays the cell-type-specific topic distribution for Basal and the right half for ER+. **b)** Predicted differentially expressed (DE) genes for each cell type between Basal and ER+. The top DE genes for Basal in contrast to ER+ were identified by subtracting the gene topic scores for ER+ from the gene topic score for Basal under the same cell type. The resulting DE scores were shown in the top half of the heatmap. The bottom half displays the DE scores of the top genes for ER+ in contrast to Basal. The pairwise Wilcox signed-rank tests were performed to compare the gene topic scores across all genes between the two subtypes for the same cell type. All tests yielded p-values lower than 2.2e^-16^. **c)** Classification of Basal and ER+ based on the phenotype probabilities. As a validation for our nested phenotype-cell-type guided approach, we evaluated the classification accuracy on the 160 held-out sparsified breast tumor samples. For each subtype, we summed the cell-type-specific topic probabilities from bottom heatmap for each sample to obtain the phenotype scores, which are shown in the top heatmap. **d)** Comparison of DE genes detected by our approach and by DESeq2. DESeq2 was applied to the bulk RNA-seq gene expression data to compare gene expression between ER+ and Basal samples. In total, 6815 DE genes were deemed significant by adjusted p-value < 0.05 (Wald test) with 2952 up-regulated and 3863 down-regulated genes in ER+ relative to Basal. The grey bar and the heatmap on the left display the - log adjusted p-value for all of the upregulated genes (top half) and the down-regulated genes (bottom half). Genes were ordered in decreasing order of the absolute test statistic for each half. The corresponding log_2_ Fold-Change of ER+ over Basal was also shown as heatmap. The heatmap on the right displays the change of gene topic score from Basal subtype to ER+ subtype. **e)** Cell-type-specific DE genes identified by nested-guided topic approach. The top and bottom part of the heatmap displays the topic scores for the up-regulated and down-regulated genes in Basal relative to ER+, respectively (p-value < 0.05; permutation test). Genes that were also detected by DESeq2 were labeled in the color bar. **f)** ORA was applied to the differential topic scores of up-regulated and down-regulated genes in ER+ relative to Basal. MSigDb HALLMARK pathway gene sets were used in ORA. The -log p-values for the significant pathways were shown in the bar plot.

Next, we identified the DE genes per cell type by subtracting the genes-by-topics entries in the **ϕ** matrix for phenotype *d* (e.g., Basal) from phenotype *d*^′^ (e.g., ER+) under the same cell type (e.g., Luminal 2). To evaluate the consistency of our DE genes, we compared them against the DE genes identified by DESeq2 (**Fig. 6d**). We observe that all up-regulated and down-regulated DE genes nominated by DESeq2 in ER+ versus Basal comparison were also deemed up-regulated and down-regulated by our approach, respectively. We visualize the expression of the 630 statistically significant DE genes (adjusted p-value < 0.05; permutation test) in the BRCA tumor samples (**Fig. 6e**). We found that most of our DE genes do not only agree with those by DESeq2 but also exhibit CTS patterns. Notably, the most DE genes correspond to the Luminal_2 cell type (**Fig. 6e**), which exhibit the larger difference between ER+ and Bassal (**Additional file 1**: **Figure S26**). Over representation analysis (ORA) of these CTS DE genes against Hallmark pathways from MSigDb revealed several meaningful pathways. As expected, Estrogen Response (Early) and (Late) pathways are highly up**-**regulated in the ER+ phenotype, mainly in the Luminal_2 cell type (**Fig. 6f**). Similarly, most DE genes in the Basal phenotype are up-regulated for typical pathways involved in cancer, such as G2M checkpoint, E2F Targets, Mitotic Spindle. This reflects the aggressive nature of Basal cell type as the origin for the cancer subtype (**Fig. 6f**). The contribution of sparse genes (i.e. genes with zero counts due to sparsification) to CTS topics and DE analysis is described in **Additional file 1 Section S7** and illustrated in **Additional file 1: Figure S28**. We obtained similar results comparing the 2 histological subtypes – ductal carcinoma and lobular carcinoma (**Additional file 1**: **Figure S29**).

To further demonstrate the phenotype-guided and phenotype+cell-type guided functionality, we applied GTM-decon to the same scRNA-seq data from breast cancer tumors using phenotypes (i.e., ER+ and TNBC) and cell-type labels as the guides and then used the inferred topics to deconvolve the bulk TCGA breast tumor transcriptome data. Detailed analysis were presented in **Additional file 1 Section S8** and **Figure S30**. This was feasible because we have scRNA-seq references that were collected from similar tissue sites from patients of the same disease phenotypes as the target bulk transcriptomes.

## Discussion

In this study, we developed a Bayesian approach called *GTM-decon* to infer CTS gene topic distributions from scRNA-seq reference data. During the topic inference of each cell, we introduce the guidance by setting the topic hyperparameter for the cell type of that cell to be relatively larger than the hyperparameters for other topics. This enables us to anchor each topic to a specific cell type and subsequently guide the inference of the global topic distributions over genes to automatically prioritize cell-type marker genes. The resulting topic distributions can then be used to infer the relative cell-type proportions from bulk RNA-seq datasets (i.e., cell-type deconvolution).

Through our analysis of the pancreatic and breast tissue datasets, we observe that for those cell types, where marker gene information is available, most of the top genes under the CTS topics correspond to known marker genes (**Fig. 3a**; **Additional file 1**: **Figure S24a**). Because GTM-decon infers a distribution over all the genes under each cell-type-guided topic, it can be used to not only quantify the contribution of the known marker genes but also score novel marker genes. In terms of cell-type deconvolution, GTM-decon confers comparable performance to the existing state-of-the-art methods (**Fig. 2**; **Additional file 1**: **Figure S10-15**). The deconvolved cell-type proportions can be used to distinguish healthy and diseased samples as shown in the case of diabetic patients (**Fig. 3c**,**e**) as well as cancer subtypes from pancreatic and breast tumors (**Fig. 4** and **Additional file 1**: **Figure S23**). This enables investigation of molecular contribution to the phenotypic differences. Phenotypic differences between healthy and diabetic patients were captured even when the scRNA-seq reference datasets were generated from different platforms (e.g., Smart-seq and Drop-seq) (**Fig. 3d, f**).

Using GTM-decon, we revisited two cancer datasets from TCGA, namely pancreatic adenocarcinoma (PAAD) and breast cancer (BRCA). To dissect the heterogeneous tumor microenvironment, we deconvolved these datasets based on scRNA-seq reference sets from pancreatic cancer and breast cancer, respectively. As a result, we identified the ductal and acinar origin of PAAD, the endocrine origin of pancreatic neuroendocrine tumors (PNET) (**Fig. 4b**), and enrichment of subtype-specific cells for the BRCA subtypes (**Additional file 1**: **Figure S23a**). Interestingly, using scRNA-seq references from pancreatic and breast tissues of healthy individuals is also sufficient to identify the cell-type of origin for all subtypes in pancreatic cancer (**Additional file 1**: **Figure S22**), as well as the basal origin of Basal subtype and the luminal origin of ER+ breast cancer (**Additional file 1**: **Figure S24**). However, using a cancer-specific scRNA-seq dataset improves the resolution of deconvolution in identifying more subtypes (**Additional file 1**: **Figure S23**). By combining the *de novo* topics inferred directly from the bulk RNA-seq data with the scRNA-guided topics, we identified putative prognostic biomarkers that correlate with survival time (**Fig. 4c**; **Additional file 1**: **Figure S21**, **S22c**).

We further extended GTM-decon by modeling the sparsified bulk RNA-seq data using the patient phenotype labels as the guide rather than cell types. In contrast to the traditional differential analysis approach, the phenotype-guided GTM-decon provides a different way to investigate gene signatures and molecular pathways underpinning the phenotypes of interest (**Fig. 5**). To leverage the scRNA-seq reference data, we further extended this framework to a nested-guided topic model by fine-tuning a dedicated set of CTS topic distributions for each phenotypic state (e.g., BRCA subtypes) of the patients from the bulk RNA-seq data. This enables learning not only intra-phenotype changes of cell-type distributions but inter-phenotype changes of gene expression. The latter allowed us to identify CTS DE genes directly from bulk RNA-seq data (**Fig. 6**). We extended this approach to infer phenotype and cell-type guided topics from single-cell breast cancer RNA-seq data with cancer subtypes as the phenotype guide. This led to accurate deconvolution of cancer subtypes from bulk TCGA-BRCA data, as well as identification of phenotype-specific and CTS DE genes from the single-cell data (**Additional file 1: Figure S30**). Only a few methods can perform both deconvolution and CTS DE analysis. For example, CIBERSORTx [18] uses a non-negative matrix factorization approach based on partial observations to identify CTS DE genes across phenotypes. Other methods such as TOAST [24] and bMIND [23] that can estimate CTS DE require precomputed cell-type proportions by an external deconvolution method.

As future works, we can adapt GTM-decon to leverage other single-cell omic data such as scATAC-seq for reference and deconvolve the equivalent omic in bulk samples. It can also be extended to work with different cell states, like in BayesPrism [22], by modelling the topics based on cell states instead of cell types. Like all other deconvolution approaches, GTM-decon alone is unable to identify novel cell types from bulk RNA-seq datasets (i.e., cell types that are not present in the reference scRNA-seq data GTM-decon is trained on). while it is able to capture perturbed cell types based on the phenotypes (e.g., **Fig. 6c**). This can be addressed by training GTM-decon on an atlas-level scRNA-seq data such as the Human Cell Landscape [49], which comprehensively covers most of the cell types in the primary tissues. The resulting model can then be used to deconvolve bulk samples of any given target tissue. Lastly, cell types are not independent entities but rather form a lineage. One future direction is to exploit the relations among the cell types, which may better capture the underlying phenotypic states of the subjects. A more recently developed method called CeDAR [50] uses known cell-type hierarchy as prior to infer CTS expression in bulk data as opposed to our *de-novo* sub-cell-type inference and will leave a more detailed comparison as future work. Moreover, we can also extend GTM-decon to modeling multi-omic single-cell data to identify multi-omic CTS topic distributions and then use them to deconvolve multi-omic bulk data. To this end, while several multi-omic modeling methods have been developed [51–54], their benefits in deconvolution are not fully realized. Lasty, deep-learning-based methods such as Scaden [22] can train on simulated or real bulk RNA-seq datasets to predict cell-type proportions. While Scaden can confer accurate deconvolution results, it compromises interpretability because of its non-linear distributed representation of the gene expression features. Some recently developed variational autoencoder and embedded topic modeling frameworks [55, 56] may be extended to strike a balance between deconvolution accuracy and model interpretability.

## Conclusions

Computational cell-type deconvolution of heterogeneous bulk transcriptome is highly cost-effective in revealing the underlying phenotypic states and identifying CTS differentially expressed genes. GTM-decon represents a significant advance in the deconvolution methods with 3 prominent contributions: (1) automatic inference of interpretable CTS topic distributions by directly modeling large single-cell RNA-seq reference data without using known marker genes, therefore providing a principled and amenable reference map for the subsequent deconvolution; (2) identifying sub-CTS gene expression distributions by inferring multiple topics anchored at the same cell type, leading to a finer resolution of the deconvolution results; (3) detecting CTS DE genes directly from bulk or single-cell samples via an extended nested-guided topic design leveraging both the phenotype states and cell type label information. Our comprehensive experiments on pancreatic and breast datasets demonstrated the utilities of all 3 contributions. Together, GTM-decon is an efficient marker-free deconvolution method that takes the full advantage of the single-cell RNA-seq reference data for cell-type deconvolution.

## Methods

### Modeling scRNA-seq reference data via topic inference guided by cell-type labels

We adapted GTM-decon from the MixEHR-Guided (Mixture of Electronic Health Records – Guided) [57] (which was in turn inspired by MixEHR (Mixture of Electronic Health Records) [58] and sureLDA (Surrogate-guided ensemble Latent Dirichlet Allocation) [59]) to model gene expression from scRNA-seq data. We assume that the scRNA-seq data are generated from the following data generative process. Each cell indexed by m ∈ {1, …, M} is a mixture of the K cell types. The cell-type mixture **θ**_*m*_ is sampled from a K-dimensional Dirichlet distribution **θ**_m_ ∼ Dir(**α**_*m*_ + 0.1) with the hyperparameter **α**_*m*_ being specific to the cell. The key assumption here that separates GTM-decon from the standard LDA [25] is the use of noisy cell-type label *y*_*m*_ ∈ {1,.., *K*} for the cell. The hyperparameter corresponding to the cell-type label *y*_*m*_ = *k* has higher value (*α*_*m*,*k*_ = 0.9 by default) in contrast to the rest of the hyperparameters *α*_*m*,*k′*_ for *k*^′^ ≠ *k*, which are randomly set to small values between 0.01 and 0.1. Note that having non-zero *α*_*m*,*k′*_ allows other cell types to be assigned to the cell (i.e., **θ θ**_*m*,*k′*_ ≥ 0) and therefore **θ**_m_ reflects the statistical uncertainty of the cell-type label *y*_*m*_, which can be error prone due to various technical and data pre-processing aspects of the scRNA-seq data. To not clutter the notation, we omit the baseline value 0.1 in the following model description and use the more general form of **θ**_m_ ∼ Dir(**α**_*m*_) instead. Following the above default setting, this is equivalent to setting *α*_*m*,*k*_ = 1 for the observed cell type and *α*_*m*,*k′*_ ∈ [0.11, 0.2] for other cell types. For each cell *m*, each scRNA-sequenced read *i* ∈ {1, …, *N*_*m*_} originates from one of the *K* cell types with the probabilities dictated by its cell-type mixture (**θ**_m_): *z*_*i*,*m*_∼Cat(**θ**_*m*_). Given the cell type *z*_*i*,*m*_ = *k*, the *i*^*th*^ read maps to a specific gene indexed by *x*_*i*,*m*_ with the probabilities dictated by the CTS topic distribution over all genes (**ϕ**_*k*_): *x*_*i*,*m*_ ∼ Cat(**ϕ**_*k*_), where **ϕ**_*k*_ follows a G-dimensional Dirichlet distribution with a fixed hyperparameter *β* across all K dimensions: **ϕ**_*k*_∼Dir(*β*). Since the topic mixture **θ**_*m*_ are softly clamped to specific cell types via the K-dimensional hyperparameter **α**_*m*_, by following the above data generative process, it is straightforward to see that the K sets of topic distributions **ϕ**_*k*_ ‘s are also CTS.

The posterior distribution for the latent variables **θ**_*m*_’s, *z*_*i*,*m*_’s and **ϕ**_*k*_’s conditioned on the scRNA-seq reference data can be either approximated by Gibbs sampling [60] or by collapsed mean-field variational inference [61]. Specifically, for algorithmic convenience, we can leverage the conjugacy of the Dirichlet to Categorical distribution by integrating out **θ**_m_ and **ϕ**_*k*_ resulting in two Dirichlet-Multinomial distributions [62]:

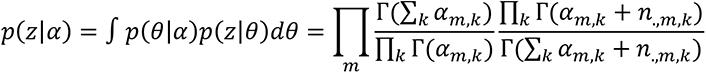

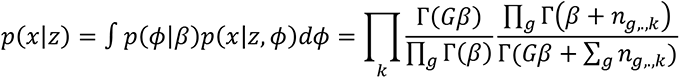

Note that we can recover the expected values for **θ**_m_’s and **ϕ**_*k*_’s given the posterior estimates of *z*_*i*,*m*_′*s* as they are proportional to the unnormalized counts *n*_.,*m*,*k*_ = ∑_*i*_[*z*_*i*,*m*_ = *k*]and *n*_*g*,.,*k*_ = ∑_*m*_[*z*_*i*,*m*_ = *k*, *x*_*i*,*m*_ = *g*], respectively.

The conditional distribution of the topic assignment for read *i* and cell *m* has a closed form expression:

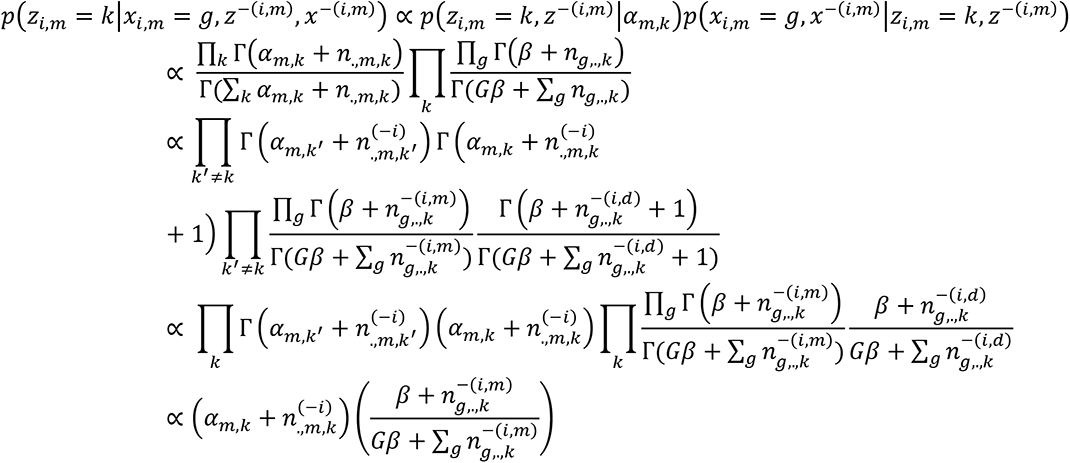

where the second last equality exploits the property of the Gamma function, i.e., Γ(*x* + 1) = Γ(*x*)*x*. Here 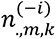 is the total number of scRNA-seq reads allocated for topic *k* for cell *m* without counting the current *i*^*th*^ read, and 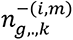 is the total read counts for gene *g* under topic *k* across all of the *M* cells, without counting the current *i*^*th*^ read in the *m*^*th*^ cell:

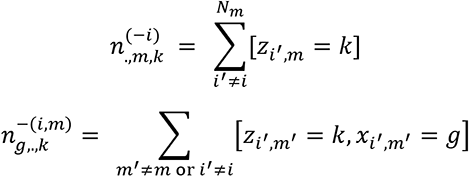

From here, the topic inference can be done by collapsed Gibbs sampling from *p*(*z*_*i*,*m*_ = *k*|*z*^−(*i*,*m*)^, *x*), while fixing the topic assignments for all other reads [60]. For a large number of cells in the scRNA-seq data, the collapsed Gibbs sampling approach tends to be slow in reaching an equilibrium state. Therefore, we took a deterministic mean-field variational inference approach, known as the collapsed variational Bayes (CVB) [61]. Specifically, we approximate the posterior distribution of the cell-type assignment *p*(*z*_*i*.*m*_|**x**, **α**_*m*_) via the variational Categorical distribution 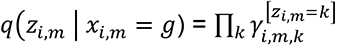, where

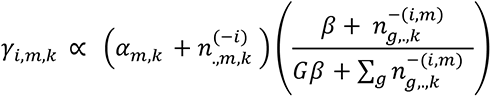

Using the variational parameters for the topic assignments, the above sufficient statistics are replaced by the soft counts:

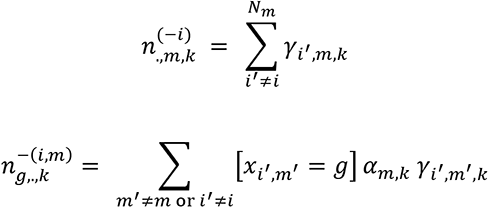

Here in updating the global gene distribution in the second equation, we further make use of the CTS prior **α**_*m*_ to obtain more interpretable results.

The above topic inference formulation operates at the level of read. For computational efficiency, our actual implementation of the topic inference was simplified to operate at the level of gene instead of the level of read:

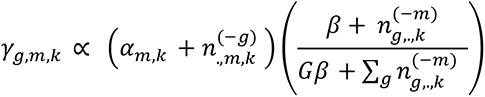

Similar to the above sufficient statistics, 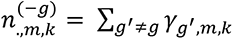 is the total number of scRNA-seq reads allocated for topic *k* for cell *m*, and 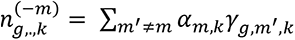 is the total read counts for gene *g* under topic *k* across all of the *M* cells, without counting the *g*^*th*^gene for cell *m*. This is equivalent to a reasonable assumption that all the reads from the same gene for the same cell are originated from the same cell type, i.e., ∀_*i*,*i′*_ *z*_*i*,*m*_ = *z*_*i′*,*m*_ if *x*_*i*,*m*_ = *x*_*i′*,*m*_.

Together, the inference algorithm alternates between two simple steps: (1) for each cell and each gene, perform coordinate ascent by computing *γ*_*g*,*m*,*k*_ while fixing the variational parameters *γ*_*g′*,*m*,*k*_ for other genes *g*^′^; (2) update the sufficient statistics. This algorithm maximizes the evidence lower bound *ELBO* = *E*_*q*_[log *p*(*x*|*z*)] + *E*_*q*_[log *p*(*z*|*α*)] − *E*_*q*_[log *q*(*z*|*γγ*)] [61]. The model is deemed converged when ELBO stops improving by a small threshold (1e^-6^ by default).

Upon convergence, the expected values for **θ**_m,k_ and *ϕ*_*g*,*k*_ are:

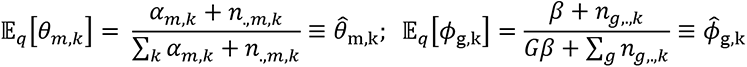

where 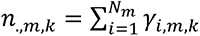 and 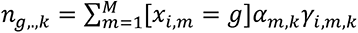. Here 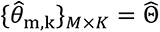 can be used to assess the “purity” of the single cells as a quality control step and 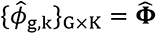 probabilities are used as the CTS topics in the subsequent deconvolution step.

### Inferring multiple topics per cell type

Suppose we use *L* topics per cell type for *K* cell types, the hyperparameter **α**_*m*_ for cell *m* can be formulated into a *K* × *L* matrix. Given that the cell-type label for cell *m* is *y*_*m*_ = *k*, we set the *k*^*th*^ row of **α**_*m*_ to relatively high values and the rest of the K-1 rows to relatively low values. For example, suppose we have 5 cell types and 3 sub-topics per cell type. If cell *m* is labeled with cell type 2, then the topic prior hyperparameter matrix **α**_*m*_ can be set to the following values: [0.01, 0.01, 0.01; 0.9, 0.9, 0.9; 0.01, 0.01, 0.01; 0.01, 0.01, 0.01; 0.01, 0.01, 0.01], where comma separates the columns and semicolons separate the rows. Here, the 3 topic prior values in the second row corresponding to cell type 2 are set to 0.9 and the values in the remaining rows are set to 0.01.

The data generative process is identical to the basic GTM-decon except having *K* × *L* topics instead of *K* topics. Specifically, to sample the cell-type mixture **θ**_m_, we flatten the *K* × *L* matrix for **α**_*m*_ to have a row vector of 1 × (*K* × *L*) so that the **θ**_m_ will have relatively high expected value for the *y*^*th*^ consecutive L values that correspond to the labelled cell type and relatively low expected value for the rest of the entries. We experimented modeling each cell type using *L* ∈ {2, 3, 4, 5} topics per cell type, with the hyperparameters *α*_*m*,*k*_ for each cell of cell type *k* set to 0.45, 0.3, 0.22, and 0.18, respectively. The prior values for the remaining topics were assigned with random values between 0.001 and 0.01. These priors were heuristically chosen based on the hyperparameter of value 0.9 for one topic divided by the number of topics per cell type.

### Inferring mixing cell-type proportions in bulk transcriptome

We assume a similar data generative process of the bulk transcriptome as the single-cell transcriptome described above. In particular, each bulk sample *j* ∈ {1,. ., *D*} is a mixture of *K* cell types. Its cell-type mixture **θ**_*j*_ is sampled from a K-dimensional symmetric Dirichlet distribution **θ**_j_∼Dir(**α**) with the hyperparameter fixed at a constant value across all K cell types (default: *α*_*k*_ = 0.1∀*k*). The flat hyperparameter value is used here since we typically do not have any prior information about the cell-type mixtures in the bulk RNA-seq data. For *N*_*j*_ total RNA-seq reads of bulk sample j, each read *i* ∈ {1, …, *N*_*j*_ } originates from one of the *K* cell types with the Categorical rates set to be **θ**_j_: *z*_*i*,*j*_ ∼Cat(**θ**_*j*_), where *z*_*i*,*j*_ ∈ {1, …, *K*}, and maps to a specific gene indexed by *x*_*i*,*j*_ ∈ {1, …, *G*} with a *known* CTS Categorical rates 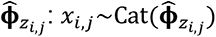.

Performing deconvolution on a bulk transcriptome profile is equivalent to inferring the posterior distribution of the CTS topic mixture given its gene expression and the CTS topic distributions: 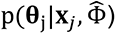. To this end, we used the G × *K* genes-by-CTS-topics estimates Φ̂ inferred from the scRNA-seq reference data (described in section “**Modeling scRNA-seq reference data via topic inference guided by cell-type labels**”) and perform variational inference to infer the cell-type mixture **θ**_*j*_ for the *j*^*th*^ bulk RNA-seq profile. Implementation-wise, similar to the scRNA-seq topic modeling, we also use the simplified topic inference algorithm at the gene level *g* ∈ {1, …, *G*} as opposed to at read level. This involves alternating between the topic assignment inference for ^*γ*^*g*,*j*,*k* and computing the sufficient statistics 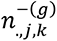 while fixing **Φ̂**. Algorithmically, for each gene, we infer the topic assignments:

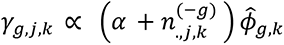

where 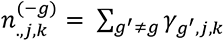. Upon convergence, we compute the expected CTS-topic mixture:

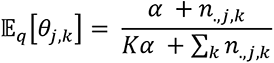

The bulk sample is transformed in the same way as the scRNA-seq reference data as described in the “**Preprocessing the reference scRNA-seq data**” section.

We also show that although GTM-decon infers RNA-fractions instead of cell-type fractions, the correlation between the RNA-fractions and cell-fractions is strong across cell types in multiple datasets, suggesting that RNA fractions per cell type can serve as a good surrogate to the cell fraction per cell type despite the potential differences in cell sizes among cell types (described in **Additional file 1: Section S9** and illustrated in **Figure S31**).

### Sparsification of the bulk data to directly infer topic distributions from them

In order to identify *de novo* topics that are not present in the reference scRNA-seq data, we applied standard LDA using the CVB implementation from MixEHR [58] to the sparsified bulk RNA-seq data. However, bulk RNA-seq data is a dense matrix with most genes having non-zero entries while topic models excel at modeling sparse matrices (e.g., scRNA-seq data). To make the bulk RNA-seq data amenable to our approach, we sparsified the dense matrix by setting all values below the 75^th^ percentile to 0. This cutoff was derived from scRNA-seq datasets, where on average 25% of genes in a cell have non-zero count values. While sparsifying works well in our applications, as a caveat, we acknowledge that it will lead to information loss. We also experimented with two other approaches to sparsify the matrix: (1) training GTM-decon only using HVG genes in bulk RNA-seq data; (2) training GTM-decon using differentially expressed genes identified between the phenotypes using DESeq2 [63].

Note that the sparsification procedure was done on the bulk data only when we directly inferred gene-topic distributions from them, which pertains to the *de novo* topic inference from bulk samples, phenotype-guided topic inference from the bulk, and nest-guided CTS-phenotype topic inference. All of the deconvolution experiments, where we first inferred topics from a single-cell reference dataset and then applied the inferred topics to deconvolve bulk data, does not involve the sparsification procedure (i.e., deconvolving the original bulk transcriptomes as they are). Identifying statistically significant marker genes per cell type

After GTM-decon topic inference, marker genes for each cell type were identified using permutation test. Specifically, for gene *g* under topic *k*, we computed the difference of its topic score *ϕϕ*_*g*,*k*_from the average topic score over the rest of the *K*-1 topics *ϕϕ*_*g*,*k′*≠*k*_ For example, for 3 cell types, the test statistic for cell type 1 is calculated as *ϕϕ*_*g*,1_ − *ϕ*_*g*,2_ + *ϕ*_*g*,3_/2. More generally, for *K* cell types and *T* topics per cell topic, the test statistic for cell type k and gene g is

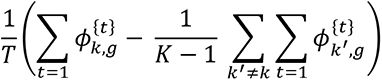

The significance of the observed statistic is compared against the same statistic calculated from 100,000 permutations. The empirical p-value is computed as fraction of permutations, where the test statistic is greater than the observed value.

### Evaluation of deconvolution accuracy using simulated and real bulk data

We evaluated the deconvolution accuracy by comparing the inferred cell-type proportions against the ground truth values for the datasets. First, we simulated bulk RNA-seq data using the scRNA-seq dataset. We summed the gene expression counts of each sample from the scRNA-seq data to represent the bulk data of that sample (artificial bulk transcriptome). The ground-truth cell-type proportions are the fraction of cells for each cell type. To avoid information bias in the evaluation, we performed leave-one-out-cross-validation (LOOCV) by inferring topics from scRNA-seq data for N - 1 subjects as the training examples and deconvolving the held-out subject as the validation example. The results were shown in **Additional file 1**: **Figure S10**.

In addition, we also used real bulk data with known cell-type proportion to benchmark our methods. To conduct different benchmarking and qualitative experiments, we used scRNA-seq datasets from several tissues for training GTM-decon (**Additional file 1: Table S1**): a) Human Pancreas (E-MTAB-5061, GSE81433) (with 4 Type 2 Diabetes and 3 healthy subjects) and mouse (GSE81433); b) Peripheral blood mononucleocytes (PBMC) of healthy human (GSE132044); c) Human blood cells (HBC) of healthy humans (GSE149938); d) Post-mortem brain tissue of frontal cortex from adult human (GSE97930); e) Breast tissue from healthy individuals (GSE113197). Apart from healthy individuals, scRNA-seq references from patients with a) Pancreatic cancer (CRA001160) and b) Breast cancer (GSE176078) were used to deconvolve cancer bulk RNA-seq datasets.

Bulk RNA-seq datasets with known ground truth values were chosen for benchmarking (**Additional file 1: Table S2**). Datasets with known ground truth proportions from flow cytometry include a) whole blood from 12 individuals (obtained from CIBERSORTx webpage) b) PBMC from a cohort of 13 individuals (GSE107011) c) PBMC from a cohort of 346 individuals (SDY67). For brain tissue, ground truth proportions from immunohistochemistry were available for a cohort of 41 individuals from the Religious Orders Study / Memory and Aging Project (ROSMAP) study (CortexCellDeconv). A unique dataset with both bulk RNA-seq data and scRNA-seq data from pancreas was accessed from E-MTAB-5060 and E-MTAB-5061, respectively, with the cell type proportions from scRNA-seq considered as ground truth values.

For the 3 immune (SDY67, Whole Blood, PBMC S13 cohort) and brain prefrontal cortex (ROSMAP) bulk datasets, we used two independent single-cell references with the most closely matched tissues of origin (**Additional file 1: Table S1**). For the pancreatic dataset (Segerstolpe), paired single-cell and bulk data from the same individuals were used in the LOOCV manner, where we used the cell-type proportions from the single-cell dataset to compare the estimated proportions from the paired bulk data in the same held-out subject.

To evaluate the concordance between the ground truth **y**_*m*_ and our inferred cell-type mixture **θ**_*m*_ for each test sample m, we computed four common metric scores: 1) Pearson correlation coefficient (PCC), 2) Spearman correlation (SCC), 3) cross entropy (CE), and 4) Residual mean squared error (RMSE) (**Fig. 2a**). Moreover, we also evaluated the deconvolution performance at each cell type using PCC and RMSE across samples (**Fig. 2b**; **Additional file 1**: **Figure S11-15**).

For the qualitative analyses on the cancer datasets, the bulk RNA-seq datasets for pancreatic cancer (PAAD) and breast cancer (BRCA) datasets from TCGA were downloaded from the GDC data portal (https://portal.gdc.cancer.gov/). Similarly, bulk RNA-seq data from the pancreas of a cohort of 89 normal and diabetic individuals was obtained from GSE50244.

### Implementation of the existing deconvolution methods

We compared GTM-decon with other state-of-the-art methods, including BSEQ-sc, BISQUE, MuSiC, CIBERSORTx, BayesPrism. For BSEQ-sc, marker genes were selected from CellmarkerDB for the brain and immune datasets, while the built-in pancreatic marker genes were used for the Segerstolpe dataset. BISQUE requires at least two paired reference bulk and reference single-cell samples, which is the case for the Segerstolpe dataset. When using Human Blood Cell (HBC) scRNA-seq data as a reference, in order to use BISQUE, artificial bulk samples were constructed using single-cell data. For the other 3 bulk datasets, since we used only one scRNA-seq reference, we left out BISQUE from the evaluation. For CIBERSORTx, all genes were provided to the web portal (https://cibersortx.stanford.edu/) for Signature Matrix generation and Cell Fraction imputation. For BayesPrism, all genes were provided to the web portal (https://www.bayesprism.org/) for cell type composition estimation, with the metadata column for tumor status set to 0 for all cells.

### Preprocessing the reference scRNA-seq data

The gene expression profiles of scRNA-seq data were used as training data. Since the performance of GTM-decon may vary depending on how the scRNA-seq count data are processed, we explored different gene selection and transformations of the scRNA-seq data. Specifically, the following gene selection were considered:

i. *all*: all genes without removal of any gene;
ii. *pp*: preprocessed genes to remove uninformative genes (frequently expressed genes – found in ≥ 80% of cells, infrequently expressed genes – found in ≤ 5 cells);
iii. *hvg:* highly variable genes identified using the highly_variable_genes (HVG) function of scanpy [64].

Furthermore, for each of these gene sets, the following transformation were considered:

i. *raw count* (all / pp / hvg) – scRNA-seq read counts per gene;
ii. *normr* (all_normr / pp_normr / hvg_normr) – normalize counts (while excluding highly expressed genes for calculating the normalization factor) as counts divided by the sum of counts per cell multiplied by a scaling factor of 10,000 (a commonly used factor for scRNA-seq data [65]), and round the values to their nearest integer to make the input suitable for topic modeling;
iii. *normr_log1p* (all_normr_log1p / pp_normr_log1p / hvg_normr_log1p): log-transform normalized counts in ii), and round the values to their nearest integer to make the input suitable for topic modeling.

### Gene set enrichment analysis (GSEA)

GSEA was performed using the fgsea package from R, on two different gene sets: a) Gene sets corresponding to specific cell types from CellMarkerDB [30]; b) Gene set corresponding to the HALLMARK pathways from MSigDb using msigdbr [66].

### Survival analysis

For TCGA datasets, after cell-type proportions are inferred using GTM-decon, survival analysis is performed using the survival and survminer R packages in order to assess if there is any correlation between survival probability of the cancer subjects and cell-type proportions for the cell types.

Kaplan-Meier curves were generated for those cell types, where the association is deemed statistically significant at p-value < 0.05 based on log-rank test.

### Learning phenotype-specific gene signatures from bulk RNA-seq data

To identify phenotype-specific genes, we applied GTM-decon to directly infer phenotype-topics from the sparsified bulk RNA-seq data (**Fig. 1b**). To evaluate phenotype prediction accuracy, we randomly split the bulk RNA-seq samples into training and test datasets in 80:20 ratio. The training dataset is guided in a manner similar to GTM-decon for the scRNA-seq datasets. Briefly, for each subject, the hyperparameter(s) for the topic(s) corresponding to the observed phenotype was (were) set to the prior value of 0.9, and the hyperparameters for the rest of the topics were set to a small value between 0.01 and 0.1. Since a model of 5 topics per cell type worked well for several scRNA-seq datasets, we opted to model each phenotype by 5 topics as well. For the subjects in the 20% test data, we summed up the inferred phenotype mixture **θ θ**_*d*,*k*_ values for all topics corresponding to the same phenotype. The predicted phenotype was the one with the highest summed up topic score. As a comparison, we also evaluated two other common machine learning methods namely logistic regression and random forest. We used their scikit-learn implementations with the default settings [67].

### Identifying CTS DE genes from bulk transcriptomes from nested-guided topics

In identifying phenotype-specific differentially expressed genes, we developed a way to leverage the CTS topics inferred from the scRNA-seq reference data by fine-tuning the CTS topics based on gene expression from the sparsified bulk RNA-seq data for different phenotypes (**Fig. 1c**). Specifically, we initialized the genes-by-topics matrix Φ_*d*_ for *each* phenotype *d*. For example, for *D* = 3 phenotypes in a tissue with *K* = 5 cell types, we will have 15 topics, comprising of 3 sets of CTS Φ_d_ matrices of 5 columns each. These matrices are guided by the phenotype labels during the variational inference.

For subject *j*, suppose his/her phenotype label is *d d* ∈ {1,. ., *D*}. The topic hyperparameters *α*_*j*,*d*,*k*_‘s for all the *K* topics corresponding to the phenotype label *d* were set to 0.9, and the *α*_*j*,*d′*,*k*_ for the other phenotypes were set to a small value between 0.01 and 0.1. These values were propagated throughout the variational inference, guiding the phenotypic inference appropriately. The inference algorithm is identical to the one described in “**Modeling scRNA-seq reference data via topic inference guided by cell-type labels**” section for modeling the scRNA-seq data. This approach also works with single-cell transcriptomes from a disease study such as the single-cell breast cancer study, where we not only have the cell type information but also phenotype states of the subjects (i.e., ER+ vs TNBC) (**Additional file 1: Section S8** and **Figure S30)**.

From the learned matrix, the test statistic of a DE candidate gene *g* between phenotypes *d* and *d*^′^ ≠ *d* per cell type *k* were identified by subtracting the *ϕϕ*_*g*,*d*,*k*_ value for phenotype *d* from the *ϕ*_*g*,*d′*,*k*_ value for phenotype *d*^′^. We identify statistically significant genes with a p-value < 0.05 by permutation test calculated from the 100,000 randomly shuffled Φ_*d*_ matrices. The p-value is the fraction of times the null statistic from the permutations is greater than the observed test statistic. As a reference, we compared our approach against the DE genes identified from the same samples using DESeq2 [63]. We identified hallmark pathways from MSigDb which were enriched for the DE genes by Over Representation Analysis (ORA) using the ClusterProfiler package from R [68].

## Declarations

### Ethics approval and consent to participate

Not applicable.

### Consent for publication

Not applicable.

### Availability of data and materials

The datasets analyzed during the current study were obtained from publicly available repositories or data portals. All the scRNA-seq datasets used for training in this study are listed in **Additional file 1: Table S1**, and all the bulk RNA-seq datasets with known ground truth proportions used in this study are listed in **Additional file 2: Table S2**. The Human pancreatic islet datasets from Segerstolpe et. al. used in this study is available in the ArrayExpress database under the accession code E-MTAB-5061. The Human and Mouse pancreatic islet datasets from Baron et. al. used in this study are available in the GEO database under the accession code GSE84133. The cancerous pancreatic tissue dataset is available in the Genome Sequence Archive database under the accession code CRA001160. The normal breast tissue dataset is available in the GEO database under the accession code GSE113197. The cancerous breast tissue dataset is available in the GEO database under the accession code GSE176078. scRNA-seq dataset for PBMC is available from the Single Cell Portal as PBMC2 or at GSE132044, and the dataset for human blood cells at GSE149938. Post-mortem brain frontal cortex from adult human is available under the accession code GSE97930. The RA synovium dataset is available in the ImmPort database under the accession code SDY998. The bulk RNA-seq dataset for normal and Type 2 Diabetes patients is available in the GEO database under the accession code GSE50244. The bulk RNA-seq datasets for pancreatic cancer (PAAD) and breast cancer (BRCA) datasets from TCGA can be downloaded from the GDC data portal (https://portal.gdc.cancer.gov/). Bulk RNA-seq datasets for PBMC can be accessed using the accession code GSE107011 from GEO database, and SDY67 from Immport database.

GTM-decon source codes have been deposited at the GitHub repository (https://github.com/li-lab-mcgill/gtm-decon).

### Authors’ contributions

Y.L. conceived of the study. S.S. and Y.L. analyzed and interpreted the data, wrote the manuscript, and wrote the code for GTM-decon. M.H. ran the experiments on benchmarking the deconvolutional accuracy. Y.L. supervised the project. All authors approved the final manuscript.

## Acknowledgements

We thank Yixuan Li for helping troubleshoot the GTM-decon software at its later stage.

## Additional file 1

### S1. Evaluation on gene selection strategies

We assessed the effects of using different genes selection strategies and different data transformations on the deconvolution performance. Specifically, we tested the effect of using all genes (ALL), preprocessed genes (PP) and only the highly variable genes (HVG) as input set for training GTM-decon. We observe that using all genes performed as well as or better than using only the HVG (**Figure S2-S4, S6**). Pre-processed genes vary in their performance depending on the dataset. The good performance on all genes also offers the added advantage of learning the individual contributions of each gene for each cell type in efforts of discovering novel marker genes.

### S2. Evaluation on raw count and transformation strategies

We experimented two transformation methods namely normalizing read counts (normr) and scaling followed by log-transformation (normr_log1p) as detailed in **Methods**. GTM-decon that operate on raw counts without any normalization performs the best for all the gene sets (**Figure S3**).

### S3. Experimenting hyperparameters **α**_m,k_ for cell-type mixture prior

We also experimented with different values for the hyperparameter values of the cell-type topic mixture prior **θ**_*m*_∼Dir(**α**_m_). In our default setting, we set the hyperparameter α_m,&_ to a relatively high value (i.e., 0.9 by default) given the cell-type label *y*_*m*_ = *k*; the rest of the K-1 α_*m,k′*_ values, where *y*_*m*_ ≠ *k*^’^, are set to a relatively low values (i.e., randomly sampled from a range between 0.1 and 0.01). To assess the impacts of the hyperparameter values, we varied the values for the CTS α_m,&_from 0.6 to 1. We also derived a data-driven prior from a multi-class logistic regression (MCLR) model that was trained to predict cell types of each cell. In particular, we use the fitted values of the MCLR in terms of the predicted probabilities as the prior values for each topic. The fitted values for all the correct cell types are above 0.8, suggesting a high confidence of the MCLR fit. We evaluated these prior settings on two cell-sorted purified immune cell datasets, GSE107011 (114 samples) and GSE64655 (48 samples) (**Table S2**). We find that the model is largely robust to the variation in topic prior (including the prior from MCLR), for all gene sets, for all the data sets (**Figure S4**). Visualizing the results as heatmaps also shows that varying the topics does not vary the clustering pattern, while confirming that the samples are largely clustered based on their primary cell type (**Figure S5**). We further evaluate on two real bulk datasets of whole blood and immune cells with known ground truth proportions (i.e., WB and S13 cohorts listed in **Table S2**) by assessing the correlation between deconvolved cell-type proportions from GTM-decon and ground truth proportions from flow cytometry data for each sample. The results also confirm that GTM-decon is robust to the variation of topic prior values in this range (**Figure S6**).

### S4. Experimenting hyperparameter *β* for CTS topics

By default, we set the hyperparameter *β* to 0.01, to let the scRNA-seq data likelihood drive the CTS topic distribution Φ. We vary this value between 0.0001 – 1 to evaluate its effect on the deconvolution of the whole blood dataset (WB) with known ground truth proportions, as well as on purified immune cells (**Figure S7**). The results show that GTM-decon is robust to the different values of the hyperparameter.

### S5. Experimenting number of topics per cell type

GTM-decon can infer multiple topics per cell type. While in most of our applications we find that the basic GTM-decon with one topic per cell type confers good performance, the GTM-decon with multiple topics per cell type tends to provide even better performance in terms of deconvolution accuracy on simulated bulk RNA-seq data using real scRNA-seq data (**Figure S8**).

### S6. Benchmark time and memory usage

We have benchmarked the time cost and memory usage as a function of the number of cells in the scRNA-seq reference data for GTM-decon in comparison with 3 SOTA methods, namely BISQUE, BSEQSC, and MuSiC. We did not benchmark CIBERSORTx because it is run remotely on a hosted web server. For the scRNA-seq reference, we used a combined PMBC datasets (**Table S2**) containing 30,000 cells, and 33,694 genes. Each method was trained to infer the cell type proportion in the purified bulk samples (GEO accession GSE64655). While the exact type of bulk data is not important in these time/memory benchmark experiments, we expect that each algorithm converges relatively fast due to the homogeneity of the bulk data. For GTM-decon, we used 1-5 topics per celltype (i.e., 9-45 topics in total for 9 cell types in the scRNA-seq reference data). For the other 3 methods, we used their recommended or default settings. We used the GNU time command to record the time and memory usage. GTM-decon scale linearly with both the number of topics per cell type and the number of cells (**Figure S17** top row), which is what we expected since its time and space complexity are both O(*n* × *G* × *K*) for N cells, G genes, and K topics. It also compares favourably with BISQUE, BSEQ-sc, and MuSiC in terms of running time and memory usage (**Figure S17** bottom row).

### S7. Effect of sparsification on phenotype classification

The sparsification procedure was done on the bulk data only when we directly inferred gene-topic distributions from them (i.e., only when we trained GTM-decon directly on the bulk data). This only pertains to inferring *de novo* topics from the bulk data using the standard LDA and the phenotype-guided topic inference from the bulk data. We observe that sparsification improved phenotype prediction accuracy on breast cancer data (TCGA-BRCA) for predicting ER+ and Basal subtypes compared to using the original bulk data as they are. Specifically, we varied the sparsification rates on the bulk data by setting all values below the n-th percentile to be zero and evaluated the prediction accuracies as a function of the sparsification rate. We observe increasing prediction accuracies for up to 90% as we increased the sparsification rate up to 60% (**Figure S27)**.

We sparsified values not genes. As a result, the percentage of the non-zero genes (i.e., genes with non-zero values in at least one sample after the sparsification) is about 70% even at 75% sparsification, suggesting that the model still contains a sufficient number of genes for capturing the cell types and phenotype specific variations. We investigated the contribution of the zero genes (i.e., genes that were removed after sparsification due to zero value across all samples). We observe that most of the zero genes occur in the lower 50th percentile of the CTS probabilities (**Figure S28**, left panel). Additionally, most of the zero genes do not feature in the top 25th percentile of the DE genes as determined by DESeq2 on the bulk TCGA-BRCA samples at FDR < 0.05 (**Figure S28**, right panel). Although differentially expressed in bulk, sparse genes do not contribute much to cell-type specific differences (**Figure S28**, left panel), and hence may not be captured by GTM-decon, which captures cell-type specific differences in gene expression.

### S8. Phenotype-CTS topic modeling of single-cell breast cancer transcriptomes for TCGA-BRCA bulk deconvolution

We can also infer both phenotype and CTS topics from the same single-cell breast cancer transcriptomes, which are composed of labeled cells from patients with 11 breast cancer patients of ER+ subtype and 10 patients with triple negative breast cancer (TNBC) subtype. From this dataset, we inferred CTS-phenotype topics simultaneously guided by the cancer subtypes (i.e., ER+ and TNBC) and cell types. We then a) applied the GTM-decon to deconvolve bulk TCGA-BRCA RNA-seq samples into CTS and phenotype-specific profiles; b) identified CTS-specific DE genes between ER+ and TNBC from the single-cell data; c) used the genes-by-CTS topics matrix to visualize the CTS distribution of DE genes identified from the bulk data.

Briefly, we modeled the phenotype-guided and cell-type-guided topic model by assigning one topic to each cell type per phenotype. During training, the topic corresponding to that cell type of that phenotype is set to a prior value of 0.9, and the rest of the topics (for the other cell types or phenotype) were set at a random value between 0.01 and 0.1 (**Figure S30a**). These priors guide the topic model to learn phenotype-CTS distribution. All the genes in the experiment were used as features.

We described the results presented in **Figure S30b-d** as follows:

1. The trained GTM-decon was evaluated by its classification accuracy in discriminating ER+ phenotypes from Basal-like phenotypes (the closest subtype to the TNBC subtype from the TCGA-BRCA data) in bulk RNA-seq data from TCGA-BRCA (n=1212). The left panel in **Figure S30b** illustrates that the samples corresponding to ER+ subtype are enriched for the ER+ topic. Interestingly, the samples corresponding to Basal-like subtype exhibit 2 different clusters - a cluster of samples showing enrichment for the TNBC topic (enclosed in green rectangle), and another cluster showing similar distributions of ER+ and TNBC topics (enclosed in brown rectangle). The former cluster likely corresponds to TNBC samples clustered into the Basal-like category, and the latter to others in the Basal-like category.
2. We deconvolved the TCGA-BRCA bulk samples without sparsification, resulting in a remarkably high phenotype classification accuracy of 92% using all genes, 90% using pre-processed genes, and 93% using highly variable genes. These results indicate that phenotype-guided topic modeling using scRNA-seq data can be effective in deconvolving bulk RNA-seq data as they are.
3. Furthermore, the CTS topics inferred for each phenotype enable us to visualize the deconvolution per sample at the CTS resolution for ER+ and TNBC/Basal-like subtypes (**Figure S30b**, right panel). We observe the enrichment of Cancer-Basal topic for TNBC (enclosed in orange rectangles), and enrichment of LumA and LumB topics for ER+ samples, as expected (enclosed in blue rectangles), and in concordance with the results obtained from cell-type guided topic modeling before (**Figure S23**).
4. Using the single-cell breast cancer transcriptomes alone, we identified CTS DE genes between the ER+ and TNBC subtypes based on the phenotypic differences in terms of the CTS probabilities per gene (**Figure S30c**). Based on 100,000 permutation tests of randomly shuffling the phenotype labels across cells, we identified 4,838 DE genes for at least one cell type at the empirical p-value < 0.05, 1,687 of which overlapped with the 5,936 DE genes identified from the TCGA-BRCA bulk RNA-seq dataset using DESeq2 at FDR < 0.05 for testing 20,501 protein-coding genes in total. Therefore, the overlap is statistically significant at p-value = 6.86E-25 based on Hypergeometric test. Similarly, as a baseline, we performed 100,000 permutation tests on the average CTS gene expression values derived from the single-cell data to compare the CTS gene expression difference between ER+ and basal-like subtypes. We identified 1892 DE genes for at least one cell type at the empirical p-value < 0.05, 592 of which overlapped with the 5,936 DE genes identified from the TCGA-BRCA bulk RNA-seq dataset using DESeq2 at FDR < 0.05 for testing 20,501 protein-coding genes in total. Therefore, the overlap is not as statistically significant as the above result at p-value = 0.01041539 based on Hypergeometric test.
5. We then visualized the expression difference of the DE genes from the bulk data in terms of the CTS-topic probabilities difference between ER+ and TNBC (**Figure S30d** left heatmap). We observe two distinct clusters of genes as highlighted in the orange and blue boxes, the former down-regulated in the ER+ phenotype topics (and up-regulated in TNBC), and the latter down-regulated in the TNBC phenotype topics (and up-regulated in ER+).

a. The cluster of genes upregulated in TNBC phenotype (enclosed in orange rectangle) consists of ∼1200 genes, predominantly upregulated in the Cancer-Basal, and Cancer-Cycling cell types. Based on over-representation analysis by ClusterProfiler on the MSigDB database, we found that these genes are enriched for E2F_TARGETS, G2M_CHECKPOINT, MYC_TARGETS_V2, and MITOTIC_SPINDLE pathways at FDR < 1.71E-18, 4.71E-11, 2.37E-05, and 1.07E-02, respectively, with an overlap of 58, 46, 17 and 29 genes. For the E2F_TARGETS gene set, some of the DE genes are MCM4, SYNCRIP, STMN1, TK1, SSRP1, CKS1B, EED, HELLS, USP1, and for the MYC_TARGETS_V2 set, some of the DE genes are MCM4, SRM, NDUFAF4, WDR43, WDR74, PLK1, PUS1, NOP2, RCL1, NIP7, SUPV3L1, MRTO4. The associations of these genes with the TNBC phenotype are supported by the literature. For instance, MYC_TARGETS_V1 and MYC_TARGETS_V2 are associated with tumor aggressiveness and poor survival in aggressive subtypes such as TNBC and HER2-positive breast cancer, when compared with ER-positive/HER2-negative tumors (MYC v1 and v2, both *p* < 0.001) [1]. Also, the genes encoding cell-cycle-related targets of E2F transcription factors (E2F_TARGETS) are associated with aggressive clinical characteristics, such as TNBC [2]. Higher G2M cell cycle pathway activity has been shown to be associated with worse clinico-pathologic features, with the TNBC subtype exhibiting the highest activity [3].
b. The cluster of genes up-regulated in ER+ cancer subtype (enclosed in blue rectangle) consists of ∼1000 genes, predominantly upregulated in the Cancer-LumA, Cancer-LumB, Cancer-Her2 cell types. These genes are over-represented in the Estrogen Response (Early and Late) pathways, with an overlap of 57 and 48 genes, at an FDR of 2.84E-25 and 9.54E-18, respectively. As the name suggests, these pathways correspond to genes up-regulated in response to Estrogen receptor activation, from which the ER+ subtype derives its name. Interestingly, 20 genes involved in Bile Acid Metabolism are also up-regulated at an FDR of 7.34E-03, which may be relevant as elevated bile metabolism is associated with better survival in breast cancer, with ER+ subtype exhibiting the highest bile acid metabolism in comparison to other subtypes [4].
c. Finally, we visualized the average gene expression in another heatmap (**Figure S30d** right heatmap), which exhibits a similar but much less salient pattern compared to the heatmap derived from the CTS-topic probabilities (**Figure S30d** left heatmap).

Together, these results further demonstrate the utility of GTM-decon in guiding phenotype and cell-type topic inference to identify disease-associated gene modules.

### S9. Effect of cell size on inference of cell-type deconvolution

Algorithmically, GTM-decon estimates the RNA fractions per cell type instead of cell fractions. Therefore, differences in cell sizes may lead to divergences between RNA fractions and cell fractions. Using the Pancreas reference data as an example, while we observe differences in the average RNA counts per cell among the seven cell types (**Figure S31a**), the relative proportion of RNA counts are quite similar to the relative proportion of the cells (**Figure S31b**). More generally, using the four scRNA-seq reference datasets that were used for the deconvolution of real bulk, we observed strong correlations between the total cell fraction and total RNA fraction associated with each given cell-type (**Figure S31c-f**). This suggests that RNA fractions per cell type can serve as a good surrogate to the cell fraction per cell type despite the potential differences in cell sizes among cell types (i.e., different average RNA counts among cell types).

### Supplementary Tables

**Table S1:** scRNA-seq datasets used as reference datasets for training.

| Dataset | Tissue | Sample size |  | Number of cells | Source | Reference |
| --- | --- | --- | --- | --- | --- | --- |
| **Segerstolpe** | Pancreas | 10 |  | 2209 | E-MTAB-5061 | Segerstolpe et al, Cell Metab, 2016 [5] |
| **Baron – Human** | Pancreas | 4 |  | 8569 | GSE81433 | Baron et al, Cell Systems, 2016 [6] |
| **Baron – Mouse** | Pancreas | 2 |  | 1886 | GSE81433 | Baron et al, Cell Systems, 2016 [6] |
| **Pancreas – Cancer** | Pancreas | 35 |  | 57530 | CRA001160 | Peng et al, Cell Res, 2019 [7] |
| **PBMC2** | PBMC | 1 |  | 11183 | Single Cell Portal / GSE132044 | Ding et al, Nat Biotech, 2020 [8] |
| **HBC** | Blood | 31 |  | 7643 | GSE149938 | Xie et al, Natl Sci Rev, 2021 [9] |
| **Lake_Human** | Brain | 1 |  | 10319 | GSE97930 | Lake et al, Science, 2016 [10] |
| **Breast - Normal** | Breast | 4 |  | 24646 | GSE113197 | Nguyen et al, Nat Comm, 2018 [11] |
| **Breast - Cancer** | Breast | 26 |  | 1000064 | GSE176078 | Wu et al, Nat Genet., 2021 [12] |

**Table S2:** Bulk RNA-seq datasets with ground truth proportions for evaluation.

| **Dataset** | **Tissue** | **Samples** | **Data Type** | **Source** | **Reference** |
| --- | --- | --- | --- | --- | --- |
| **Whole Blood (WB)** | PBMC | 12 | Bulk RNA-seq | CIBERSORTx – Fig2b | Steen et.al, Stem Cell Tran. Net, 2020 [13] |
| **PBMC S13 cohort** | PBMC | 11 | Bulk RNA-seq | GSE107011 | Monoco et. al. Cell Rep, 2019 [14] |
| **SDY67** | PBMC | 346 | Bulk RNA-seq | Immport SDY67 | Zimmerman et. al. Front Immunol, 2017 [15] |
| **PBMC - GSE107011** | Purified immune cells | 127 | Purified cell-sorted Bulk RNA-seq | GSE107011 | Monoco et. al. Cell Rep, 2019 [14] |
| **PBMC - GSE64655** | Purified immune cells | 56 | Purified cell-sorted Bulk RNA-seq | GSE64655 | Hoek et. al., PLoS One, 2015 [16] |
| **ROSMAP** | Brain | 41 | Bulk RNA-seq | CortexCellDeco nv tutorial | Patrick et. al., PLOS Comp. Biol, 2020 [17] |
| **Segerstolpe** | Pancreas | 7 | Bulk RNA-seq | E-MTAB-5060 | Segerstolpe et al, Cell Metab, 2016 [5] |

The ground truth proportions are available from the supplementary materials or web links of the references. Monaco et. al. contains ground truth proportions of PBMC S13 cohort and SDY67.

**Table S3:**
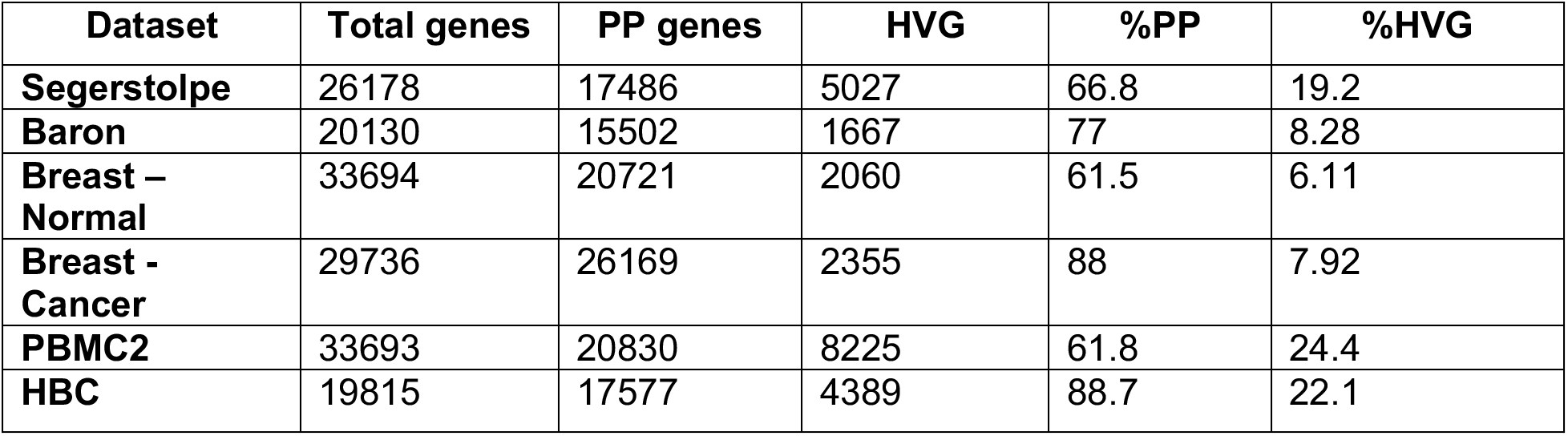
Percentage of preprocessed (PP) genes and HVG for scRNA-seq datasets.

| **Dataset** | **Total genes** | **PP genes** | **HVG** | **%PP** | **%HVG** |
| --- | --- | --- | --- | --- | --- |
| **Segerstolpe** | 26178 | 17486 | 5027 | 66.8 | 19.2 |
| **Baron** | 20130 | 15502 | 1667 | 77 | 8.28 |
| **Breast – Normal** | 33694 | 20721 | 2060 | 61.5 | 6.11 |
| **Breast - Cancer** | 29736 | 26169 | 2355 | 88 | 7.92 |
| **PBMC2** | 33693 | 20830 | 8225 | 61.8 | 24.4 |
| **HBC** | 19815 | 17577 | 4389 | 88.7 | 22.1 |

### Supplementary Figures

**Figure S1:**
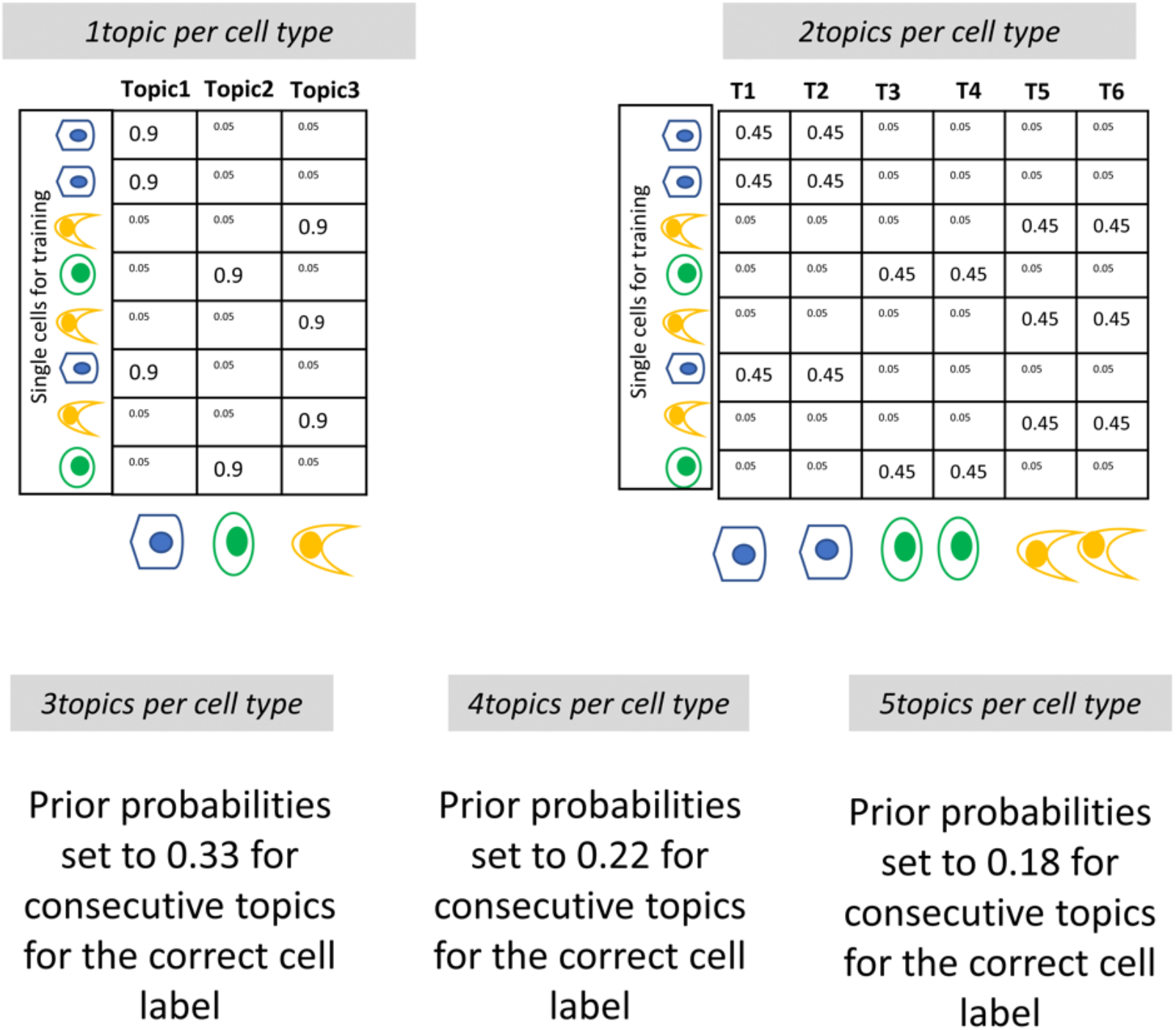
Modeling a cell type using multiple topics. GTM-decon can model each cell type via multiple topics (2, 3, 4, 5). It achieves this by setting the prior values to be assigned to the hyperparameter **α**_*m*,*k*_ for each of cell of cell type *k* to 0.45, 0.33, 0.22, and 0.18, respectively, for ‘x’ consecutive topics. For example, a 2-topic model with 3 cell types, as shown above, is modelled using 6 topics, with each set of 2 adjacent topics modeling a cell type. Here, the 2 adjacent topics modeling the cell type corresponding to cell ‘m’ are set to 0.45 and the remaining topics were assigned random probabilities between 0.001 and 0.01.

**Figure S2:**
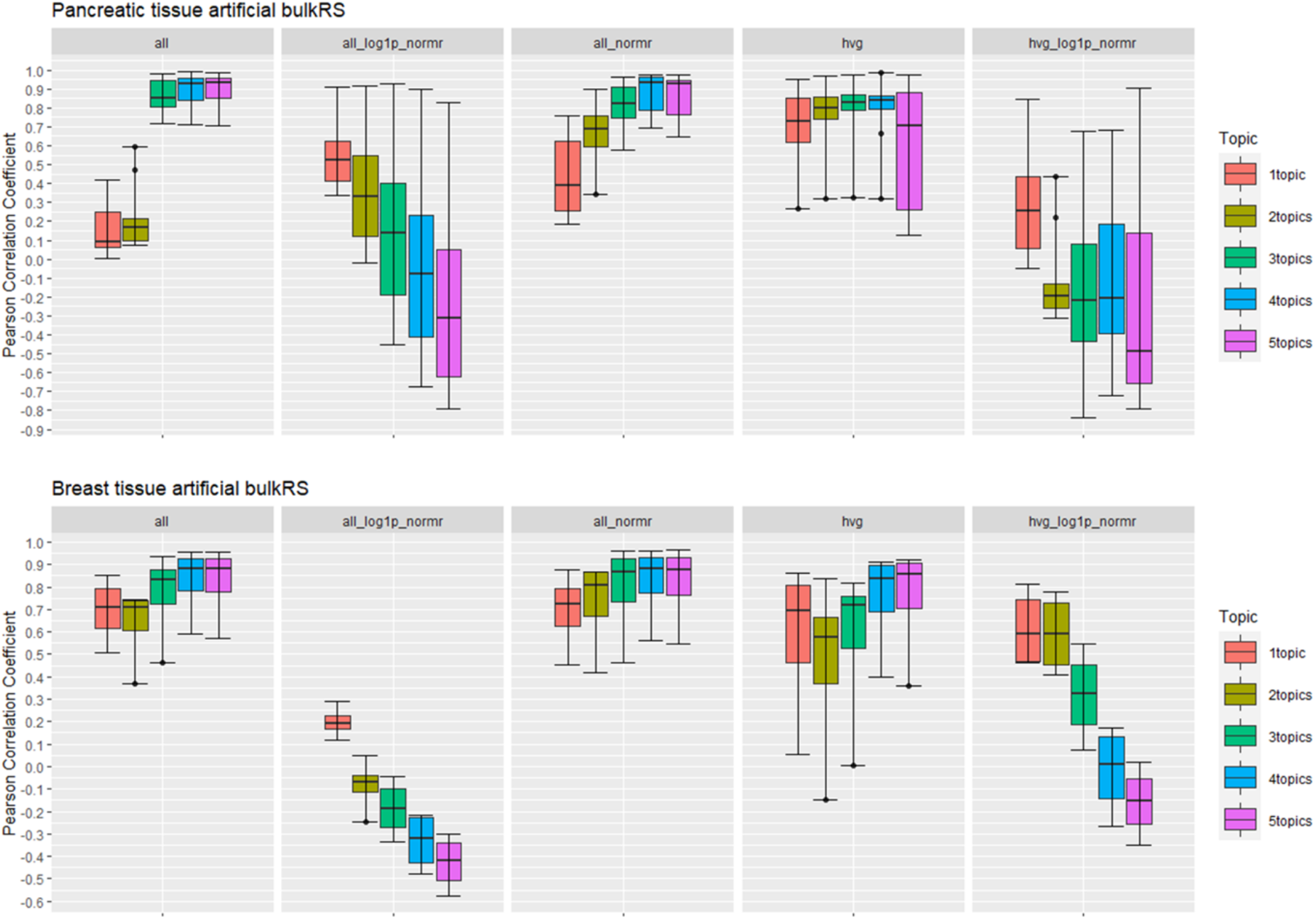
Deconvolution accuracy for different gene sets from simulated data. Each GTM-decon model was trained by modelling a cell type using different gene selection strategies and their transformations i) all genes – corresponding to raw gene counts; ii) all_normr – corresponding to normalized of raw genes by total counts in a cell; iii) all_log1p_normr – corresponding to log-transformed form of all_normr; iv) hvg – corresponding to raw counts of highly variable genes; v) hvg_log1p_normr – corresponding to log-transformed form of normalized hvg. Each of these models was further trained based on K number of topics, with K varying from 1 to 5. We evaluated the deconvolution accuracy of these models on simulated bulk data from two datasets: Pancreatic (Segerstolpe) and Breast (Normal). We simulated the bulk RNA-seq (bulkRS) data by randomly sampling cells from the held-out set of scRNA-seq data (20%) and summing up the values for each gene from all cells belonging to an individual. The cell proportions inferred for each individual was compared with the ground truth values and evaluated by Pearson correlation. The box and the whiskers in each boxplot indicate the 25%-75% quartile and min-max of the evaluation scores over the individuals, respectively.

**Figure S3:**
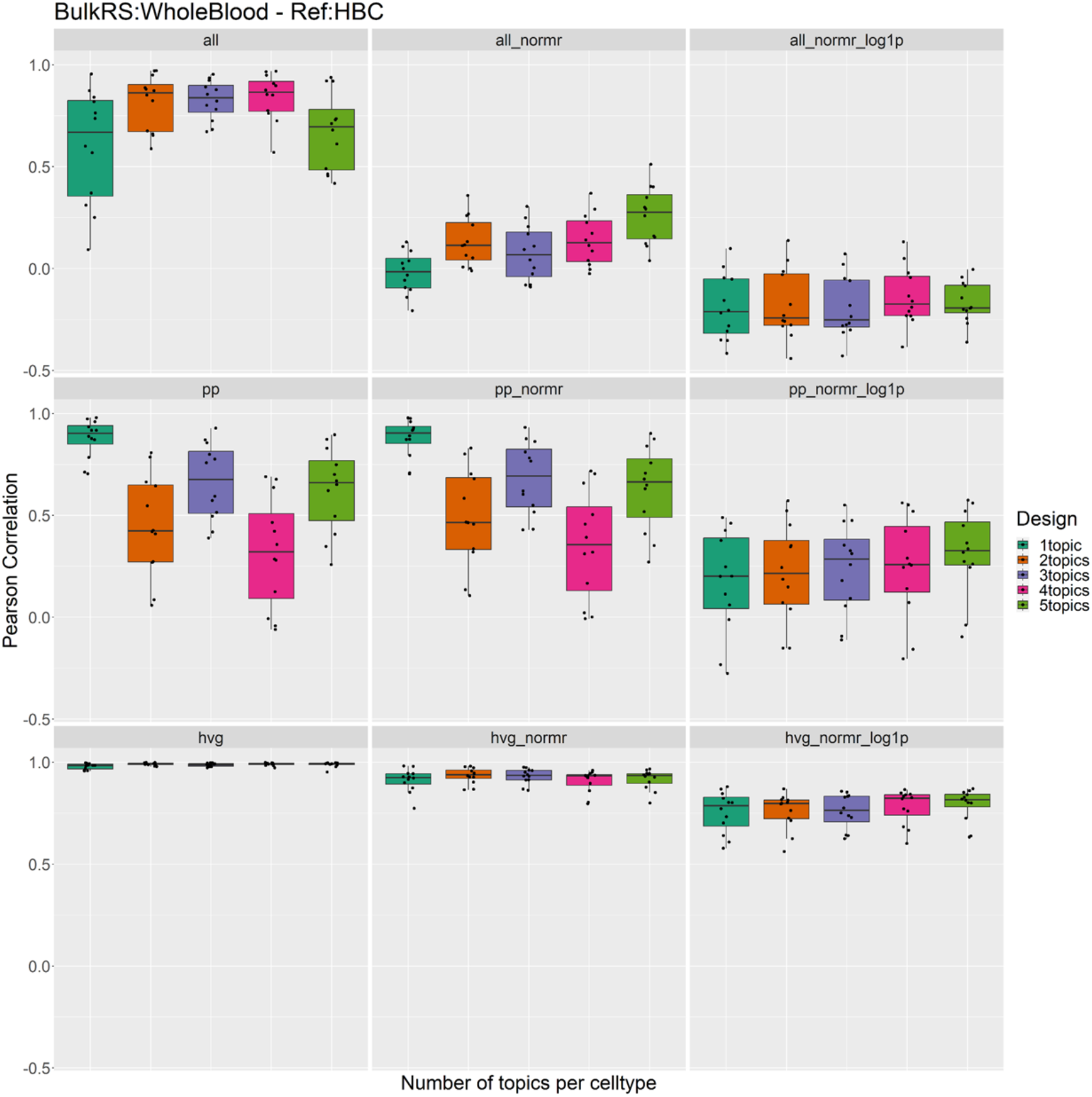
Deconvolution using different normalization strategies for different gene sets: We evaluate the performance of different gene sets and their normalizations on Whole Blood RNA-seq bulk dataset. The boxplots show the deconvolution accuracy in terms of Pearson Correlation between ground truth cell type proportions estimated by flow cytometry for each sample vs. deconvolved cell type proportions inferred by GTM-decon (y-axis). The box and the whiskers in each boxplot indicate the 25%-75% quartile and min-max of the evaluation scores for each of the samples. Deconvolution performance is shown for different models using a varying number of topics (1 to 5) to model each cell type. The box plots reveal that raw counts (i.e., no normalization) performs the best for all the gene sets, whereas normalization decreases the performance to varying degrees for different gene selection strategies.

**Figure S4:**
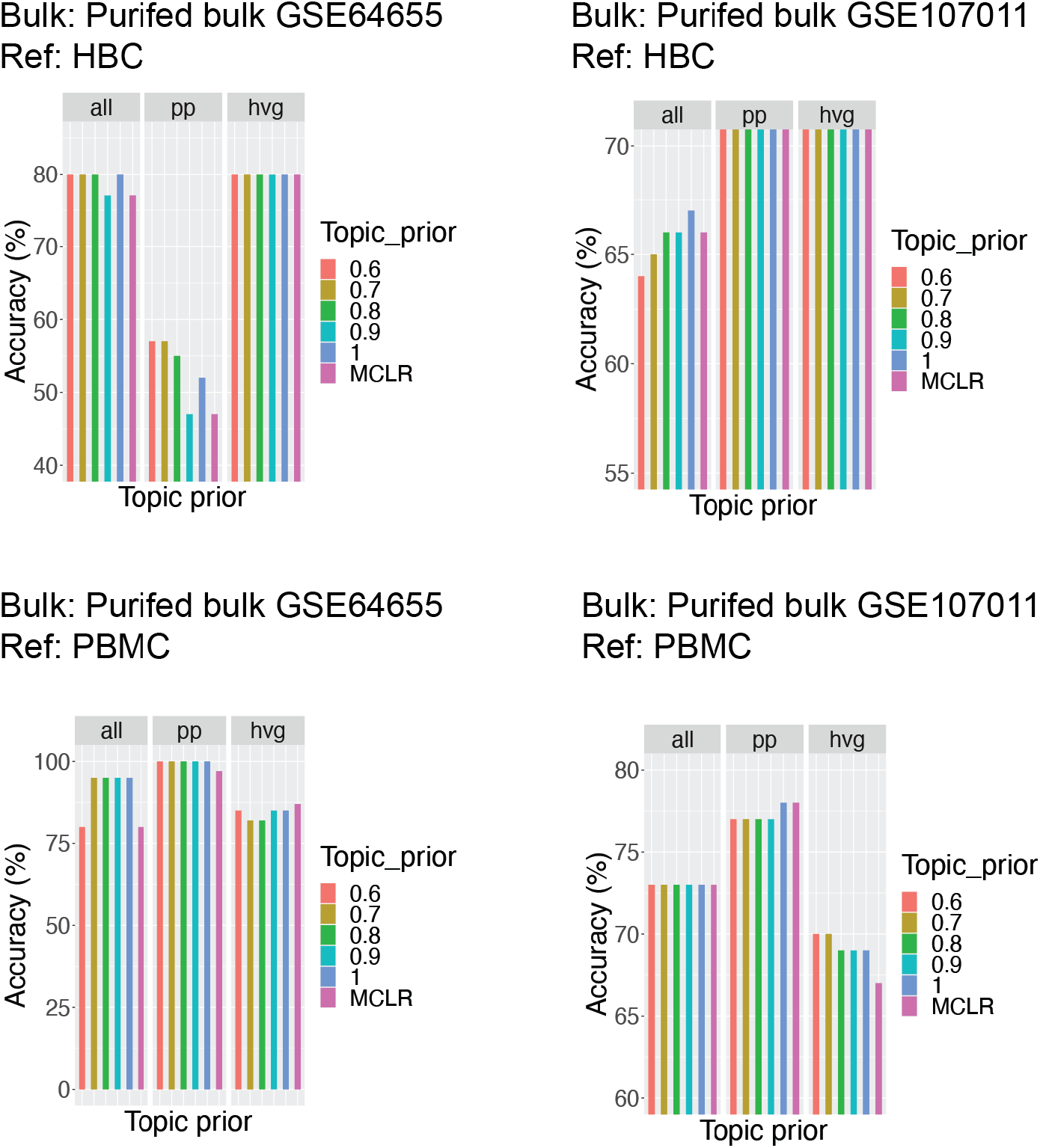
Effect of varying CTS topic prior on deconvolution accuracy in purified immune bulk RNA-seq samples. We evaluated the performance of GTM-decon on GSE107011 and GSE64655 using two reference datasets, HBC and PBMC2. For each purified bulk sample, the cell type corresponding to the highest inferred cell-type proportion by each method was used as the predicted cell type. The barplots show the prediction accuracy as the percentage of the correctly predicted samples. We varied topic prior hyperparameter **α**_*m*,*k*_ from 0.6 to 1, as well as topic priors derived from a multi-class logistic regression model fitting the scRNA-seq data to its cell types (MCLR). Three different gene selections namely all genes (ALL), preprocessed genes (PP), and highly variable genes (HVG) were experimented. The bar plots reveal that the method is largely robust to variation in topic prior.

**Figure S5:**
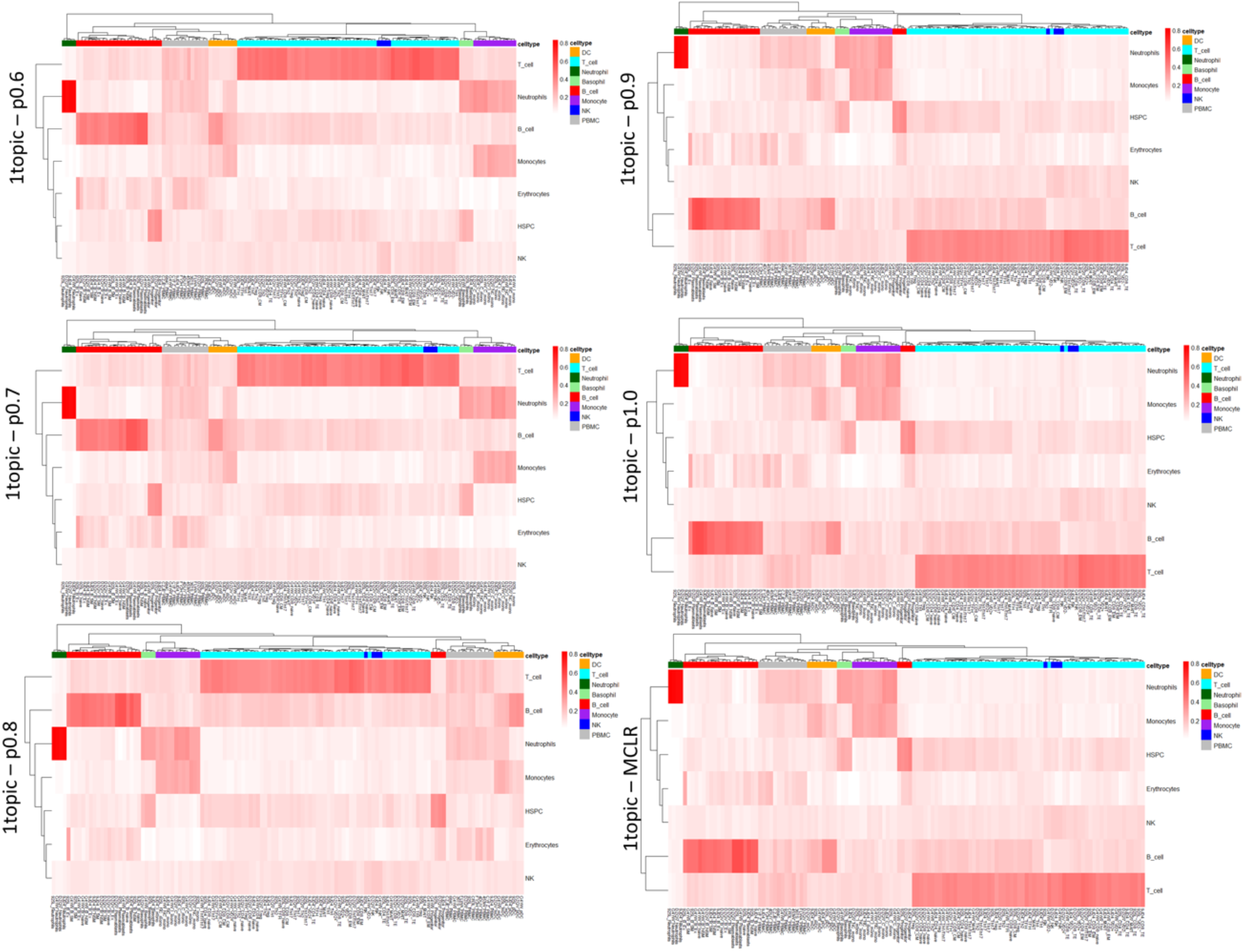
Heatmaps showing deconvolved cell type proportions when CTS topic prior is varied. The heatmaps plot the deconvolved cell type proportions from GTM-decon models using different CTS topic prior values, including prior values derived from a multi-class logistic regression model (1topic-MCLR), respectively. The rows correspond to CTS topics and the columns represent bulk RNA-seq samples derived largely from purified cell-sorted immune cells (GSE107011), except for the PBMC samples. The columns are colored based on the primary cell type, and shows that in most cases, the CTS topic with the highest deconvolved proportions corresponds to the correct cell type for the sample. The heatmaps for the different priors are also largely comparable, suggesting the robustness of the approach to different priors.

**Figure S6:**
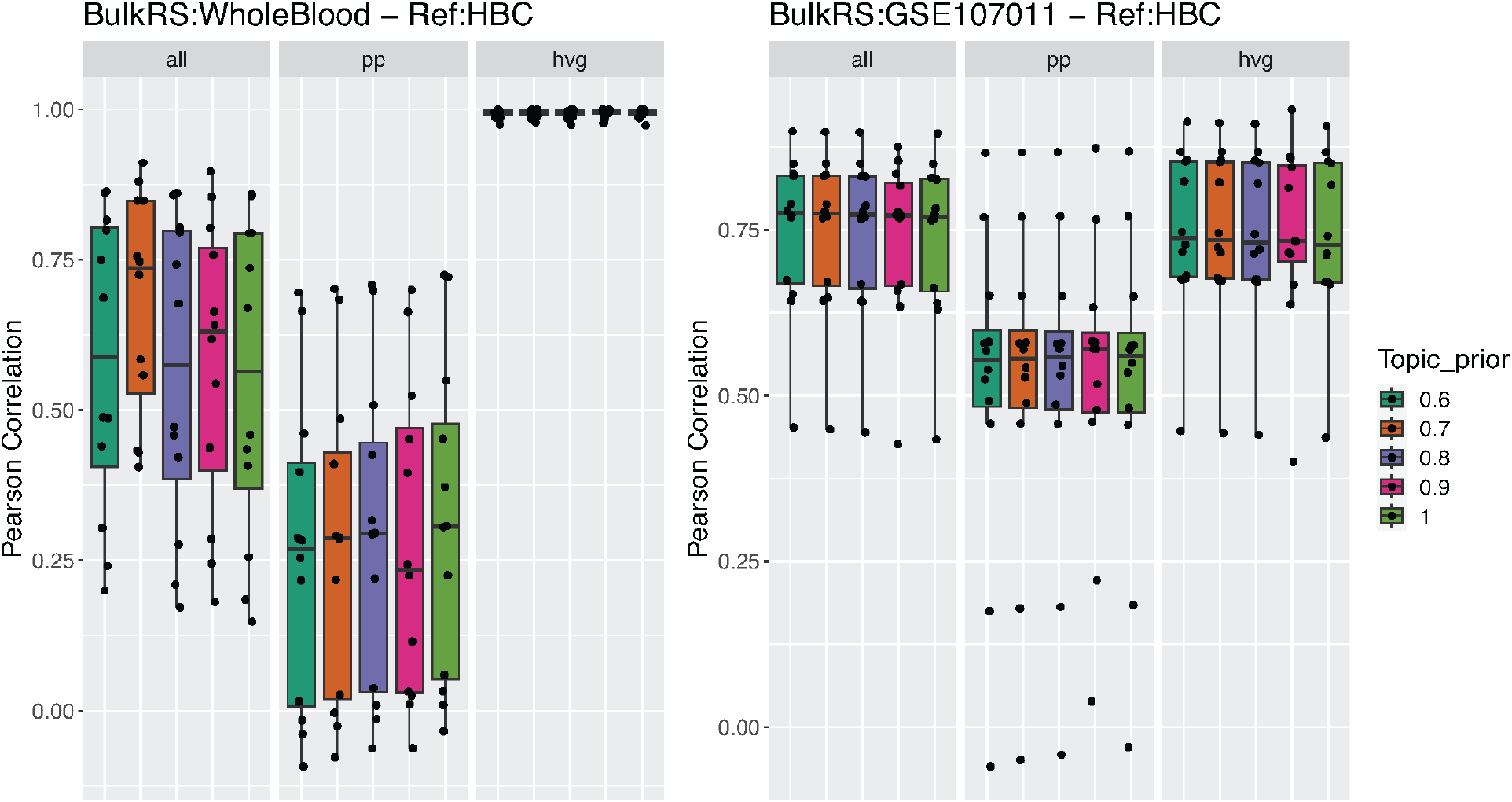
Effect of varying CTS topic prior on deconvolution accuracy in two bulk RNA-seq datasets with ground truth proportions. We evaluated performance of GTM-decon on Whole Blood (WB) and GSE107011 datasets, while varying the CTS topic prior from 0.6 - 1. The boxplots show the deconvolution accuracy in terms of Pearson Correlation between ground truth cell type proportions estimated by flow cytometry for each sample vs. deconvolved cell type proportions inferred by GTM-decon (y-axis). The box and the whiskers in each boxplot indicate the 25%-75% quartile and min-max of the evaluation scores for each of the samples. The bar plots reveal that the method is largely robust to variation in topic prior, with all genes and highly variable genes performing better than pre-processed genes.

**Figure S7.**
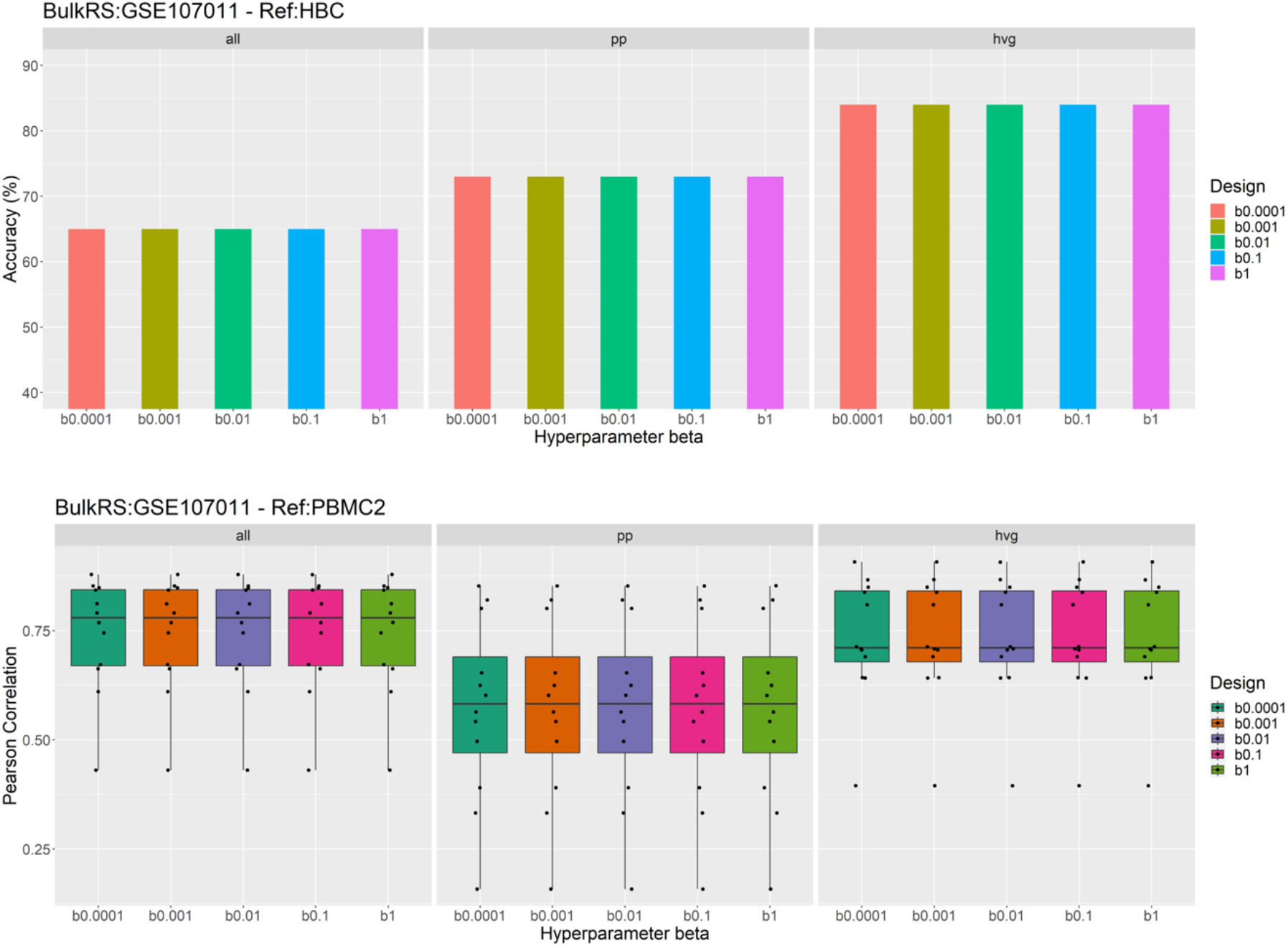
Effect of varying hyperparameter. *β* **on deconvolution accuracy.** Deconvolution accuracy was estimated in purified bulk samples (GSE107011) using HBC as the reference, and the real bulk data PBMC S13 cohort from the same dataset. The hyperparameter *β* was varied from 0.0001 to 1. For each purified bulk sample, the cell type corresponding to the highest inferred cell-type proportion by each method was used as the predicted cell type. The barplots show the prediction accuracy as the percentage of the correctly predicted samples. Similarly, deconvolution accuracy on real bulk RNA-seq (GSE107011, S13 cohort) was estimated n terms of Pearson Correlation between ground truth cell type proportions estimated by flow cytometry for each sample vs. deconvolved cell type proportions inferred by GTM-decon (y-axis). The plots reveal that the method is robust to variations in hyperparameter *β*.

**Figure S8:**
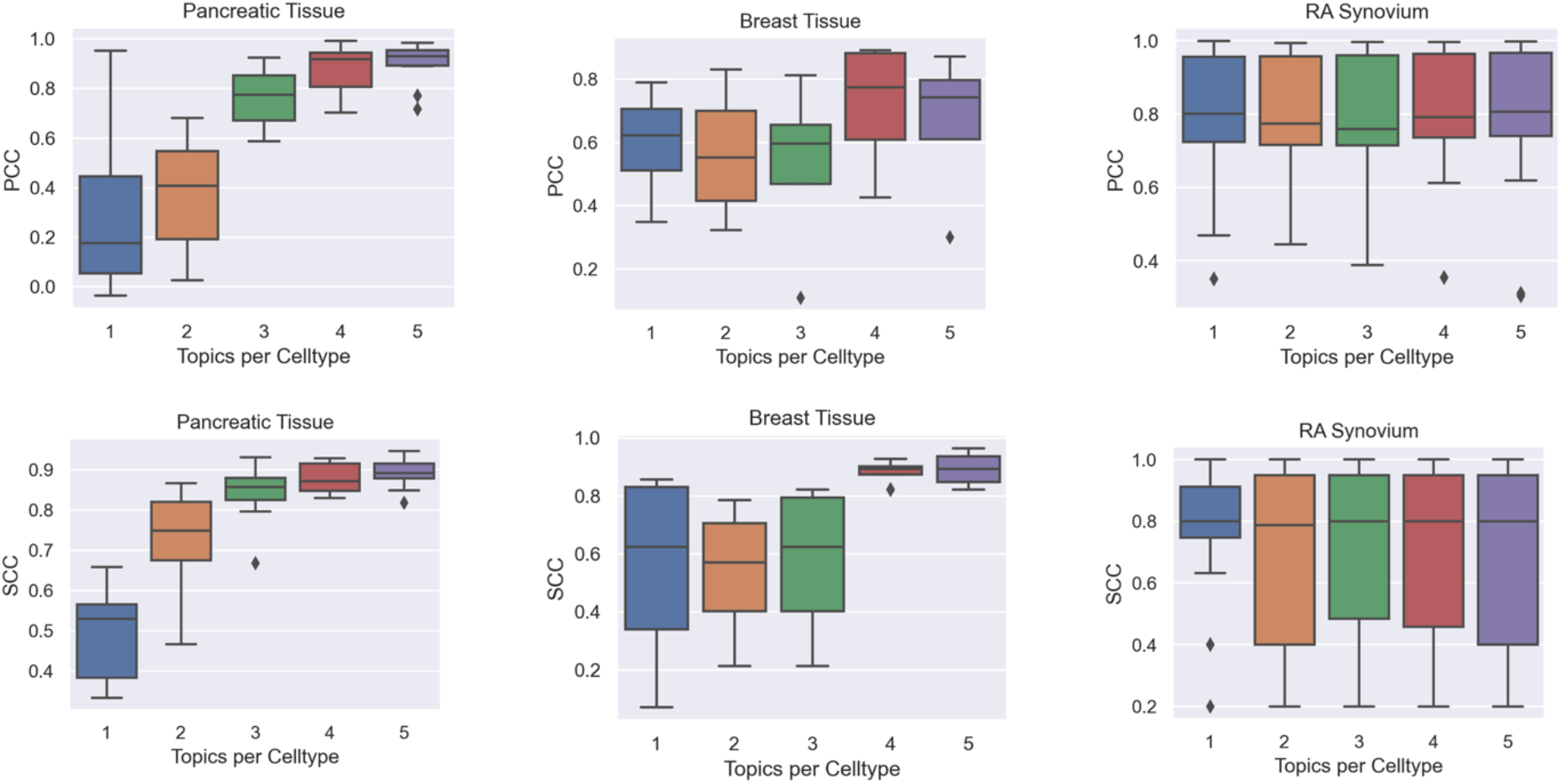
Deconvolution accuracy for different topics per cell type. Each GTM-decon model was trained by modelling a cell type using K number of topics, with K varying from 1 to 5, using all genes. We evaluated the deconvolution accuracy of these models on simulated bulk data from three datasets: Pancreatic (Segerstolpe), Breast (Normal), and RA Synovium. We simulated the bulk data by summing up counts per gene for all cells per individual. For unbiased evaluation, we adopted a leave-one-out cross-validation (LOOCV) design. Each model was trained on N-1 individuals and used to deconvolve the simulated bulk transcriptome from the held-out individual for validation. The inferred cell proportions by each model were evaluated by Pearson correlation coefficient (PCC) (top row) and Spearman correlation coefficient (PCC) (bottom row). The box and the whiskers in each boxplot indicate the 25%-75% quartile and min-max of the evaluation scores over the LOOCV individuals, respectively.

**Figure S9:**
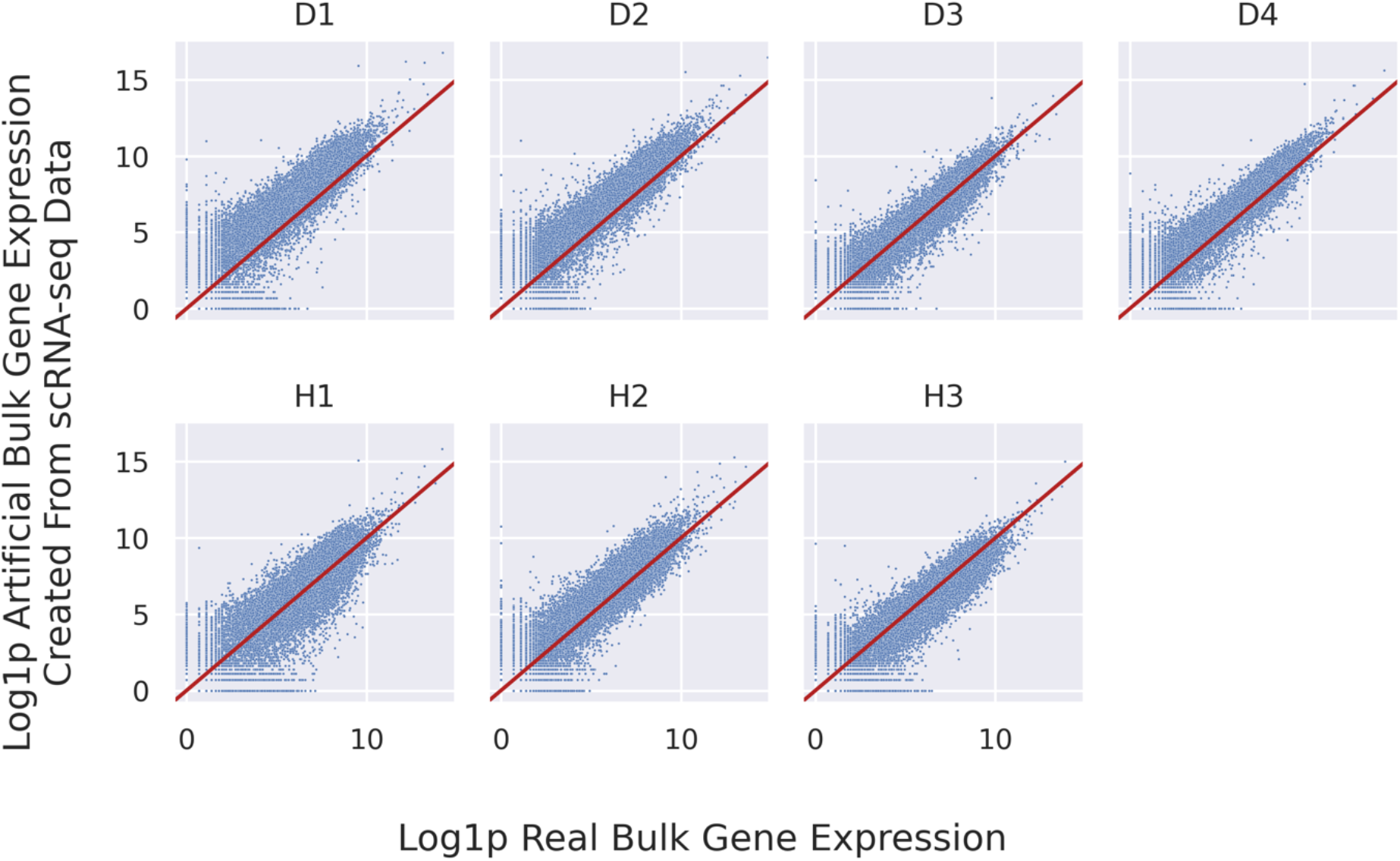
Correlation between real bulk gene expression and log1p artificial bulk gene expression. The scatter plots show the correlation between log1p normalized values of real bulk gene expression (x-axis) versus log1p normalized values of artificially constructed bulk gene expression values (y-axis) for each gene. The plots are shown for each of the individuals in the Segerstolpe datasets E-MTAB5061 for scRNA-seq, and E-MTAB5060 for bulk RNA-seq, respectively. H1, H2, H3 denote samples from healthy individuals and D1, D2, D3, D4 denote samples from diabetic individuals.

**Figure S10:**
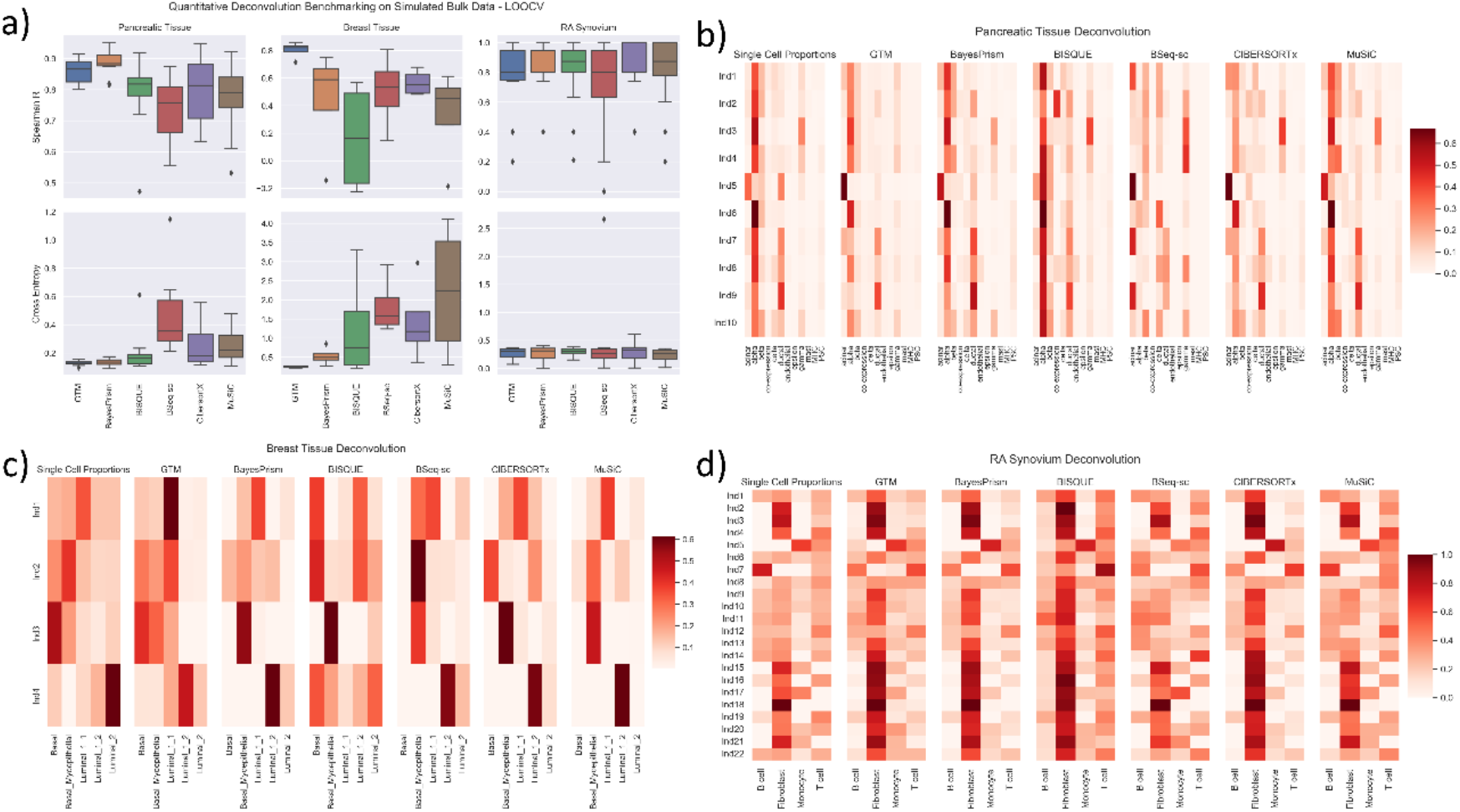
Evaluation of deconvolution accuracy of simulated bulk data. a) Evaluation of cell-type deconvolution accuracy. We simulated the bulk data by summing up counts per gene for all cells per individual. For unbiased evaluation, we adopted a leave-one-out cross-validation (LOOCV) design. Each deconvolution method was trained on N-1 individuals and deconvolved the bulk transcriptome simulated using the scRNA-seq data from the held-out individual for validation. The inferred cell proportions by each method were evaluated by Spearman correlation and Cross Entropy. The box and the whiskers in each boxplot indicate the 25%-75% quartile and min-max of the evaluation scores over the LOOCV individuals, respectively. b-d) Ground truth and Inferred cell-type proportions by each method on Pancreatic, Breast and RA Synovium Tissues simulated bulk data, respectively. In each heatmap, the rows are individuals, and the columns are cell types. The color intensity is proportional to the inferred cell-type proportions.

**Figure S11:**
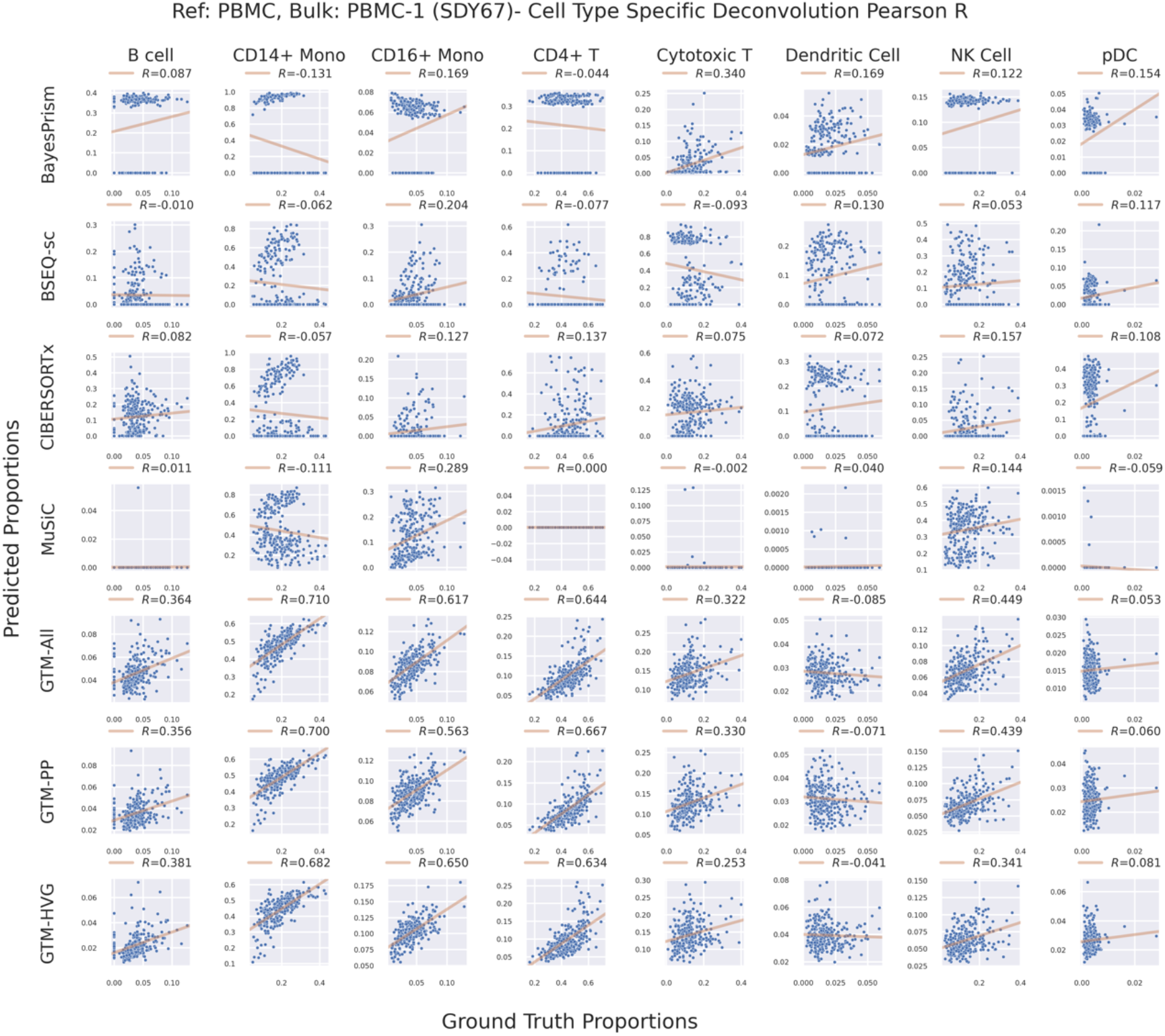
Scatterplot of inferred cell-type proportion and ground-truth true proportion of real bulk data from PBMC-1 (SDY67 in Table S2) based on a separate scRNA-seq reference of PBMC2.

**Figure S12:**
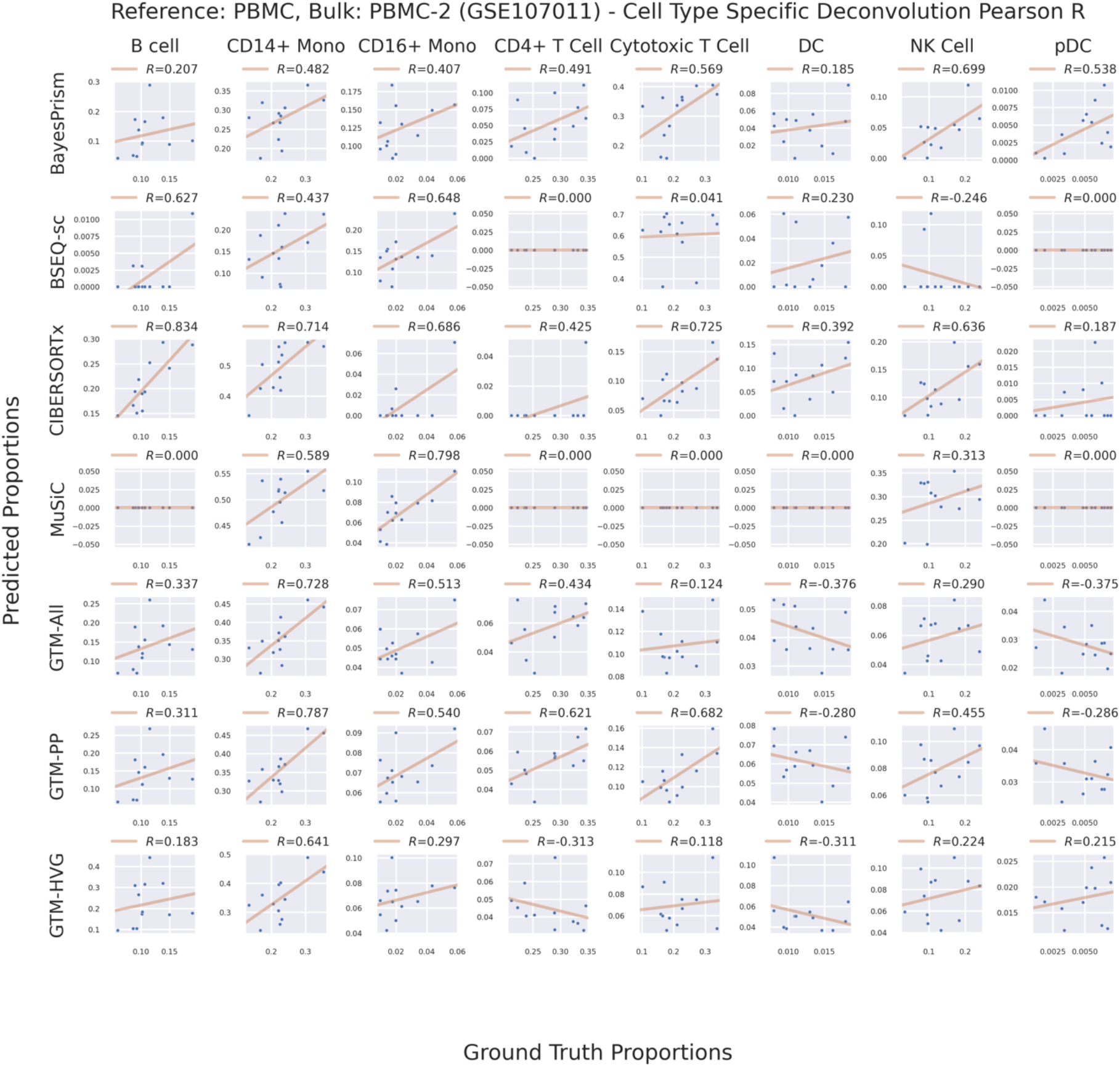
Scatterplot of inferred cell-type proportion and ground-truth true proportion of real bulk data from PBMC-2 (GSE107011 in Table S2) based on a separate scRNA-seq reference of PBMC2 (Table S1).

**Figure S13:**
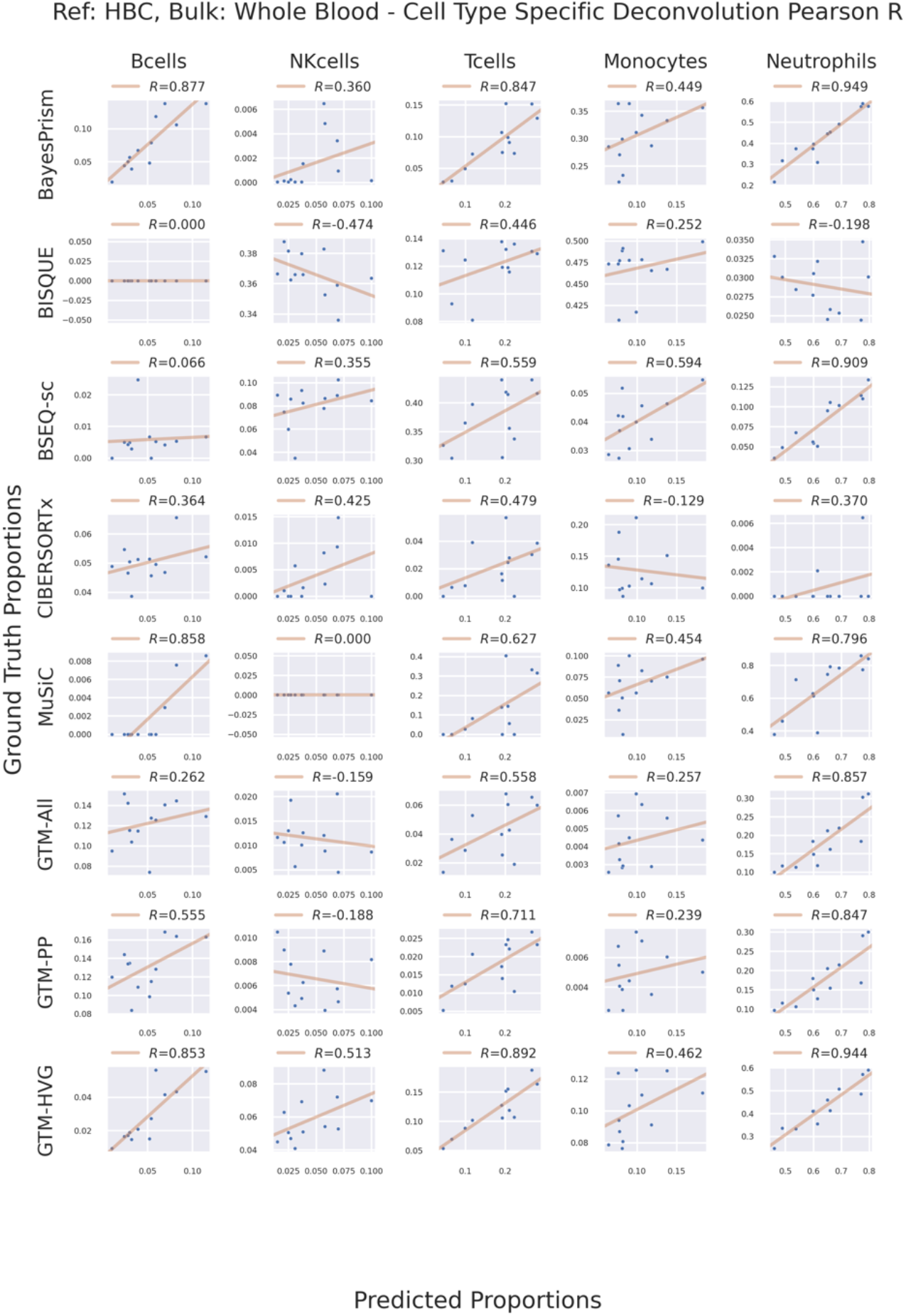
Scatterplot of inferred cell-type proportion and ground-truth true proportion of real bulk data from Whole Blood (WB in Table S2) based on a separate scRNA-seq reference of Human Blood Cell (HBC in Table S1).

**Figure S14:**
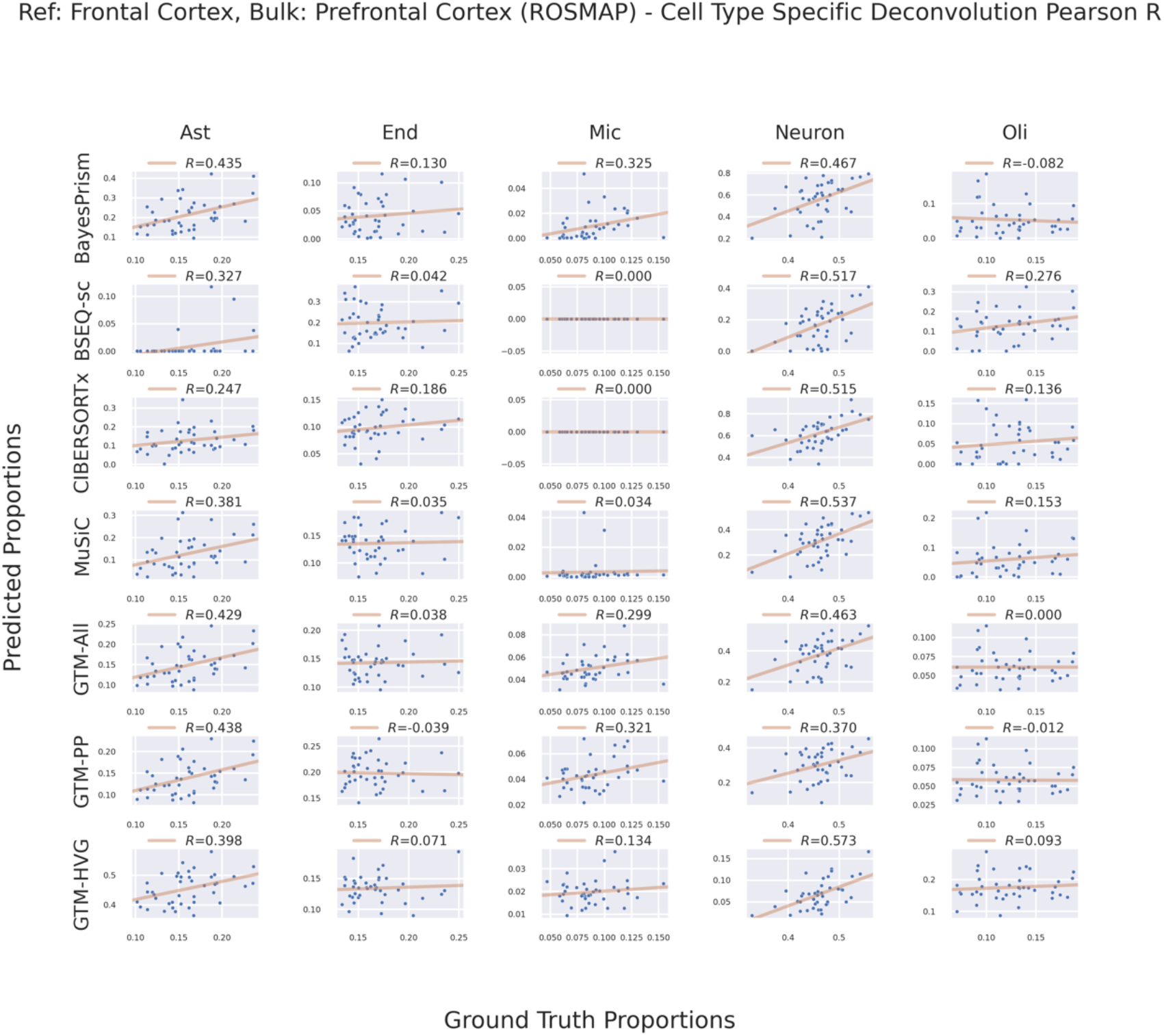
Scatterplot of inferred cell-type proportion and ground-truth true proportion of real bulk data from Prefrontal cortex (ROSMAP in Table S2) based on a separate scRNA-seq reference of Frontal Cortex (Table S1).

**Figure S15:**
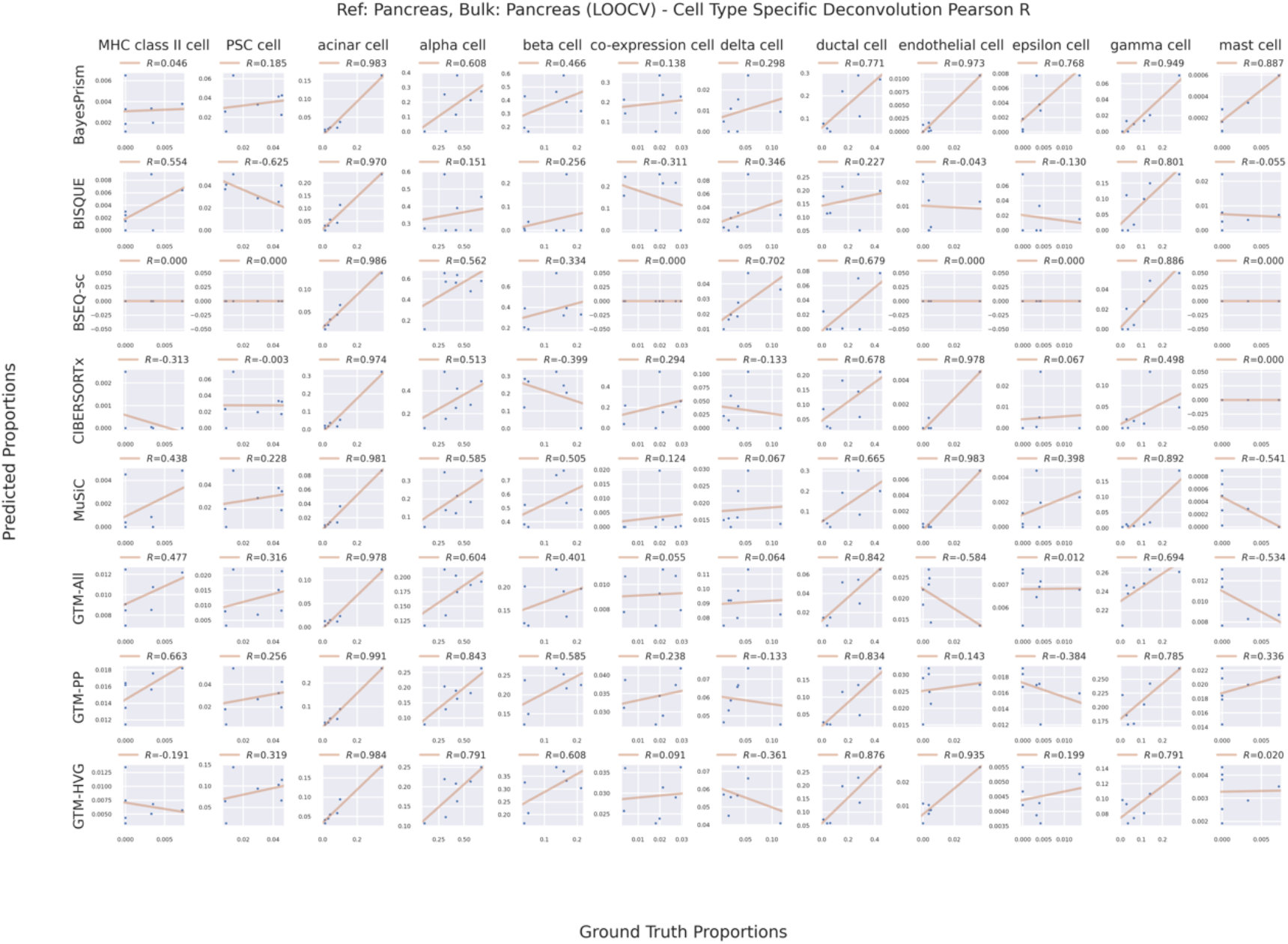
Scatterplot of inferred cell-type proportion from real bulk data from Pancreas (Segerstolpe in Table S2) and estimated proportion from the paired scRNA-seq data in the same held-out subject. The scRNA-seq reference used to train each model came from the other subjects (Segerstolpe in Table S1).

**Figure S16:**
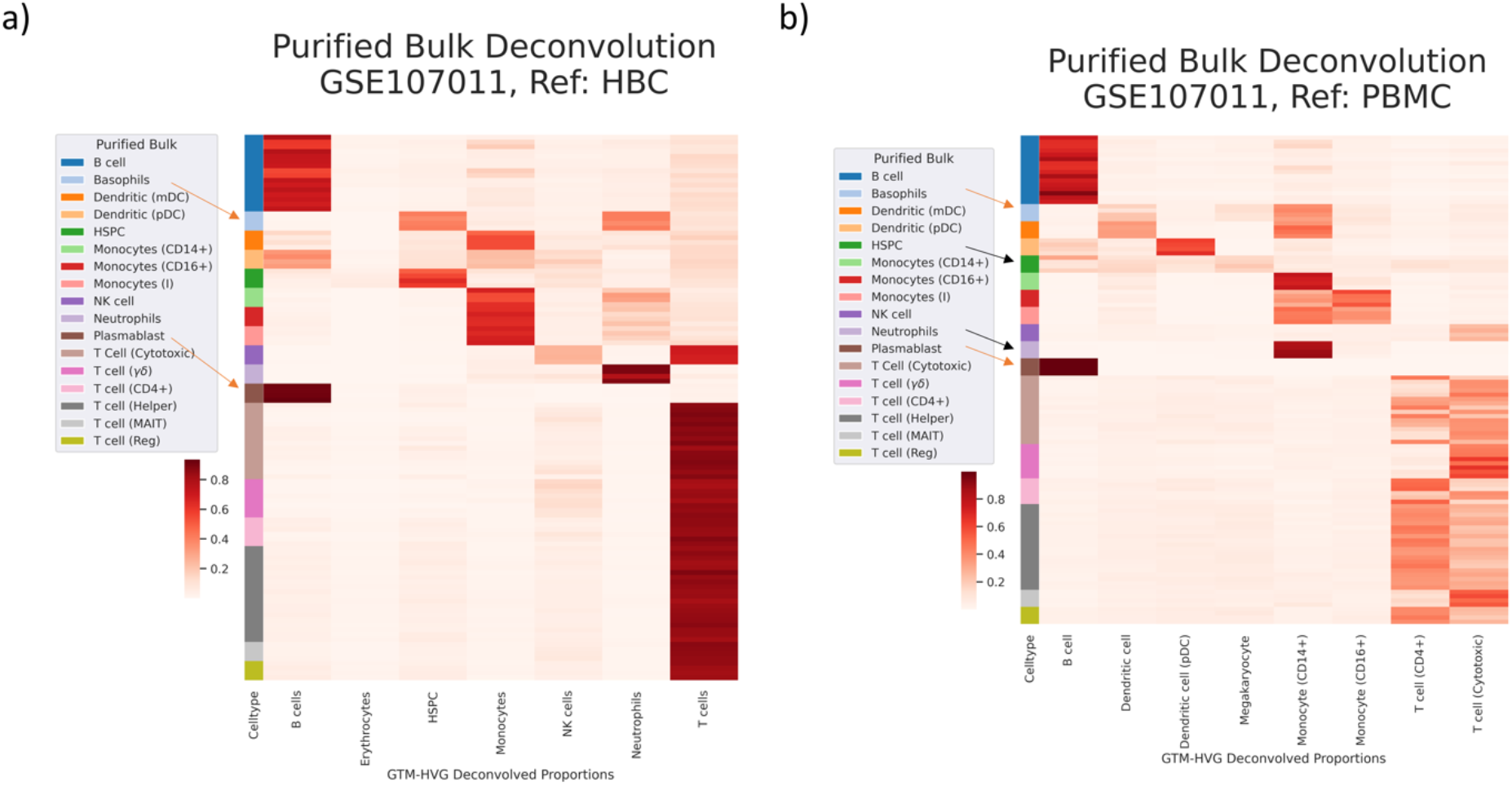
Inferred cell-type proportion over purified bulk samples of immune cell types (GEO access: GSE107011). The CTS topics were inferred separately from two distinct scRNA-seq references with incomplete cell types with respect to the target bulk samples. We also experimented using either all genes or highly variable genes (HVG). The heatmaps show a qualitative comparison of the combinations of the reference and gene selection, where the rows are the bulk samples with color legend indicating their cell types and the columns are the cell types from the scRNA-seq HBC reference (left panel) and PBMC2 reference (right panel).

**Figure S17:**
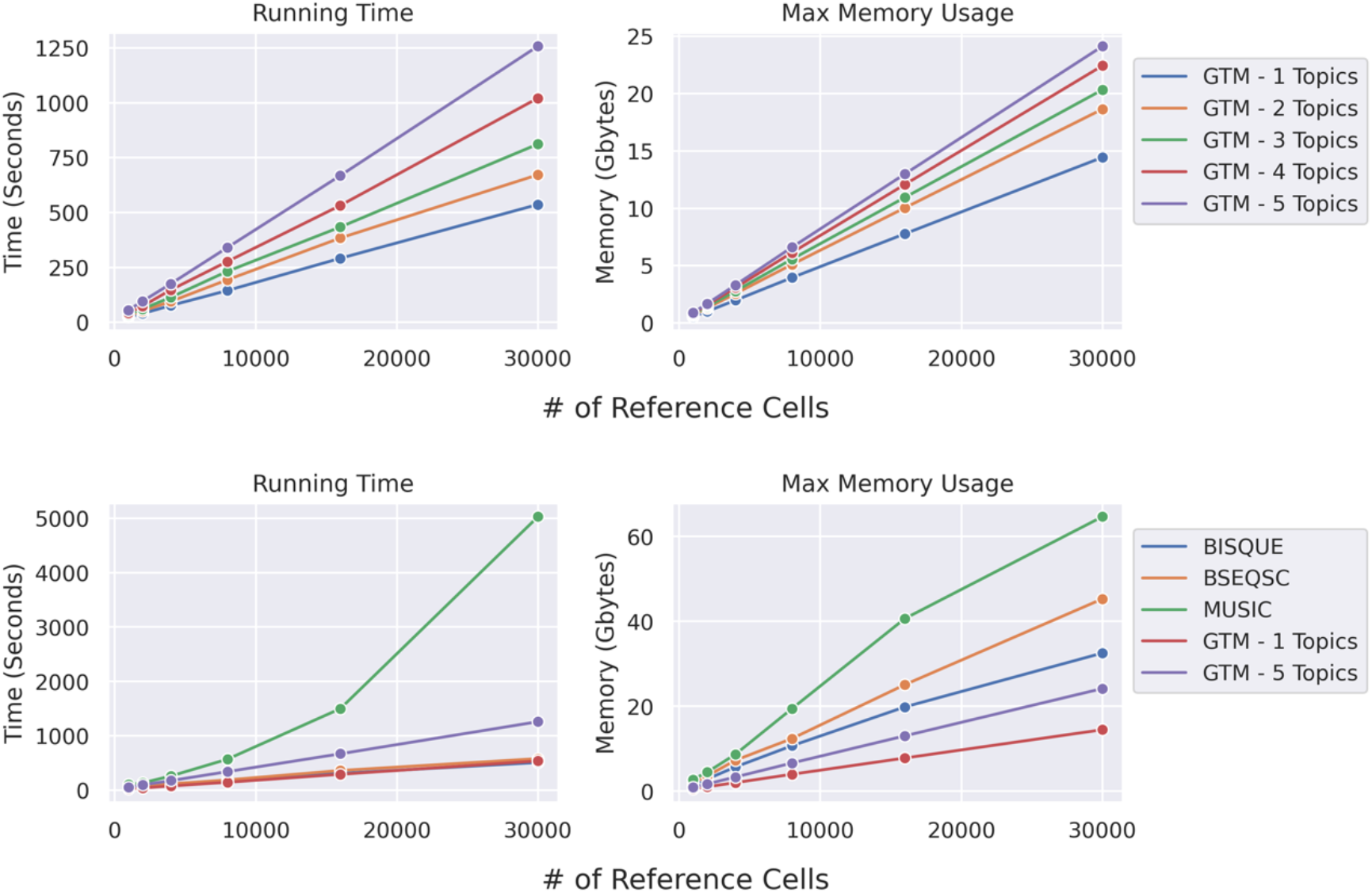
Benchmark of time and memory usage of the deconvolution methods. We benchmarked the time and memory behavior or GTM-decon as a function of both input reference size and number of topics per cell-type. We trained on 1e3, 2e3, 4e3, 8e3, 1.6e4, 3e4 cells, with 33694 genes and deconvolved the same purified bulk PBMC data (GSE64655). BISQUE, BSEQ-sc, and MuSiC were all run with their default settings. All methods were run on a server with 62Gb RAM, 29Gb swap, with 20 Intel Xeon CPU e5-2650 v4s at 2.20Ghz. GTM-decon time and memory cost increase with the number of topics, but not exponentially, and GTM-decon at 1 topic per cell type has the lowest max memory usage and comparable run times to BSEQ-sc and BISQUE, and GTM-decon with 5 topics per cell-type still achieves much higher speed than MuSiC, the slowest model.

**Figure S18:**
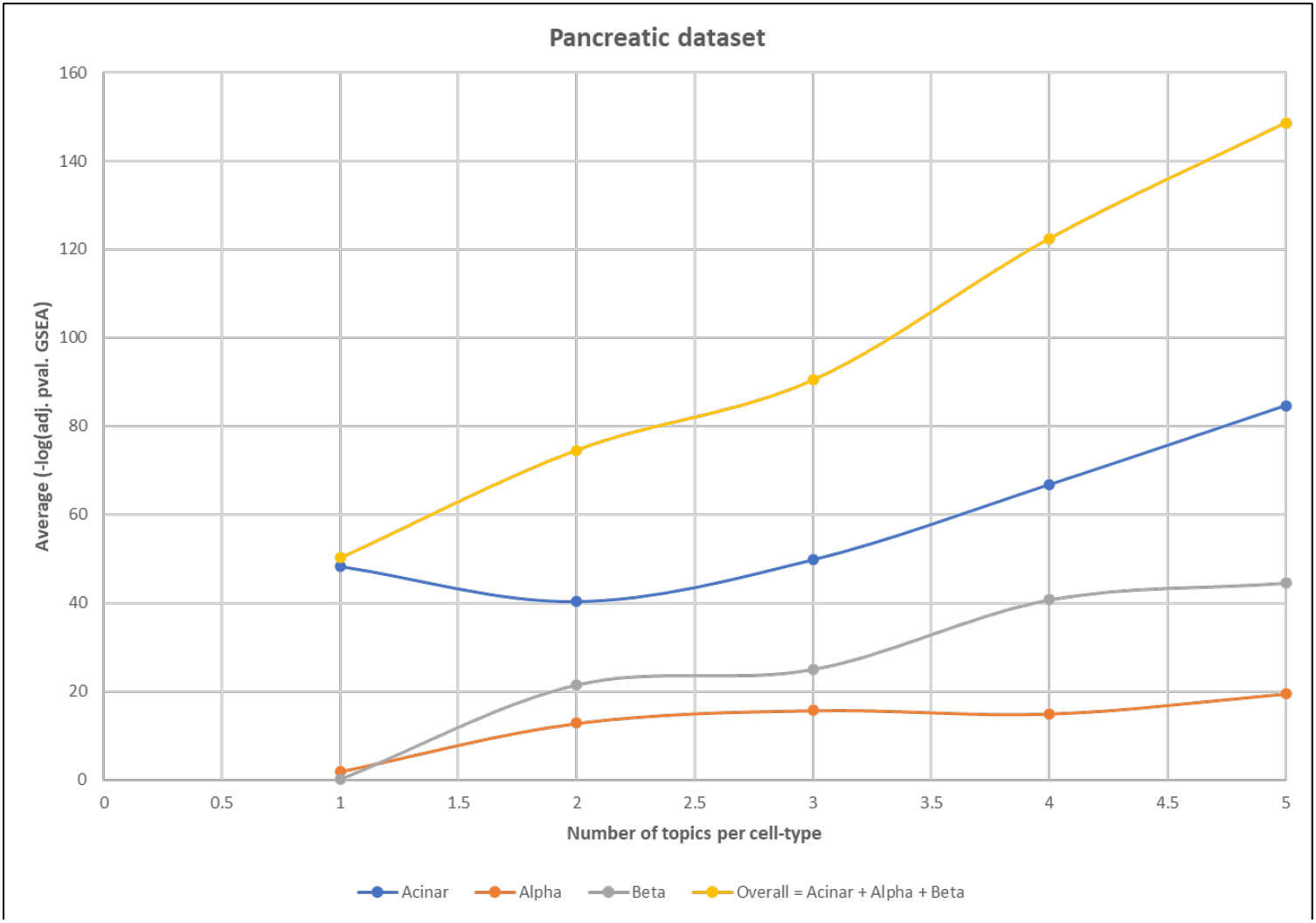
Evaluating the number of topics per cell-type based on GSEA of known marker genes. Cell-type-specific topics for Acinar, Alpha, and Beta cell types from Pancreatic Islet dataset from Segerstolpe et al. were evaluated based on whether the top genes are enriched for the known marker genes (from CellMarkerDB) under that cell type using gene set enrichment analysis. Each model was trained with varying numbers of topics per cell type (from 1 to 5). The plot shows the average (-log10 adjusted p-value) for the cell type of interest for the different topics. An overall value is calculated by summing up the values for the three cell types.

**Figure S19:**
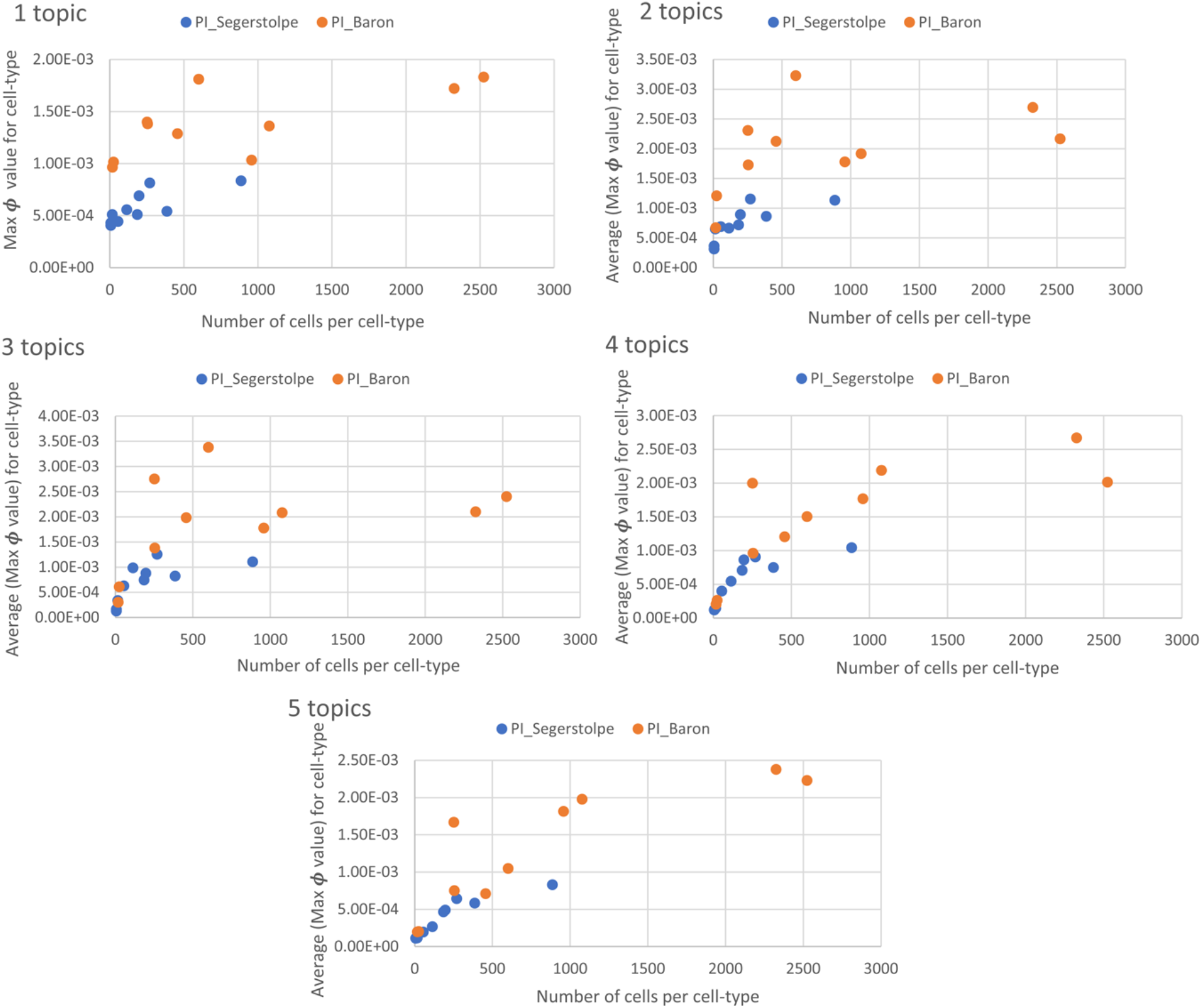
Evaluating topic confidence in terms of number of cells per cell type. We evaluated topic confidence by correlating the topic confidence scores with the number of cells for each cell type available for training. The topic confidence scores were calculated by the first taking the maximum probability from the G-by-1 column vector #_*k*_ for each cell type *k* ∈ &, …, *K* and then averaging over all K cell types. The resulting topic confidence scores were plotted as a function of number of cells per cell type. Each panel correspond to the number of topics per cell type (varying from 1 to 5) for two pancreatic datasets, Segerstolpe and Baron, generated using different approaches, Smart-seq2 and Drop-seq, respectively.

**Figure S20:**
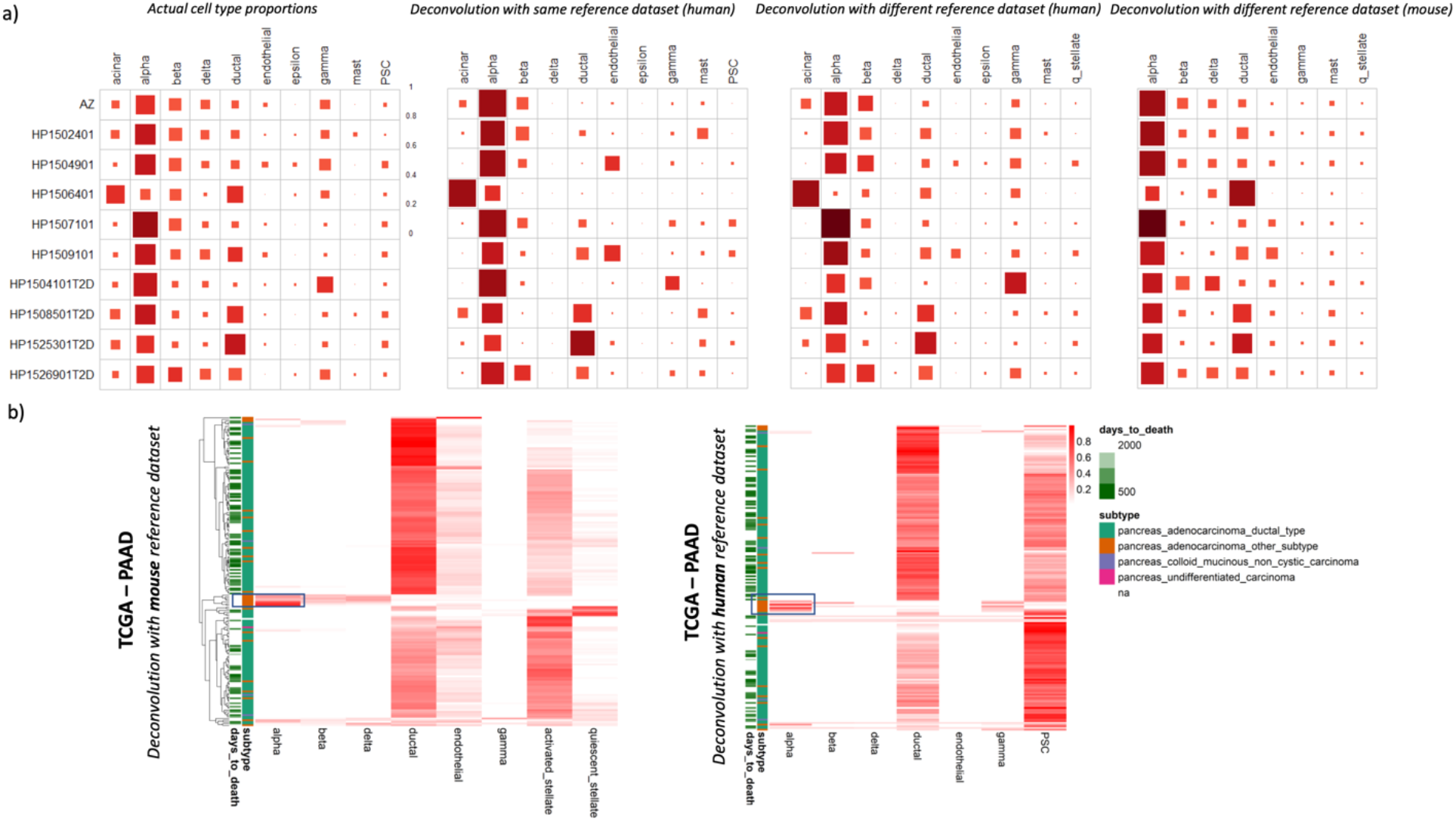
Cross-reference and cross-species deconvolution. a) Deconvolution of PI-Segerstolpe human pancreas samples using different scRNA-seq reference data. The four Hinton plots from left to right display the ground truth cell-type proportions, the deconvolved proportions using the same reference dataset as target (i.e., Segerstolpe pancreas islet dataset), the deconvolved proportions using Baron pancreas islet scRNA-seq data as the reference, the deconvolved proportions using mouse pancreas scRNA-seq data as the reference. The rows are subjects and columns are cell types. b) Deconvolution of TCGA-PAAD dataset using human and mouse pancreatic datasets as reference. The samples (rows) were clustered by agglomerative clustering based on the GTM-decon model trained on the mouse reference (i.e., left heatmap). The same row order was applied to the heatmap based on human reference on the right. PSC in human dataset represents pancreatic stellate cells, which are represented as two different subtypes in mouse dataset. Four clinical phenotypes were shown as color bar on the left of each heatmap. Highlighted region for the alpha cell type was described in the main text.

**Figure S21:**
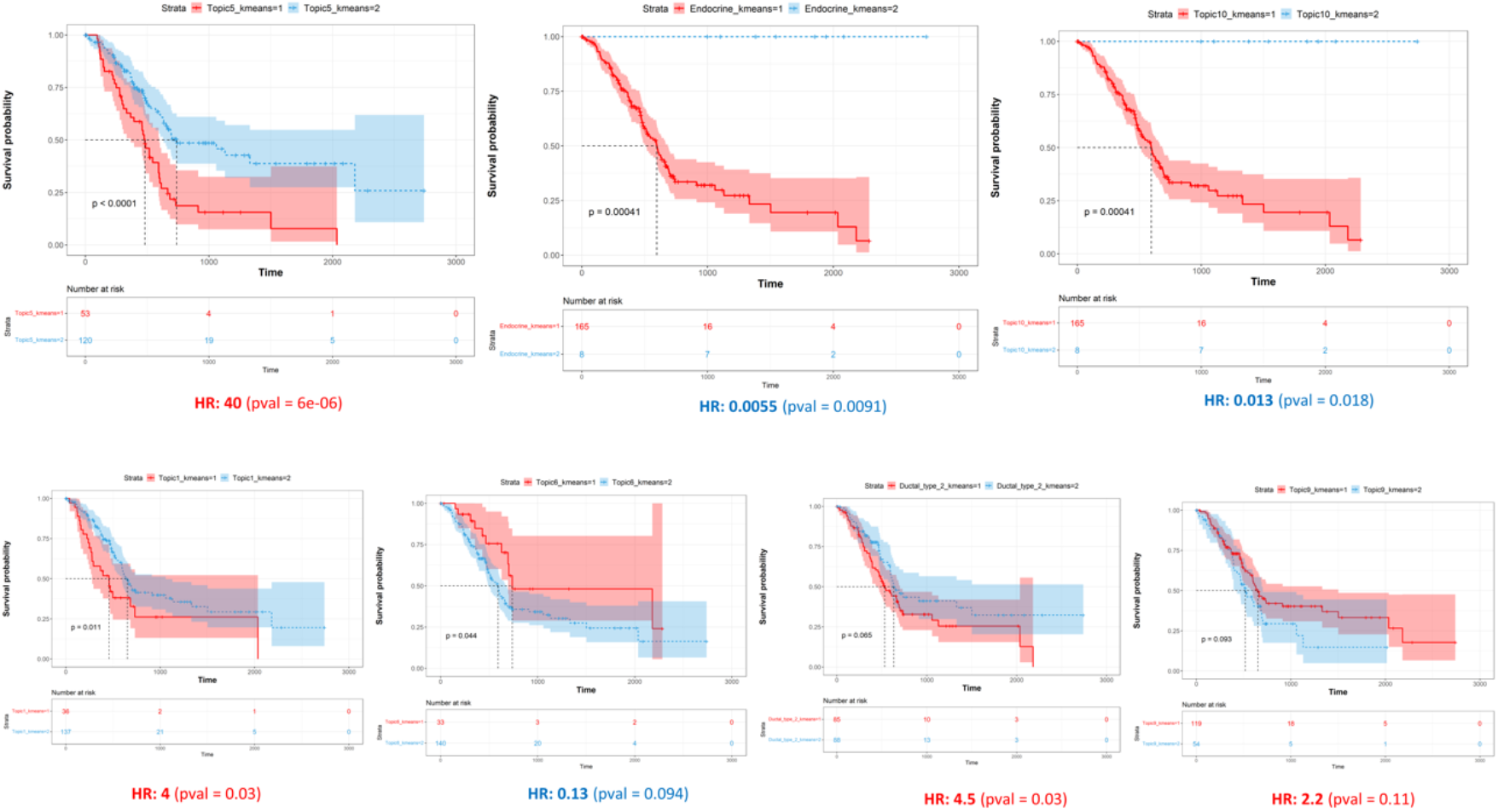
Pancreatic cancer survival analysis of CTS and *de novo* bulkRS topics. To explore the marginal effect of individual cell type proportions on survival, we performed Kaplan Meier analysis by separating patients into two groups based on K means clustering. This analysis was performed on both CTS and *de novo* topics. Figure 4c lists the topics with the most significant differences (p-value < 0.1; log-rank test). The Kaplan-Meier curves for all those topics are shown here. The curve and shaded area represent the mean and standard deviation of the cell type proportions in the two groups, respectively. The number of subjects for each cluster were indicated below each Kaplan-Meier plot. The hazard ratios are shown at the end, with values < 0 indicating good outcomes, and values >0 suggesting poor outcomes.

**Figure S22:**
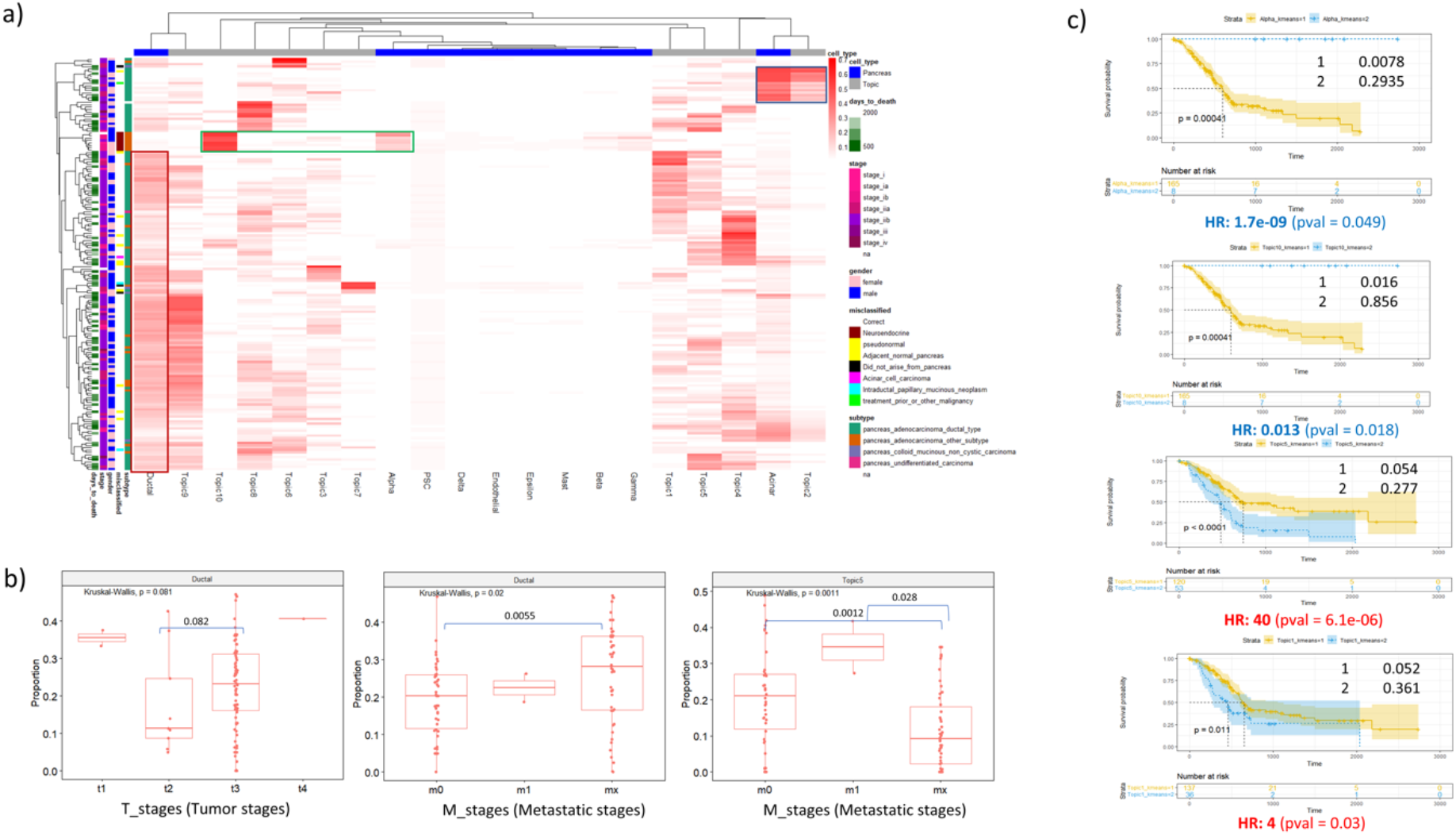
Deconvolution of bulk RNA-seq samples for pancreatic cancer from TCGA-PAAD using healthy pancreatic scRNA-seq. a) Inferred cell-type proportions of TCGA-PAAD tumor samples. GTM-decon was trained an scRNA-seq dataset from individuals with pancreatic cancer. The trained GTM-decon model was then used to deconvolve the 174 TCGA-PAAD bulk RNA-seq profiles. In addition, we also ran unguided topic model (i.e., LDA) on the TCGA-PAAD bulk RNA-seq profiles directly to detect *de novo* topics that are not present in scRNA-seq reference. The above heatmap visualizes the combined deconvolution results based on the 10 pancreatic cell types, and 10 *de novo* topics (i.e., columns). Each of the 174 rows represents a subject. Five types of demographic or clinical phenotypes were shown in the legend to aid result interpretation. These include days to death, cancer stage, sex, misclassified status, and cancer subtype. The regions in highlighted boxes are described in the main text. c) Survival analysis using inferred cell-type proportions. The 174 subjects were divided into two groups based on K*-means cluster with K* set to 2 (not to be confused with the K cell types or topics). Kaplan-Meier curves for cell types and the *de novo* topics that resulted in significant differences in terms of their hazard ratios were displayed. The curve and shaded area represent the mean and standard deviation of the cell-type proportions in the two groups, respectively. The number of subjects for each cluster were indicated below the Kaplan-Meier plot.

**Figure S23.**
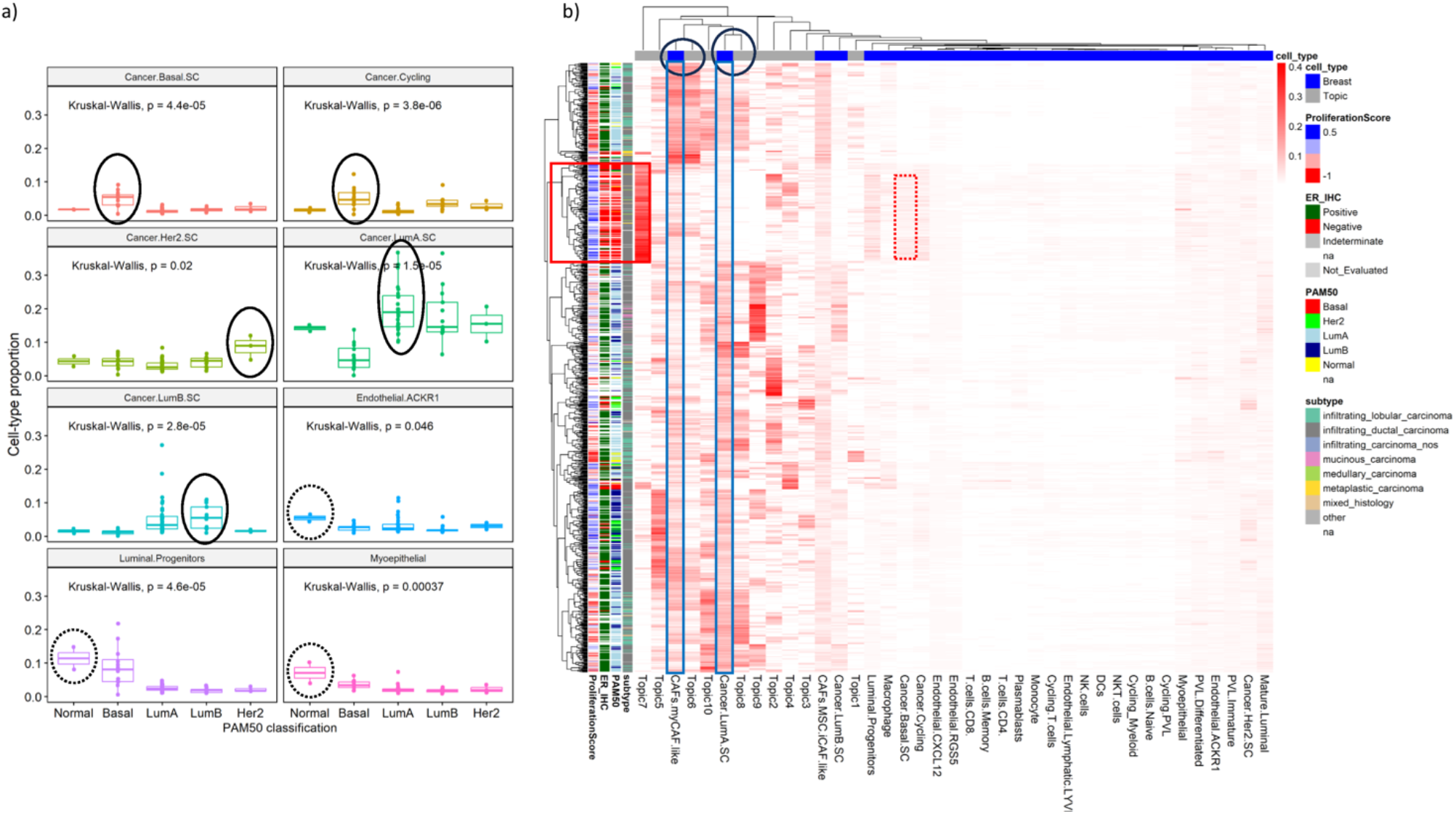
Deconvolving breast cancer bulk transcriptome based on single-cell transcriptomes from breast tumors. a) Inferred cell-type proportions matched the cancer subtypes. We trained GTM-decon on the scRNA-seq breast cancer dataset using the 8 cell types as a guide. Each boxplot shows the distribution of the cell-type proportion for the samples separated by the cancer subtypes as defined by the PAM50 system. P-values based on the Kruskal-Wallis were shown. The subtype that exhibits the highest cell-type proportion in each panel was circled. b) Detailed visualization of the deconvolved proportions. The consolidated heatmap illustrates the cell type and *de novo* topic proportions from GTM-decon and unguided LDA separately trained on the scRNA-seq breast cancer data and TCGA-BRCA data, respectively. These two sets of topics were colored in grey and blue, respectively. The rows represent the 1212 tumor samples and the columns represent the cell types and *de novo* topics. The clinical phenotypes were annotated in the left color bar to help interpret the topics. Abbreviations of cell-types: CAFs – cancer associated fibroblasts; myCAF – myofibroblast; iCAF – inflammatory CAF; DCs – dendritic cells; NK – natural killer; NKT – natural killer T cells; PVL – perivascular-like.

**Figure S24.**
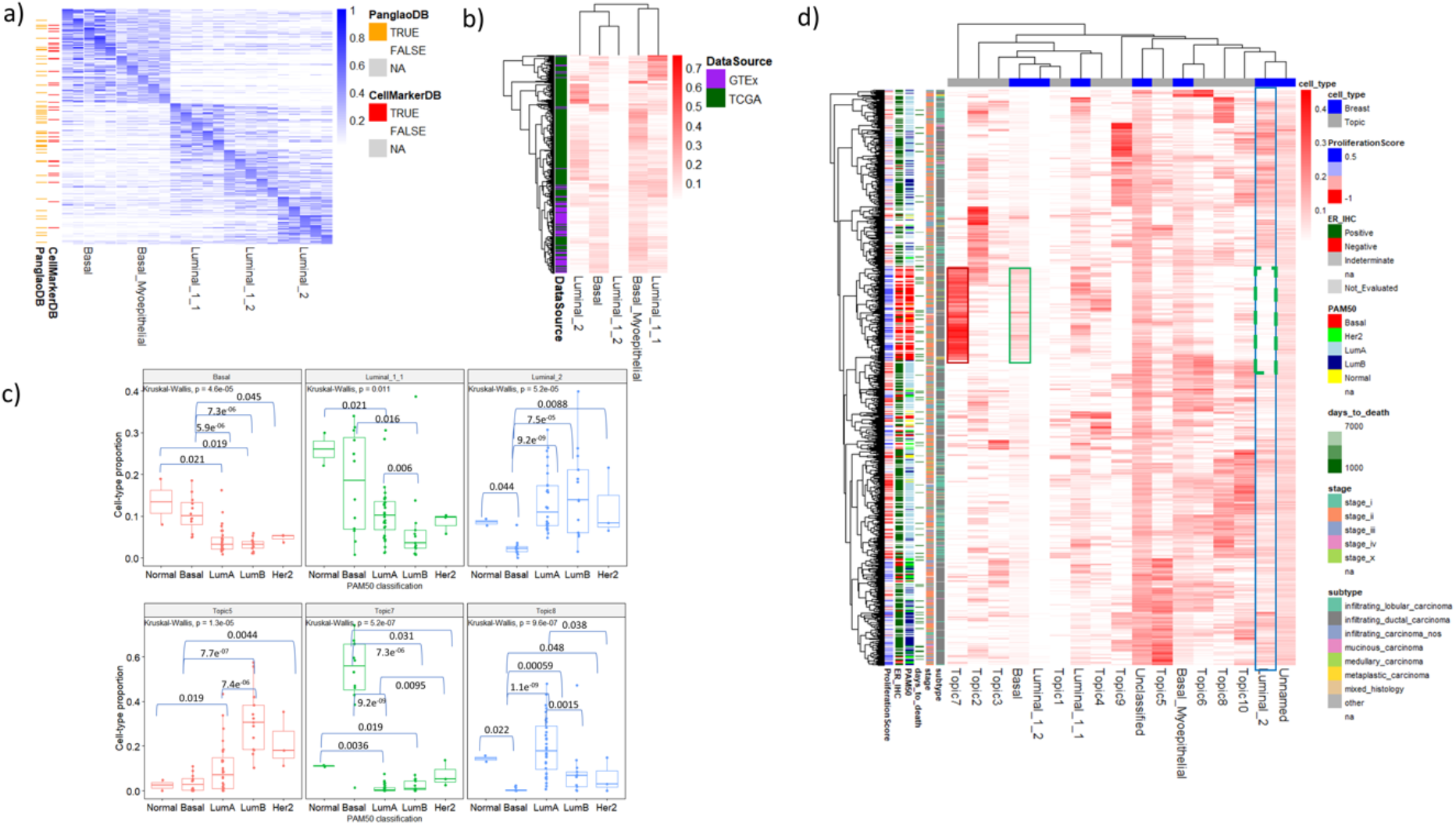
Cell-type-specific inference and deconvolution of breast transcriptomes using HVG gene set from scRNA-seq data of healthy breast tissue. a) Gene signatures of cell-type-specific topics in breast tissue. We trained GTM-decon on a scRNA-seq reference of normal breast tissue [11] using 5 topics per cell type, on the set of highly variable genes. Same as Figure 3, the heatmap visualizes the top 20 genes per topic for the 5 main cell types in normal breast tissue (i.e., 25 topics in total for the 5 cell types). The heatmap intensity is proportional to the gene topic probabilities. Whenever available from CellMarkerDB and PanglaoDB, cell-type marker genes are indicated on the left. For the cell types, where marker genes are not available, “NA” were indicated on the left. b) Heatmap of deconvolved cell-type proportions for the main cell types for GTEx (normal samples) and TCGA-BRCA (cancer samples). The rows represent samples and columns represent the reference cell types. The heat intensity is proportional to the inferred cell-type proportions. Agglomerative clustering was applied to both the rows and columns to cluster samples and cell types respectively using Euclidean distance metric and complete linkage clustering. c) Comparison of inferred cell type or *de novo* topic proportions across breast cancer types. Each boxplot displays the distribution of cell-type proportions or topic proportions (y-axis) for the breast tumor samples separated by their cancer classes based on Prediction Analysis of Microarray 50 (PAM50) (x-axis). In each panel, p-value based on Kruskal-Wallis test was indicated on the top left corner and p-values based on Wilcox Rank Sum test for the significant pairwise comparison between PAM50 classes were also indicated. d) Comprehensive visualization of deconvolution results for the 1212 breast tumor samples from TCGA. The deconvolution results were consolidated from the inferred topics of three topic models, namely GTM-decon trained on breast scRNA-seq reference (i.e., the same model from panel a), GTM-decon trained on immune scRNA-seq reference, and unguided 10-topic LDA directly trained on the TCGA-BRCA. The three sets of topics were indicated by blue, green, and grey colors, respectively on the top color bar. The row represents the 1212 tumor samples with 6 types of clinical phenotypes annotation on the left color bar to aid result interpretation. The regions in the highlighted boxes were discussed in the main text. Deconvolution results using all genes were shown in **Figure S25**.

**Figure S25:**
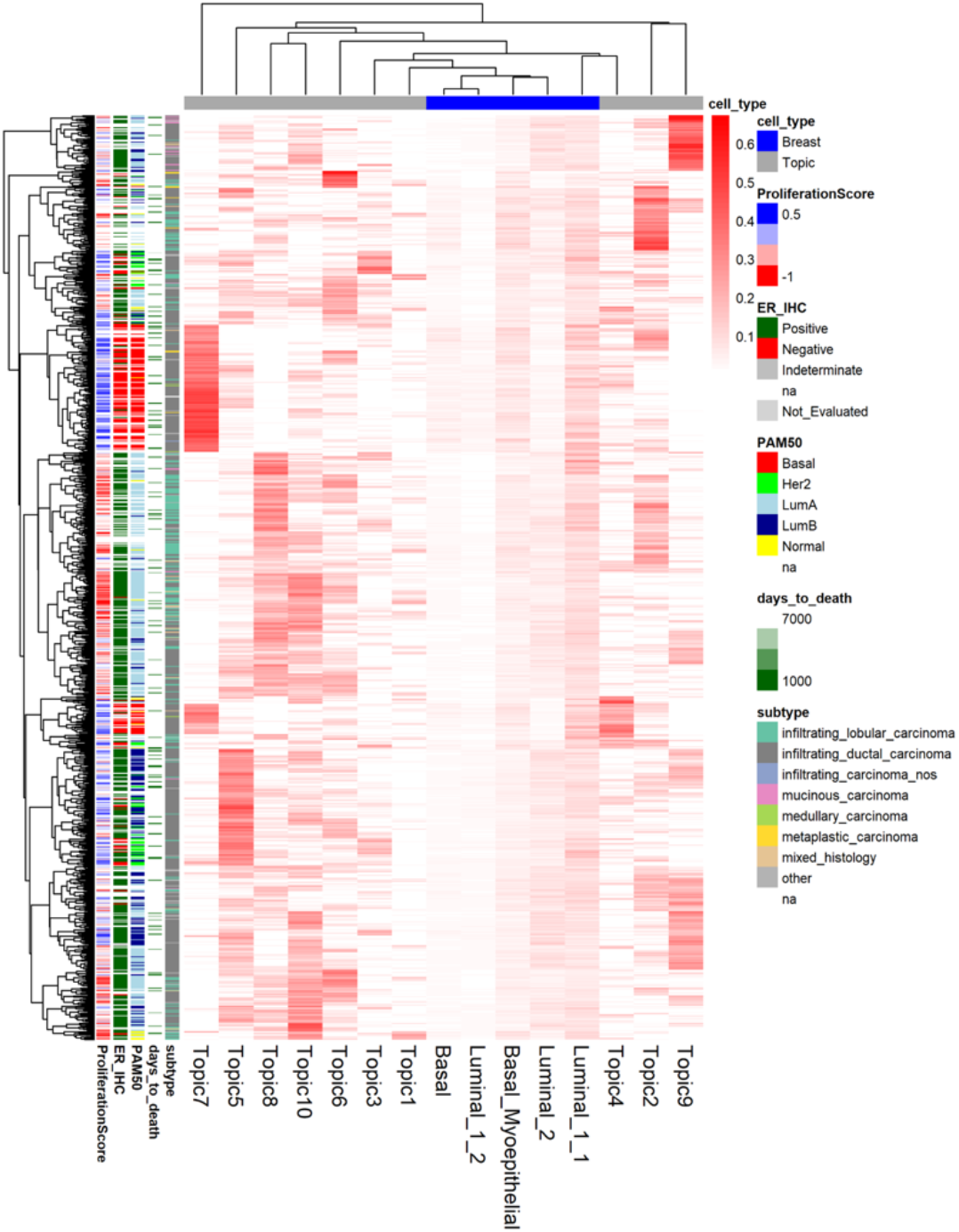
Deconvolution of breast cancer transcriptomes from TCGA using *all genes*. Visualization of deconvolution results for the 1212 breast tumor samples from TCGA. The deconvolution results were consolidated from the inferred topics of three topic models, namely GTM-decon trained on breast scRNA-seq reference using all genes with 5 topics per cell type, GTM-decon trained on immune scRNA-seq reference using all genes with 5 topics per cell type, and unguided 10-topic LDA directly trained on the TCGA-BRCA. The three sets of topics were indicated by blue, green, and grey colors, respectively on the top color bar. The row represents the 1212 tumor samples with 6 types of clinical phenotypes annotation on the left color bar to aid result interpretation.

**Figure S26:**
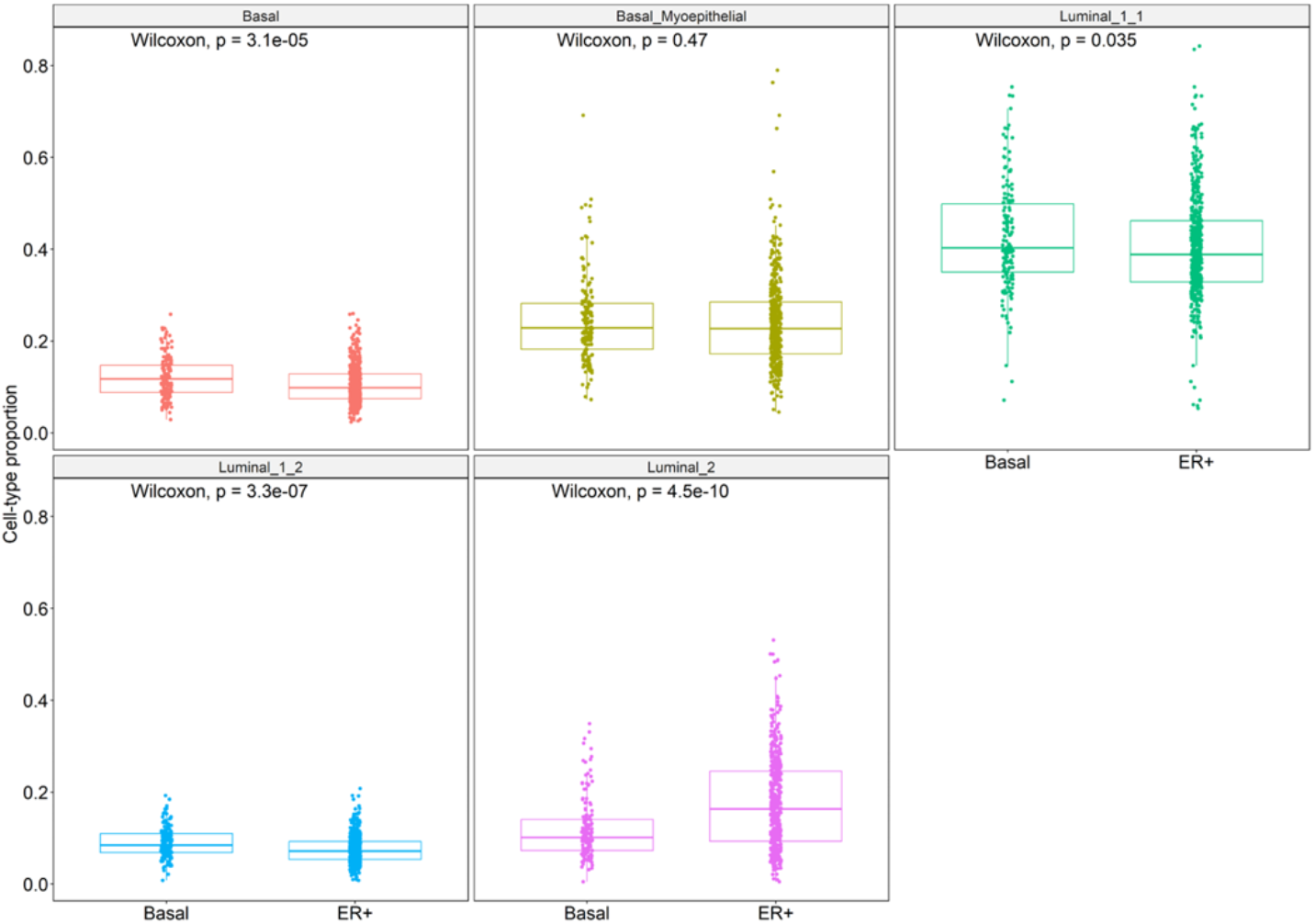
Differential analysis of cell-type proportions by breast cancer subtypes of Basal and ER+. The box plots show the distribution of inferred cell type proportions for the five cell types in terms of the breast cancer subtypes – Basal and ER+. The box and the whiskers in each boxplot indicate the 25%-75% quartile and min-max of the evaluation scores for each of the samples. Statistical significance calculated using the Wilcoxon test suggests that four of the cell types show significant differences in proportions across the subtypes, with Luminal_2 exhibiting the highest difference.

**Figure S27:**
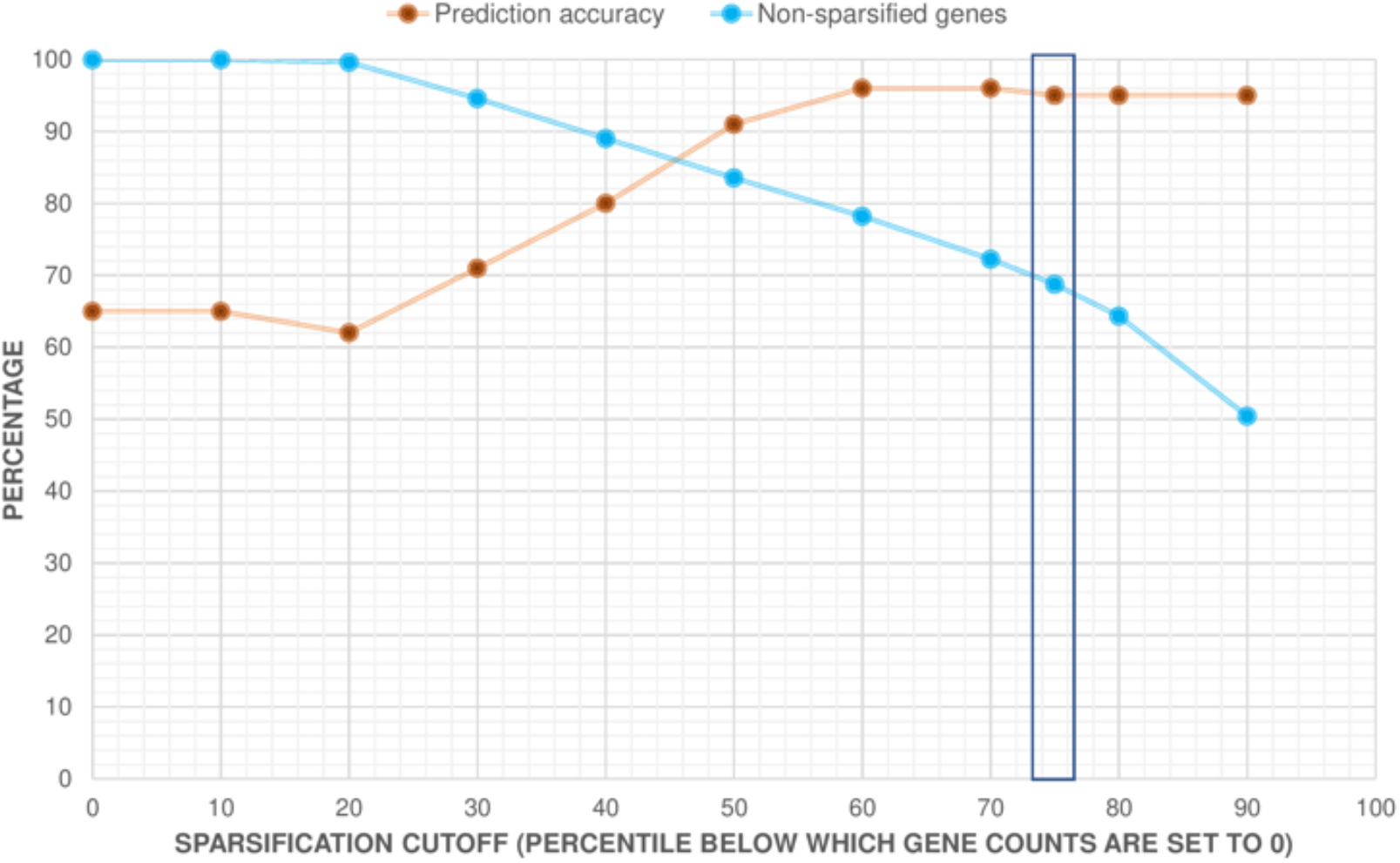
Effect of sparsification rate on prediction accuracy of ER+ versus Basal subtype and percentage of non-zero genes. In order to effectively capture phenotype signatures directly from the bulk RNA-seq data using our topic modeling approach, the TCGA-BRCA RNA-seq data were sparsified to varying degrees by setting the values below the “n-th” percentile to be zero. We varied the percentiles from 10 to 90, with the 75-th percentile representing the value chosen based on scRNA-seq datasets. The model performance was evaluated as the phenotype prediction accuracy comparing the phenotype-specific topic probabilities with the groundtruth ER+ and Basal on the held-out 20% subjects. The percentage of the non-zero genes (i.e. genes which have non-zero counts in at least one sample after the sparsification procedure) was also shown in the blue curve as a function of sparsification rate.

**Figure S28.**
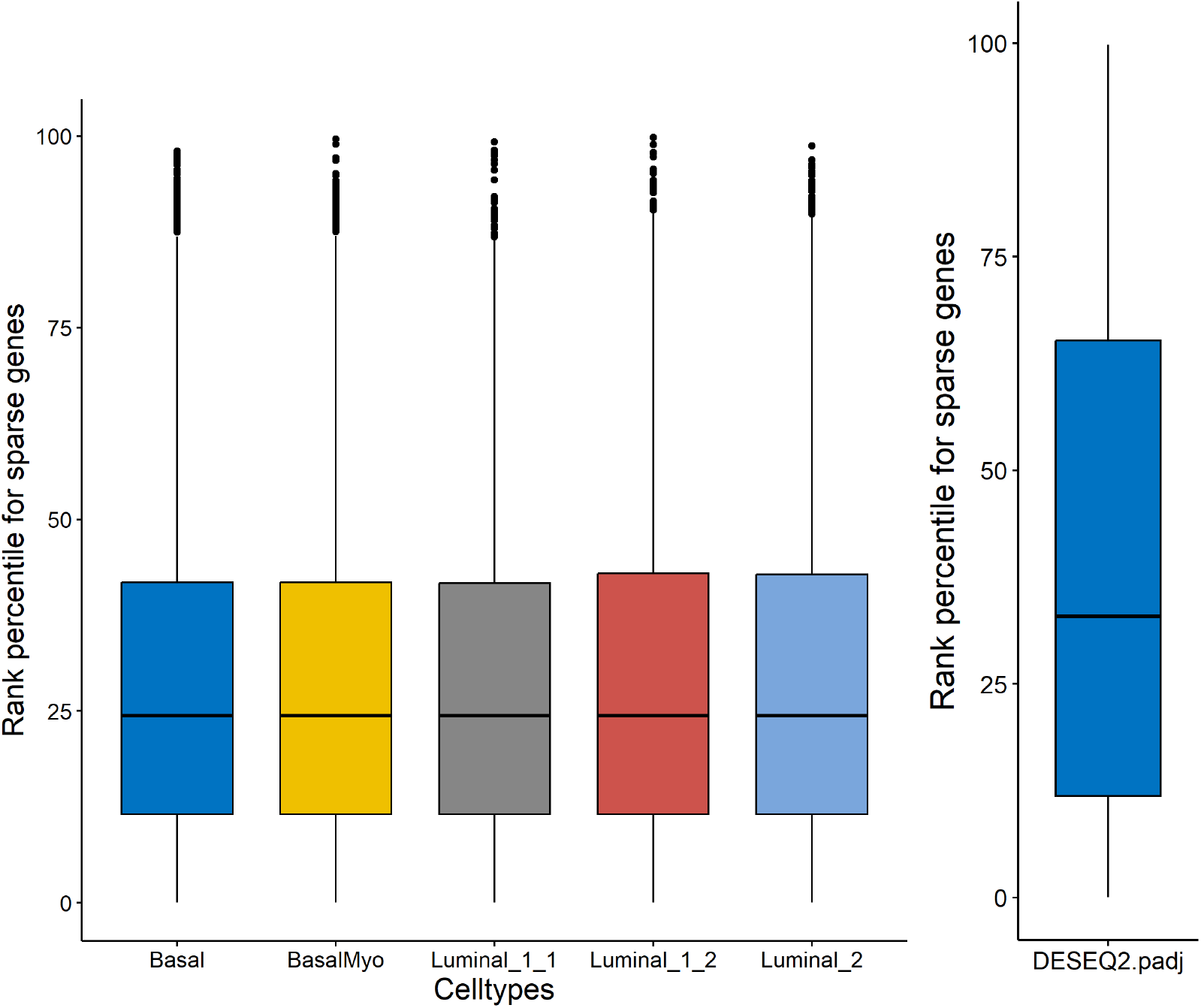
Marginal contribution of zero genes to the CTS topics and DE analysis. <u>Left panel</u>: the boxplots show the distribution of rank percentiles of the genes-by-CTS topic probabilities of the zero genes due to 75-percentile sparsification in the entire gene list for the 5 cell types from the scRNA-seq data of breast tissues. The gene with the highest probability for that cell type is set at 100-th percentile rank and the gene with the lowest probability at the 0-th percentile. <u>Right pane</u>l: The boxplot shows the distribution of the rank percentiles of adjusted p-values of DE genes identified by DESeq2 from whole bulk. The gene with the lowest p-value was set at 100-th percentile rank and the gene with the highest p-value at the 0-th percentile.

**Figure S29:**
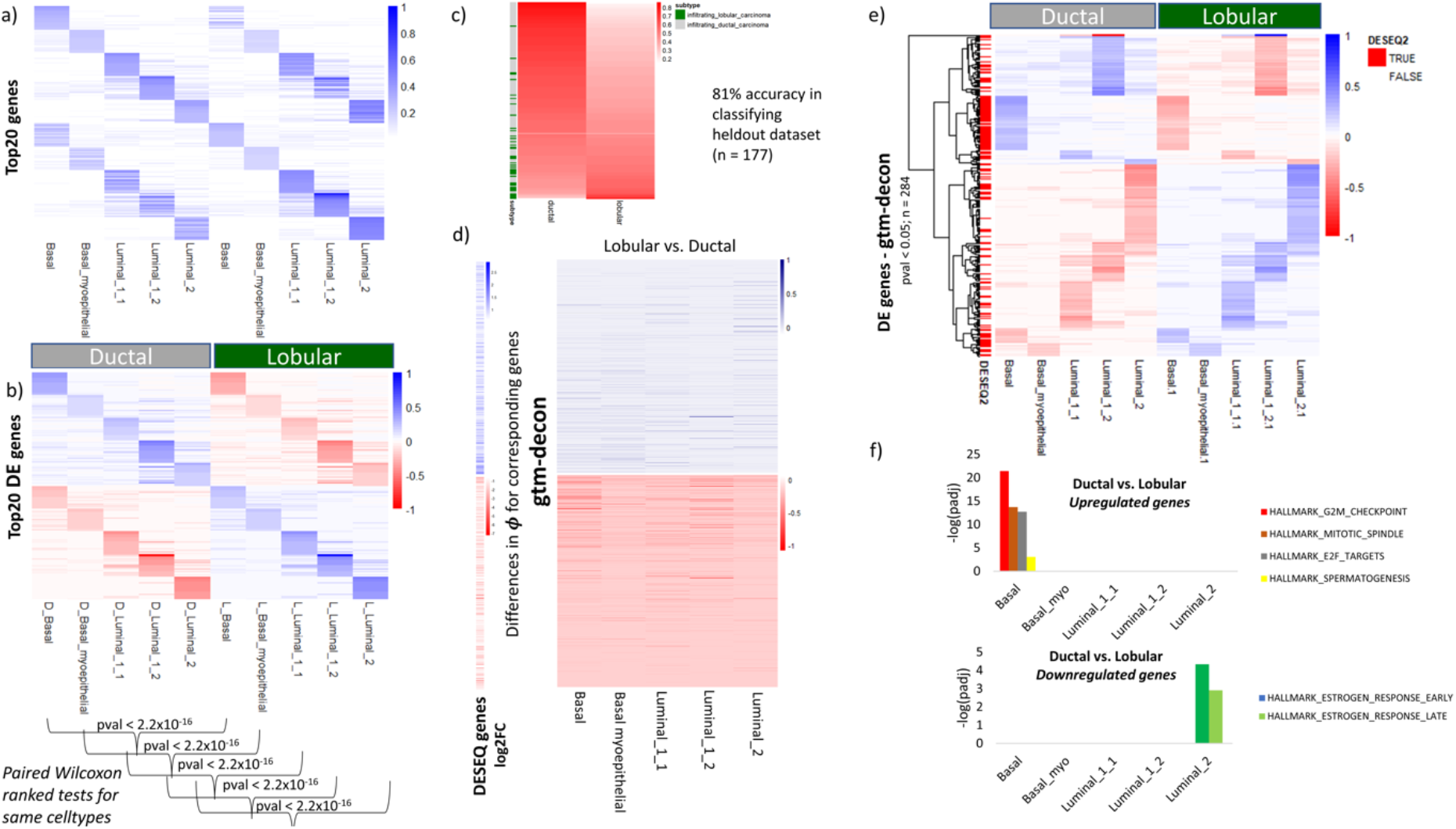
Identification of cell-type-specific differentially expressed genes from bulk RNA-seq data comparing between ductal and lobular breast carcinoma subtypes. a) Bulk RNA-seq samples of ductal and lobular subtypes of breast carcinoma from TCGA were analyzed using the same approach described in Fig. 6 to identify cell-type-specific differentially expressed (DE) genes corresponding to these two subtypes. a) Top cell type specific genes in each subtype. Visualization of top cell-type-specific gene signatures for Ductal and Lobular carcinoma in terms of gene-by-cell-type proportions for each subtype. b) Top DE genes in each subtype. Visualization of top predicted differentially expressed (DE) genes for each cell type between Ductal and Lobular carcinoma in terms of gene-by-cell-type proportions for each subtype. Paired Wilcoxon-ranked tests for same cell types in the two subtypes reveals significant differences in gene-cell-type probabilities for all cell types (p-value < 2.2x10^-16^ for all comparisons). c) Classification of subtypes based on phenotype probabilities. Visualization of phenotype probabilities inferred for ductal and lobular subtypes for the held-out set. The plot is ordered based on decreasing values of inferred phenotype probabilities for ductal subtype (rows), with the phenotype labels indicated on the left color bar. d) Comparison with DE genes from DESeq2. Comparison of gene-topic scores inferred by our approach for DE genes detected by DESeq2 in a Lobular versus Ductal comparison. On the left, the log2 fold change in expression of upregulated genes (top half) and downregulated genes (bottom half) identified by DESeq2 are arranged in decreasing order of the adjusted p-values. On the right, a heatmap of differences in gene-by-topic proportions for the same cell type, inferred by our method for a lobular versus ductal comparison, is shown for corresponding genes. e) DE genes identified by our method. Heatmap of 284 statistically significant DE genes (p-value < 0.05) identified by our method (rows), depicting difference in gene-topic-scores for the same cell types. The top cluster shows the upregulated genes in ductal subtype while the bottom cluster shows the downregulated genes. DE genes corresponding to those identified by DESeq2 are indicated in the left color bar. f) ORA analysis of differences in gene-topic proportions. Statistically-significantly enriched HALLMARK pathways identified in an ORA analysis based on differences in gene-topic-scores per cell type for ductal versus lobular comparison are indicated, for both upregulated and downregulated genes. Their significance level is indicated by the - log (adjusted p-value) on the y-axis, for the different cell types on the x-axis.

**Figure S30:**
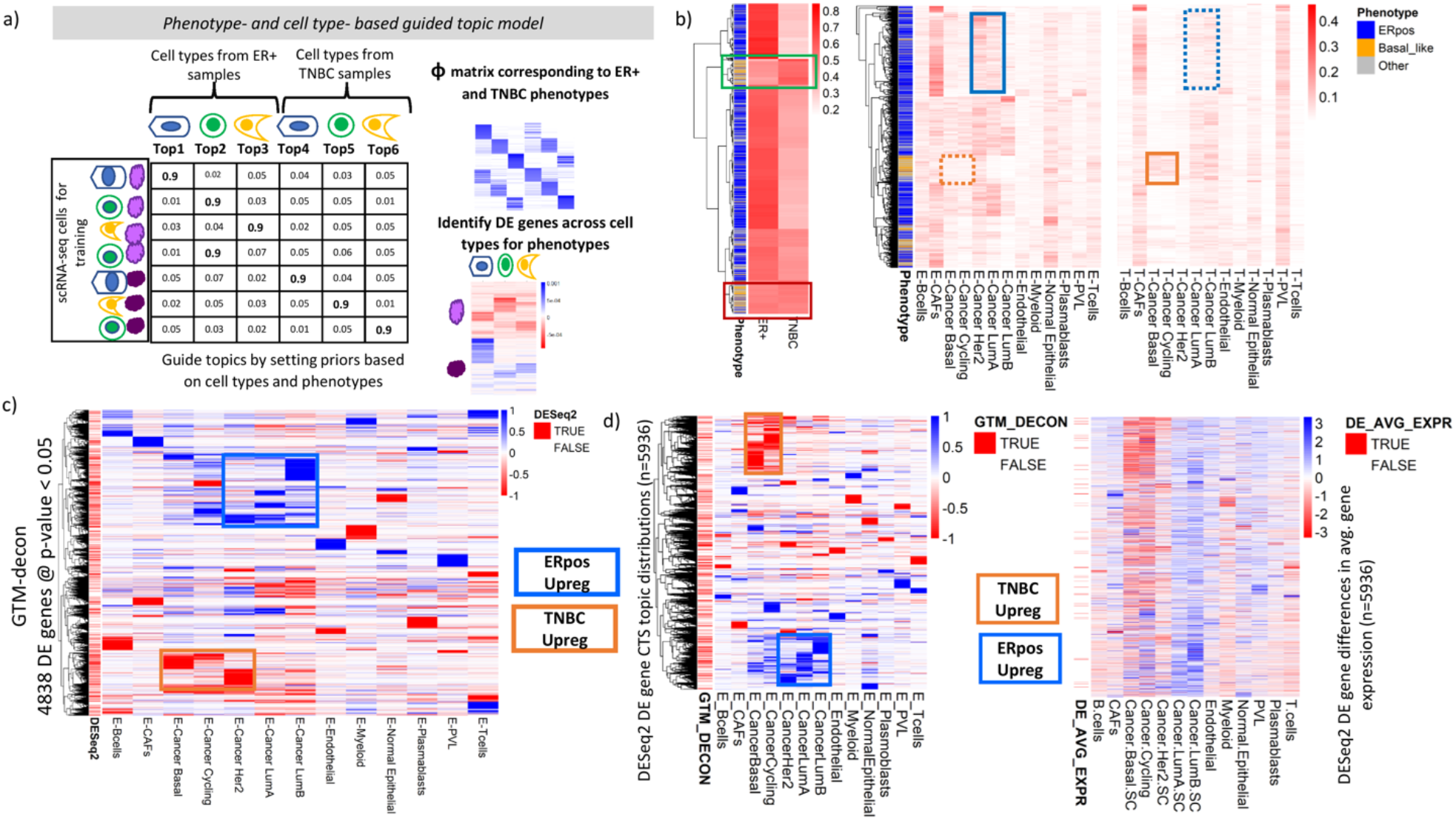
Phenotype-CTS guided topic inference of ER+ and TNBC breast cancer scRNA-seq data. **a)** Briefly, we inferred the phenotype-CTS topics by assigning one topic to each combination of cell type and phenotype. During GTM-decon training, the topic designated for the observed cell type and phenotype is endowed with a topic prior value of 0.9, and the rest of the topics were set at a random value between 0.01 and 0.1. As a result, these priors guide the topic model to infer phenotype-CTS topic distributions. All the genes in the experiment were used as features. **b)** Phenotype and phenotype-CTS deconvolution of TCGA-BRCA samples (n=1212). In both heatmaps, each row represents a bulk sample. For the left heatmap, the columns indicate the phenotype-specific topics, and for the right heatmap the columns indicate phenotype-CTS topics, where the prefixes ‘E’ and ‘T’ before the cell type names indicate ER+ and TNBC cancer subtype, respectively. Enrichment (solid rectangle) and depletion (dash rectangle) for known cell types for the cancer subtypes were indicated using blue rectangles (for ER+) and orange rectangles (for TNBC), respectively. **c)** Genes-by-CTS topic proportions for the 4838 statistically significant DE genes at the empirical p-value < 0.05 based on 100,000 permutation tests, out of which 1687 genes overlap with the 5936 DE genes identified from bulk TCGA-BRCA via DESeq2. The rows correspond to DE genes, and the columns to the change of CTS topics for ER+ with respect to TNBC (therefore indicated as columns with prefix ‘E’). The differences for TNBC minus ER+ comparison are the mirror image of this and hence not shown. **d)** Differential signals of the TCGA-BRCA DE genes from the DESeq2 analysis of the TCGA-BRCA data. We visualized the differential signals of the DE genes in two ways: (i) Differential CTS topic probabilities. The left heatmap shows the change of the genes-by-CTS topic probabilities in ER+ w.r.t. Basal-like subtype from the single-cell breast cancer transcriptomes for the 5936 DE genes detected by DESeq2 for ER+ cancer subtype w.r.t. Basal-like cancer subtype from the bulk TCGA-BRCA transcriptomes. Among these 5936 DE genes, 1687 genes were among the 4838 DE genes detected by 100,000 permutation tests from the scRNA-seq breast cancer data based on the differences in CTS-topic probabilities between the ER+ and TNBC phenotypes for each cell type. These genes are shown by the row annotations on the left side of the heatmap. (ii) Differential average gene expression for ER+ subtype w.r.t. TNBC subtype per cell type. The right heatmap shows the average gene expression differences between the ER+ subtype and TNBC subtype calculated directly from the observed single-cell data. For the ease of comparison, the rows were ordered according to the clustering pattern from the left heatmap. Among the 5936 DE genes from DESeq2, 592 genes were among the 1892 DE genes detected by 100,000 permutation tests based on the differences in average gene expression between the ER+ and TNBC phenotypes for each cell type. These genes are shown by the row annotations on the left side of the heatmap.

**Figure S31.**
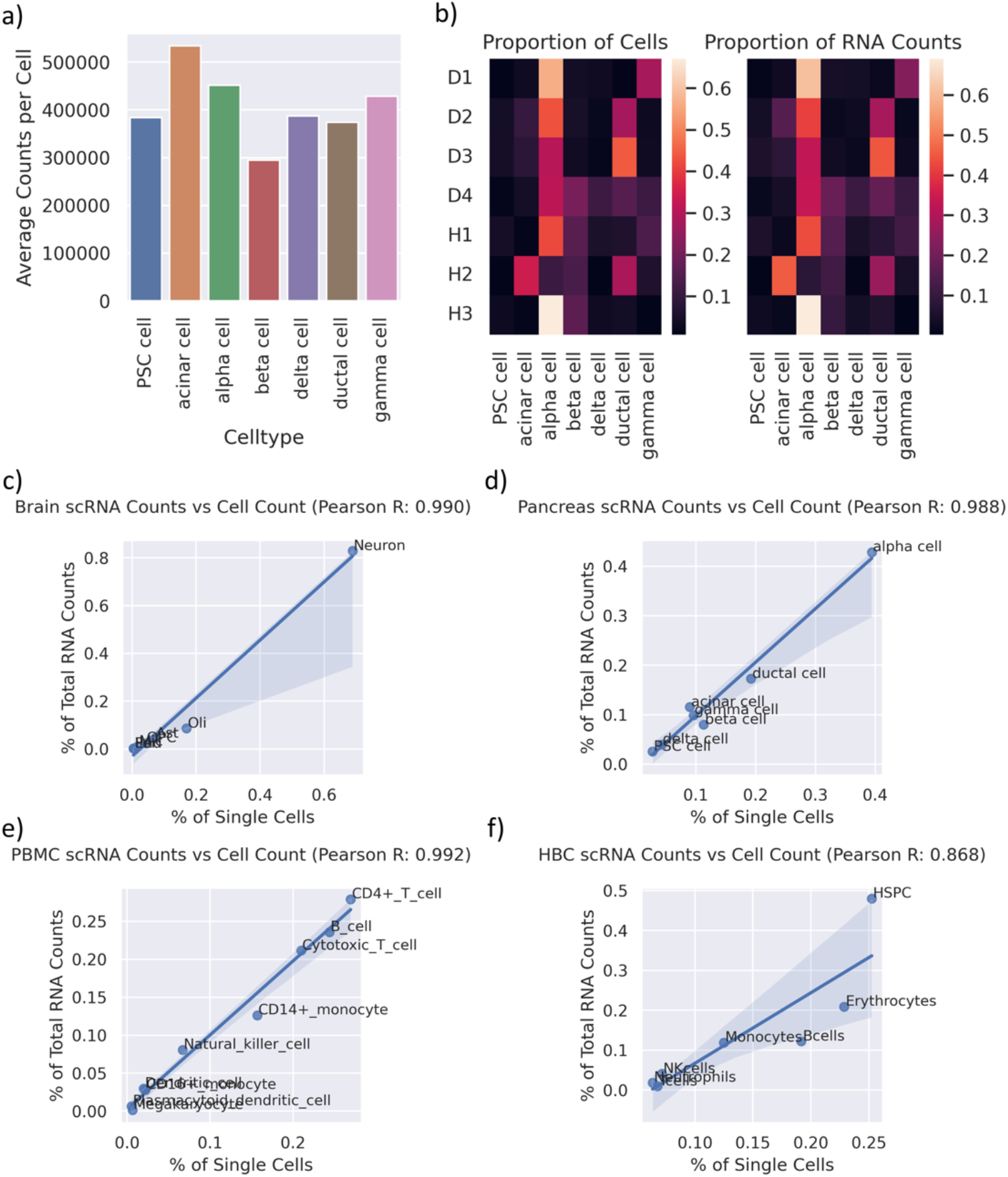
Correlation between Cell Fraction and Cell-type Specific RNA Fraction. a) Average number of RNA read counts per cell-type in single-cell Pancreas Data (cell-types with >40 cells available). b) Heatmaps comparing cell fraction and RNA fraction per cell-type for Pancreas single-cell Data across 7 patients. c-f) Scatterplots comparing the total percentage of RNA read counts for a given cell-type vs the fraction of single cells of that given cell type for the Prefrontal Cortex, Pancreas, PBMC, and HBC single-cell references respectively.

